# MYC Binding Near Transcriptional End Sites Regulates Basal Gene Expression, Read-Through Transcription and Intragenic Contacts

**DOI:** 10.1101/2024.07.11.603118

**Authors:** Huabo Wang, Bingwei Ma, Taylor Stevens, Jessica Knapp, Jie Lu, Edward V. Prochownik

## Abstract

The MYC oncoprotein regulates numerous genes involved in cellular processes such as cell cycle and mitochondrial and ribosomal structure and function. This requires heterodimerization with its partner, MAX, and binding to specific promoter and enhancer elements. Here, we show that MYC and MAX also bind near transcriptional end sites (TESs) of over one-sixth of all annotated genes. These interactions are dose-dependent, evolutionarily conserved, stabilize the normally short-lived MYC protein and regulate expression both in concert with and independent of MYC’s binding elsewhere. MYC’s TES binding occurs in association with other transcription factors, alters the chromatin landscape, increases nuclease susceptibility and can alter transcriptional read-through, particularly in response to certain stresses. MYC-bound TESs can directly contact promoters and may fine-tune gene expression in response to both physiologic and pathologic stimuli. Collectively, these findings support a previously unrecognized role for MYC in regulating transcription and its read-through via direct intragenic contacts between TESs and promoters.

## Introduction

The c-MYC (MYC) bHLH-ZIP oncoprotein transcription factor (TF) and its obligate bHLH-ZIP heterodimerization partner MAX supervise the transcription of hundreds-thousands of genes involved in essential processes such as cell cycle progression, mitochondrial and ribosomal structure and function and translation.^1–4^ Sequence-specific DNA binding by MYC-MAX heterodimers classically occurs at canonical “E boxes” that reside in proximal promoters ^4,5^.This prompts the recruitment of histone acetylases, which modify and relax chromatin so as to facilitate subsequent entry by RNA polymerase II (RNAPII) and other transcriptional co-factors. Together, these drive mRNA capping and overcome blocks to elongation that otherwise suppress the initiation and maintenance of robust levels of transcription. ^6,7^. MAX can also associate with 6 additional bHLH-ZIP TFs (MXD[1-4], MNT and MGA), collectively known as the “MXD” family. ^4,8,9^ Often expressed in ways that are tissue-specific and developmentally-dependent, MAX-MXD heterodimers compete for many of the sites occupied by MYC-MAX.^4,9^ In doing so, they recruit histone de-acetylases, which revert the adjacent chromatin to its original compacted and transcriptionally suppressed state. ^4,8^ Together, the above factors comprise the so-called “MYC Network”, which exerts sensitive, flexible and integrative regulation and balance of MYC target gene expression in response to what are often rapidly changing and conflicting environmental and proliferative cues.^4,9^ The importance of maintaining strict control among MYC Network member level and function is underscored by the embryonic lethality of body-wide *Myc* or *Max* gene knockout (KO) in mice, by the accelerated aging phenotype that accompanies post-natal *Myc* KO, by the association of *MYC* over-expression with numerous naturally-occurring human cancers and by the ability of MYC over-expression to drive experimental tumors in mice. ^6,10–13^

MYC also negatively regulates gene expression. This is mediated indirectly by MYC-MAX heterodimers interacting with and suppressing positively-acting TFs such as MIZ1, SP1 and SP3, which normally bind sequence-specific Initiator elements and GC-rich SP1 sites, respectively, and, like E boxes, tend to lie in close proximity to transcriptional start sites (TSSs).14,15

Previous studies of MYC-mediated target gene regulation have focused on its associations with the above-mentioned promoter-proximal regions or enhancer elements. The locations of the latter sites cannot be predicted *a priori* and, while often located many kilobases upstream or downstream of the gene’s coding region, may also be embedded within the gene body itself ^5,16,17^ Here, however, we describe and characterize evolutionarily conserved MYC and/or MAX. binding located in the vicinity of transcriptional end sites (TESs) of nearly one-sixth of all annotated genes. Like TSS-associated 5’-end binding, MYC/MAX binding near 3’-end TESs regulates target gene expression in ways that reflect the identities of the bound factors, their cooperation with MYC/MAX binding at other sites and tissue-specific preferences. TES-associated MYC/MAX binding also tends to occur in genes with functions that differ from those lacking such binding. Finally, it contributes to both basal and stress-induced transcriptional read-through and the formation of intragenic contacts both with promoters and more distally located enhancers. Collectively, these studies reveal previously unrecognized forms of gene regulation mediated by unconventional but common MYC binding near TESs.

## Results

### MYC and/or MAX frequently bind in proximity to TESs

Using genome-wide ChIPseq data sets compiled from 5 human and 2 murine cell lines from the ENCODE and GEO databases, we performed an unbiased search for MYC and MAX binding sites at the 5’- and 3’-ends of all annotated genes (Figure 1A and Table S1).^18–20^ To maximize the likelihood that binding locations were assigned as unambiguously as possible and to functionally meaningful sites, we included only those with peaks residing within +/- 2.5 kb of TSSs or TESs. In doing so, we excluded peaks that could potentially be assigned to either 3’- or 5’-ends in closely adjacent genes orientated in a tail-to-head configuration. Binding sites for closely neighboring genes in head-to-head or tail-to-tail configurations were assigned to both genes. To further simplify our classification, the minority populations of genes containing multiple binding sites within the regions of interest were included in the same categories as those with single sites (Figure S1).

**Figure 1.**
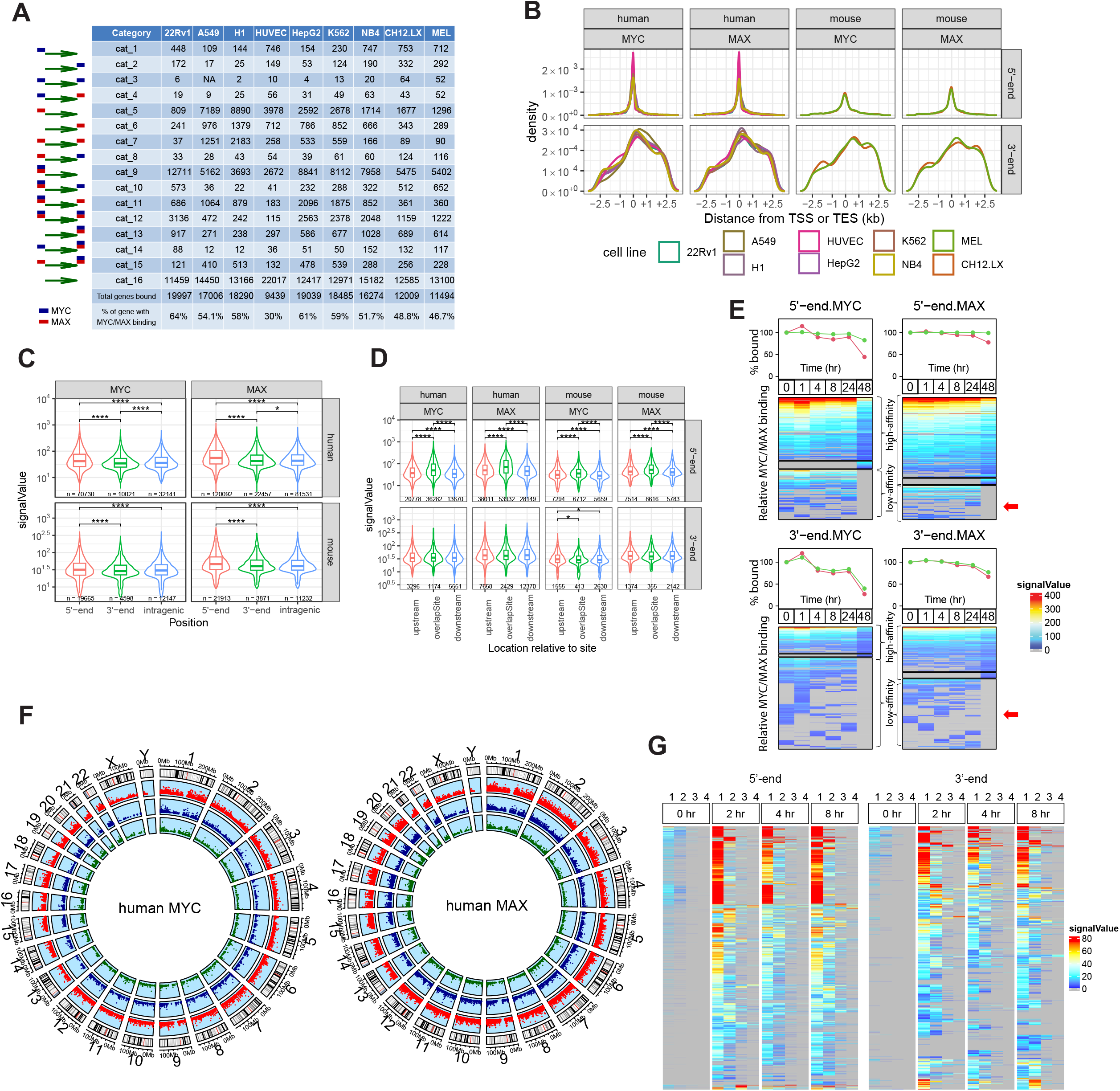
MYC and/or MAX binding site distribution and affinities in the vicinity of TSSs and TESs of human and murine genes. (A). Frequency of MYC and MAX binding sites in proximity to TSSs and TESs of all annotated genes in the indicated human and murine cell lines. All possible configurations of MYC(blue) and/or MAX(red) binding are depicted at the left and are arbitrarily designated as categories (cat.) 1-15. Each column indicates the number of genes from each category that were identified in the referenced cell line. To simplify classification, no distinctions were made between genes with single or multiple binding sites (Figure S1). (B). Distribution of MYC and MAX binding sites relative to TSSs and TESs, which are defined as residing at position “0” within their respective genes. Some genes are repeated to allow for assignments to multiple TSSs and/or TESs (Figure S1). Binding site distribution patterns in individual cell lines are shown in Figure S2. (C). Mean binding signals +/- 1 S.E. for MYC and MAX around TSSs, TESs and intragenic regions for the genes depicted in A. Numbers below each violin plot indicate the actual number of binding peaks that were counted. *: P<0.05; ****: P <0.0001. (D). Mean binding signals +/- 1 S.E. for MYC and MAX whose peaks reside up to 2.5 kb upstream or downstream of TSSs and TESs. (E). Binding near TSSs and TESs stabilizes MYC. Upper portion of panels: 22Rv1 prostate cancer cells were exposed to 10 µM of the MYC inhibitor Myci975 for the indicated times.^19,62^ MYC and MAX ChIPseq were then performed and their binding signals around TSSs and TESs of all target genes were quantified. Green circles indicate percent of MYC or MAX remaining bound. Red circles indicate binding signal intensity at these sites relative to that at t=0. Lower portion of panels: heat maps indicating the amounts of MYC or MAX remaining bound to individual genes with high- and low-affinity MYC and/or MAX binding. Immuno-blotting of whole cell lysates from the cells used in these ChIPseq studies had previously demonstrated near-total depletion of MYC protein by 48 hr.^19^ Red arrows indicate groups of genes with low-affinity sites that are selectively depleted of Myc binding and completely so by 48 hr. (F). Circos plots indicating the chromosomal locations of annotated human genes that bind MYC and/or MAX only at their 5’-ends (red dots), only at their 3’-ends (green dots) or at both ends (blue dots). The results were complied from all 7 of the human cell lines shown in panel A and Table S1.and Figures S3 and S4 show these sites in individual human cell lines. Figures S5 and S6 show Circos plots for individual murine cell lines. (G). Time-dependent binding of MYC and MAX at multiple TSS- and TES-associated sites of genes in murine B lymphocytes following exposure to LPS at “0 hr”. All genes identified as containing more than one MYC binding site in the vicinity of TSSs and TESs at any time during LPS induction are depicted. The binding sites were identified at each point during this exposure and the binding signals (affinities) quantified as explained in E. Numbers at the top of each panel indicate the individual MYC binding sites that were identified as being closest to (1) and furthest from (4) the TSSs or TESs of their respective genes.

The number of genes represented by each category varied up to 2.1-fold across cell lines (ex. 22Rv1 vs. HUVEC). The near-identical and tightly clustered 5’-end binding patterns among all cell lines showed that 64.4-83.5% of MYC and MAX binding sites resided within +/-500 bp of TSSs and 79.0-91.5% resided within +/-1.0 kb (Figure 1B and Figure S2). 3’-end binding patterns of MYC and MAX were also quite similar among cell lines, although more broadly distributed, with 49.1-56.9% of sites located within +/- 1.0 kb of TESs. Binding of MYC and/or MAX was detected at the 3’-ends of over one-sixth (17.5%) of all annotated genes. When considering both TSS- and TES-proximal binding sites, some configurations were more common than others although this also varied among cell types. For example, “category 9” binding (MYC+MAX binding at 5’-ends) was the most common configuration in 6 of the 9 cell lines and accounted for 20.2-63.6% of all bound annotated genes (Figure 1A). The least common categories (3 and 4), representing 5’-MYC/3’-MYC and 5’-MYC/3’-MAX binding, collectively comprised no more than 0.50% of all bound genes in any cell type. Other categories showed large differences in their representation among the cell lines. For example, category 7 (5’-MAX/3’-MAX) varied over >60 fold range as a percentage of all bound genes and category 10 (5’-MYC+MAX/3’-MAX) varied by nearly 50-fold. At both 5’- and 3’-ends, the number of individual MYC and/or MAX binding peaks for the vast majority of genes in all cell lines (as determined by the presence of distinct individual ChIP-seq peaks) was no greater than 3-4, with multiple sites being more frequent at 5’-ends (Figure S1). Collectively, these results indicated that different cell lines displayed distinct MYC+MAX gene binding profiles centered around TSSs and TESs. They also suggested that these patterns represent only snapshots of single conditions, points in time, cell types and states of differentiation that almost certainly change in response to different intracellular and environmental cues, including the levels of expression of various MYC Network members *(vide infra)*.

Peak signal intensities of MYC and MAX binding around TSSs and TESs from the above cell lines were used to approximate each factor’s accessibility to DNA and its binding affinities. On average, these signals were stronger at 5’-ends although considerable overlap was documented (Figure 1C). They were also stronger for peaks in closest proximity to TSSs (Figure 1D). In contrast, MYC binding at 3’-ends was equally strong irrespective of where it occurred relative to TESs whereas MAX binding was modestly stronger at upstream sites. By comparison, Myc and Max binding affinities at TESs in both human and mouse genes were equal to or only modestly lower than the affinities at intragenic sites (>2.5 kb downstream of TSSs and >2.5 kb upstream of TESs) that likely reflected enhancer-specific binding.

The above binding signals were also used to measure MYC and MAX displacement from their 5’- and 3’-end sites in 22Rv1 prostate cancer cells following exposure to the potent and specific small molecule MYC inhibitor MYCi975 (Figure 1E). ^19^ This led to 4 observations. First, and consistent with the results shown in Figure 1C, the mean binding affinities for MYC and MAX around TESs in untreated cells were, on average, weaker than those around TSSs as estimated both by the strength of their basal binding signals and the rates at which they were diminished in response to MYCi975. Second, low-affinity MYC and MAX binding, particularly at 3’-ends, tended to be depleted within 24-48 hr of MYCi975 exposure (Figure 1E). Third, despite a >90% reduction in total MYC levels as determined by whole cell immuno-blotting, ^19^ significant amounts of MYC and MAX remained bound to higher affinity sites at both ends for >48 hr after exposure to MYCi975 (82.5% vs. 39.8%, respectively, for MYC [P=1.1x10^-61^] and 99.0% vs. 76.9%, respectively, for MAX [P=1.3x10^-12^]). Finally, MAX’s displacement from both 5’- and 3’- ends at 48 hr was less than that of MYC’s (P=5.5x10^-21^ and P= 6.5x10^-35^, respectively). This could reflect MAX’s predilection to re-associate with MXD members and retain its binding as MYC levels declined following exposure to Myci975 as well as its intrinsically longer half-life. ^4,9,19^ The unexpected persistence of significant MYC binding indicated that its association with DNA markedly extends its half-life well beyond the standardly accepted 15-30 minutes. ^21^

The loss of MYC and MAX binding around high-affinity TSSs and TESs closely paralleled the MYCi975 exposure time (Figure 1E). In contrast, low-affinity binding site displacement kinetics were, in some cases, more complex. For example, binding was initially lost at some sites, only to reappear at later times and then disappear again by 24-48 hr. In other cases, no binding was seen initially but paradoxically appeared after a period of exposure to MYCi975 only to subsequently vanish. These findings suggested that MYCi975 treatment initially altered the local chromatin environment in ways that allowed some binding sites, previously unoccupied by MYC and/or MAX to become permissible for transient binding, even though overall MYC levels were in decline.^19^

Although the combined data from the above human cell lines revealed no obvious preferences in the genomic locations of MYC or MAX binding at TSS- or TES-associated sites (Figure 1F), some differences were noted among individual cell lines. For example, less frequent binding of MYC to TES-associated sites was noticeable on chromosomes 4 and 13 in A549 cells and on chromosome 4 in HUVECs and H1 cells (Figure S3). Similarly, less frequent binding of MYC to both 5’ and 3’ sites was noted on chromosome 13 in A549 cells and on chromosomes 8 and 13- 15 in H1 cells. This type of segregation was less notable for MAX binding at TES-associated sites, although a relative paucity was seen in A549 cells (chromosome 13) and HUVECs (chromosomes 4 and 13). A relative lack of genes binding MAX around both TSSs and TESs was also seen on chromosome 15 in HUVECs (Figure S4). These findings support the idea that some of the gene categories shown in Figure 1A are distributed unequally across the genome and can sometimes be bound by MYC and/or MAX in tissue-specific ways. Such unequal distribution was less evident across the murine genome although it may have been due to a lower number of evaluable cell lines, particularly for MAX (Figures S5 and S6).

Finally, we assessed ChIPseq results from *D. melanogaster* for dMYC and dMAX binding sites.^18^ Because shorter regions of non-coding inter-genic DNA exist in the fly genome, more stringent criteria were used for the search. We therefore included only those genes that bound these factors within +/- 1.0 kb of TSSs or TESs. As was true for humans and mice, many genes bound dMYC and/or dMAX near both TSSs and TESs (Figure S7). Their more homogeneous chromosomal binding may have reflected the significantly smaller genome of the fly as well as the extensive larval tissue heterogeneity. At the very least, our findings provide support for the notion that both MYC- and MAX-associated TES binding have remained highly conserved over >700 million years of metazoan evolution.^22^

As mentioned above, some genes contain multiple TSS- and/or TES-associated MYC binding sites (Figure S1). To determine the order in which these sites bind Myc and their relative affinities, we again used ChIPseq data to catalog the kinetics of newly synthesized MYC binding following lipopolysaccharide (LPS)-mediated activation of quiescent murine B cells. ^18^ Prior to LPS induction, and despite the extremely low levels of MYC protein in these cells, small numbers of presumably high-affinity MYC binding peaks were nonetheless detected at both 5’- and 3’-ends and tended to be those in closest proximity to TSSs or TESs (Figure 1G). Within 2 hr of LPS induction, large increases in the intensities of these signals were observed as were new signals at more distal sites that were either of lower affinity or had been previously inaccessible. Thus, in response to MYC induction, the kinetics of binding site occupancy, the number of sites bound and the order in which this progressed were very similar at 5’- and 3’- ends. They largely reflected increasing Myc levels and the distance of binding sites from TSSs and TESs, with the highest affinity binding occurring almost exclusively at sites closest to these landmarks.

### MYC binding around TESs impacts gene expression and transcriptional read-through

RNAseq results from the ENCODE and GEO databases^18^ allowed transcript expression comparisons for the 16 gene categories depicted in Figure 1A. As expected, genes comprising categories 10, 11 and 12, all of which bound MYC+MAX at TSS-proximal sites, were expressed at higher levels than those lacking this binding (categories 6,13 and 16) (Figure 2A and 2B). However, within the former categories were also subsets of genes that were expressed at low levels in cell type-specific ways. Conversely, the latter categories sometimes contained highly expressed gene subsets, particularly in HUVECs and MEL cells. Evidence that gene categories binding MYC and/or MAX only near TESs also differed in their expression relative to categories that did not (i.e. categories 2, 6 and 13 versus category 16) was seen in several cell lines (A549, H1, NB4, CH12.LX and MEL). These findings indicated that the binding of MYC and MAX at the 3’-ends of genes could impart variable degrees of generally positive control in a cell type-specific manner when assessed agnostically across all expressed genes. The remarkable intercellular consistency of expression patterns across all 16 gene categories, despite differences in their gene content, supported the idea that the presence and location of MYC and/or MAX binding sites at both 5’- and 3’-ends is a general and perhaps more fundamental aspect of gene expression that operates independently of tissue-specific factors or specific gene identities.

**Figure 2.**
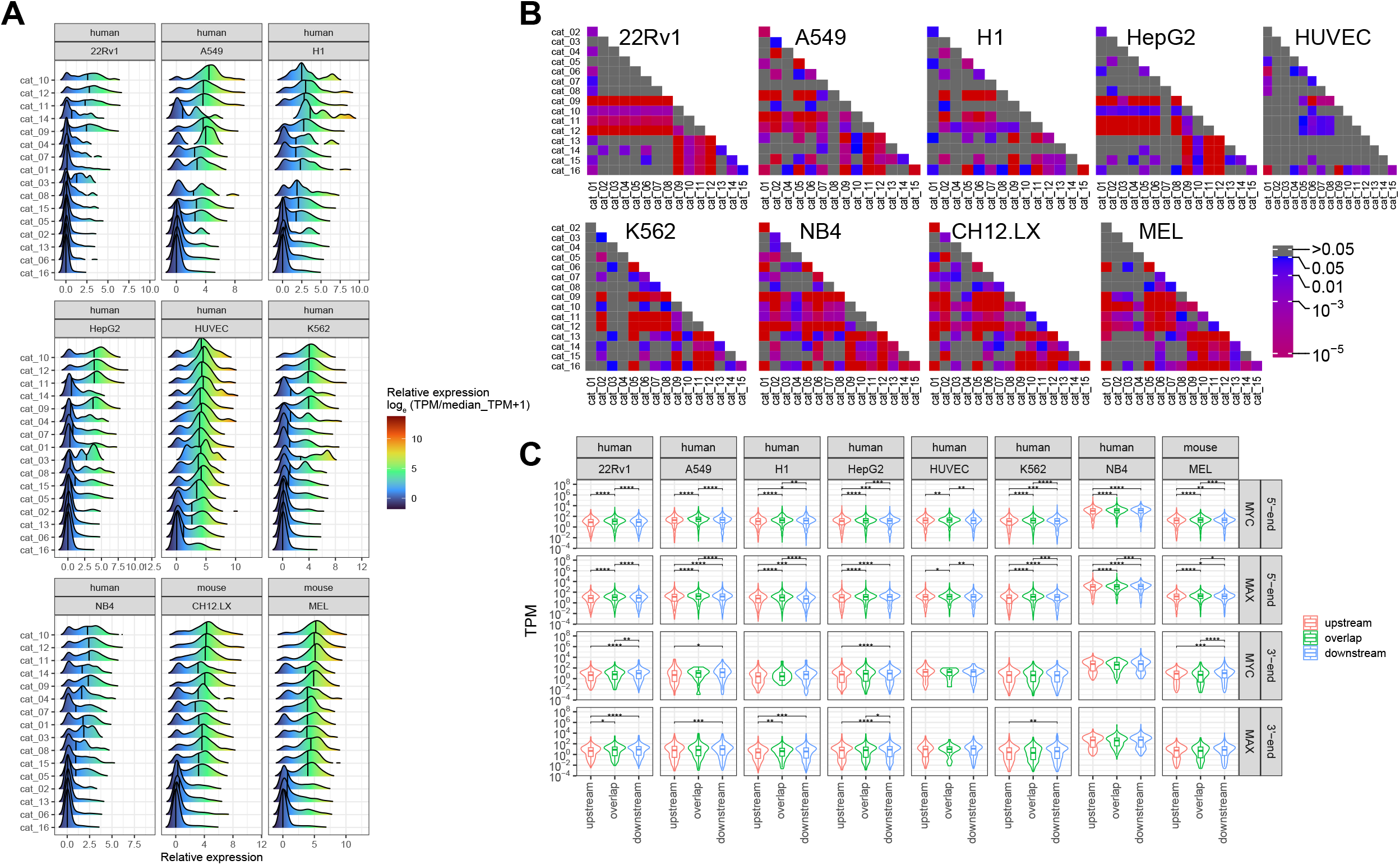
Gene expression is influenced by MYC and/or MAX binding to TESs of human genes. (A). Ridge plots of RNAseq results from all 16 categories of genes depicted in Figure 1A. Results are plotted to allow for both inter- and intra-cell line comparisons. Blank areas indicate categories for which too few genes were available to allow for meaningful comparisons. B). The results from A were compared for significant differences in mean gene expression levels among all genes within the indicated categories. (C). Gene expression levels among cell lines correlate with the location of MYC and/or MAX binding around TSSs and TESs. Using the data shown in Figure 1 C&D and Figure S2, MYC and MAX binding peaks were assigned to sites that either overlapped TSSs or TESs or resided upstream or downstream of them. The mean transcript levels were then plotted for each group among the cell lines for which data was available. **: P <0.01; ***: P <0.001

Related to the way that MYC and MAX binding signal strengths reflected the distance of their peaks from TSSs and TESs (Figure 1C,1D and 1G) so too was the expression of the genes to which they bound. Those binding MYC and MAX at sites closest to or overlapping TSSs tended to be more highly expressed than those in which binding was more remote (Figure 2C). In contrast, only modestly higher levels of expression were associated with MYC and MAX binding downstream of TESs.

Chip-seq and RNA-seq results from the LPS-treated murine B cells described above were examined to allow correlations to be made between MYC binding signals and gene expression when Myc was induced by a physiologic stimulus. Despite the high-affinity binding of MYC to the previously mentioned minority of target genes prior to LPS exposure (Figure 1G), very little effect on their expression was seen relative to that in control *Myc-/-* B cells (Figure 3A-3C, red arrows). For these genes, as well as for those without any initial evidence of initial MYC binding, LPS treatment led to rapid MYC association that was largely terminated within 2 hr at both high- and low-affinity sites, although examples of delayed binding were observed as well (Figure 3A- 3C, blue and green arrows, respectively). Gene expression changes typically lagged DNA binding by 2-6 hr although exceptions were again noted. Seen in all 3 groups, but particularly prominent among genes with TES-associated MYC binding only, were many examples in which MYC binding post LPS induction was transient despite more persistent changes in gene expression (Figure 3B magenta arrows). Comparisons of signal intensities again indicated, as noted previously (Figure 1C), that MYC binding at 3’-ends was of somewhat lower average affinity. This was best observed in genes with MYC binding at both ends where high-affinity binding prior to LPS induction was overwhelmingly confined to 5’-ends (Figure 3C). Following LPS induction, however, a substantial number of these genes showed roughly equivalent levels of 5’- and 3’-end MYC binding and some showed more robust binding at 3’-ends (Figure 3C, gray and orange arrows, respectively).

**Figure 3.**
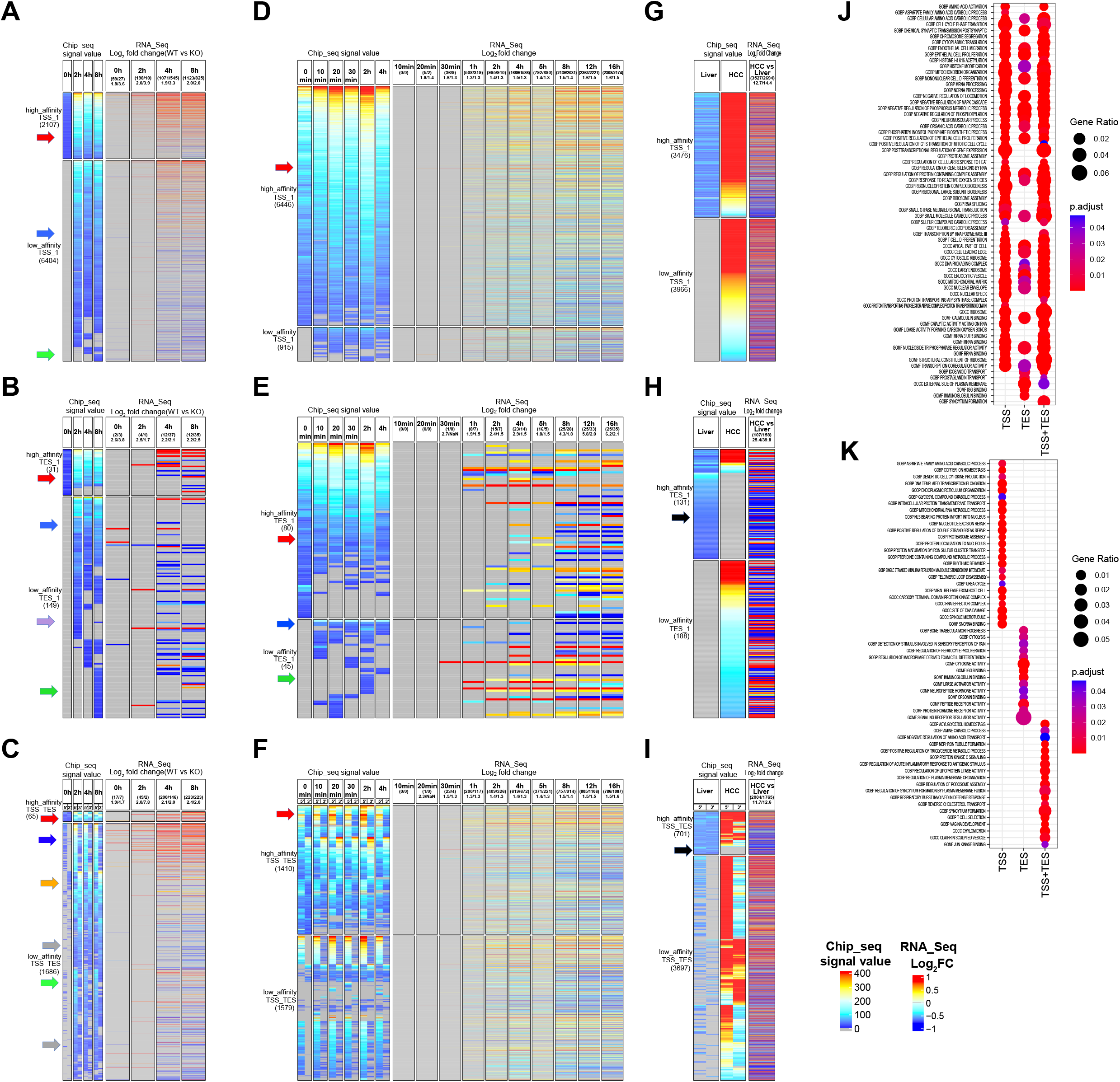
MYC binding near TSSs and/or TESs correlates with differences in gene expression and function. (A-C). Quiescent wild-type (WT) primary murine B lymphocytes treated with LPS ^63^. Myc ChIPseq and RNAseq were performed at the indicated times after LPS exposure and compared to the results obtained from LPS-treated *Myc*-/- B lymphocytes. Panel A: Chipseq and RNAseq results for genes binding MYC around TSSs only. Panel B: Chipseq and RNAseq data for genes binding MYC around TESs ‘only. Panel C: Chipseq and RNAsea data for genes binding MYC at both TSS- and TES-associated sites. Red arrows: examples of gene groups with high-affinity MYC-binding sites; blue arrows: examples of low-affinity sites showing early occupancy by MYC in response to LPS; green arrows: examples of low affinity binding sites showing delayed occupancy by MYC; magenta arrows: examples of MYC binding sites transiently occupied by MYC; gray arrows: example of MYC binding sites of about equal affinities around TSS and TES sites; orange arrows: example of MYC more robust binding at TES sites; black arrow: example of high-affinity sites failed to maintain MYC binding. Numbers above the heat maps indicate the total number of gene differences between the indicated comparisons and the mean fold up- or down-regulation. (D-F). Logarithmically growing murine fibroblasts harboring a conditional Myc-estrogen receptor chimaera were treated with 4-hydroxytamoxifen to activate Myc for the indicated periods of time. MYC ChIPseq and RNAseq were then performed to allow correlations between binding and expression for the subsets of genes that bound MYC only in proximity to TSSs (D), TESs (E) or both sites (F). Arrows indicate similar gene sets described in panels A-C. (G-I). HCCs were generated in mice by the hepatocyte-specific and doxycycline-regulatable induction of a human *MYC* transgene.^12^ MYC ChIPseq and RNAseq were then performed to allow correlations between binding and expression for the subsets of genes that bound MYC only in proximity to TSSs (G), to TESs (H) or to both sites (I). Black arrow in I: examples of high- affinity MYC binding sites present only in normal liver. Other arrows indicate examples of genes whose behaviors are described in panels A-C. (J). Over-representation analysis from the MSigDB database’s C5 GO collection. Genes that bound MYC exclusively at or around TSSs, exclusively at or around TESs or at or around both TSSs and TESs were categorized into functional groups, the most significant of which are indicated. (K). Functional categories of genes uniquely associated with binding of MYC to the sites indicated in J.

MYC binding to TSS- and TES-associated sites under “pathological” conditions of MYC over- expression was examined in logarithmically growing immortalized murine fibroblasts following high-level induction of a *MYC*-estrogen receptor (*Myc*ER) fusion transgene. ^23^ Because these cells initially expressed substantial amounts of endogenous MYC, its binding to lower affinity target genes prior to MYC induction was more prominent and extensive than documented previously in quiescent B cells (Figure 3D-3F, red arrows). As was true for B cells, many genes were associated with low-affinity 3’-end sites that bound MYC at different rates (Figure 3 E, blue and green arrows).

Another example of target gene binding and gene expression changes in response to pathologic levels of MYC occurred in undifferentiated hepatocellular carcinomas (HCCs) induced by conditional *MYC* over-expression.^11,12^ Despite its exceedingly low level expression in livers, MYC binding was nonetheless detected at high-affinity 5’- and 3’-ends, whereas its binding to lower-affinity sites was noted only in response to its over-expression (Figure 3G-3I). A small subset of high-affinity sites also failed to maintain MYC binding despite the continued dysregulation of the associated target genes, most of which were down-regulated (Figure 3H and 3I black arrow).

MYC target genes disproportionately participate in the maintenance of energy production, the support of ribosomal structure and function, the regulation of the cell cycle and the response to and repair of genotoxic damage.^1,3,4,12,13,24^ A search of the C5 GO Collection from the MSigDB showed that genes binding MYC only at their 5’-ends, only at their 3’-ends or at both ends tended to share many of these and other functions (Figure 3J) However, certain non-canonical functions were uniquely associated with the latter two groups. For example, genes with MYC binding restricted to 3’-ends were more likely to be implicated in differentiation, proliferation and cytokine/immune responses whereas those with MYC binding at both ends were more likely to be involved in transport and membrane maintenance (Figure 3K).

MYC and MAX also disproportionately bound at or near the TESs of genes encoding long non-coding RNAs (lncRNAs) (Figure 4A and 4B). In all cell lines examined, a greater proportion of lncRNA genes bound MYC and MAX at 3’-ends than at 5’-ends (human: 19% vs. 14%, P= 5.40e-61; mouse: 7% vs. 4%, P= 1.62e-164). Overall binding at both 5’- and 3’-ends was higher in human cell lines than in mouse cell lines (5’ end 14% vs 4% P=2.7e-196; 3’ end 19% vs. 7%, P= 2.2e-96). In those cell lines for which RNA-seq data was available, expression patterns of lncRNAs also differed among several of the 16 binding categories (for example, see categories 10, 12 and 14 Figure 4C).

**Figure 4.**
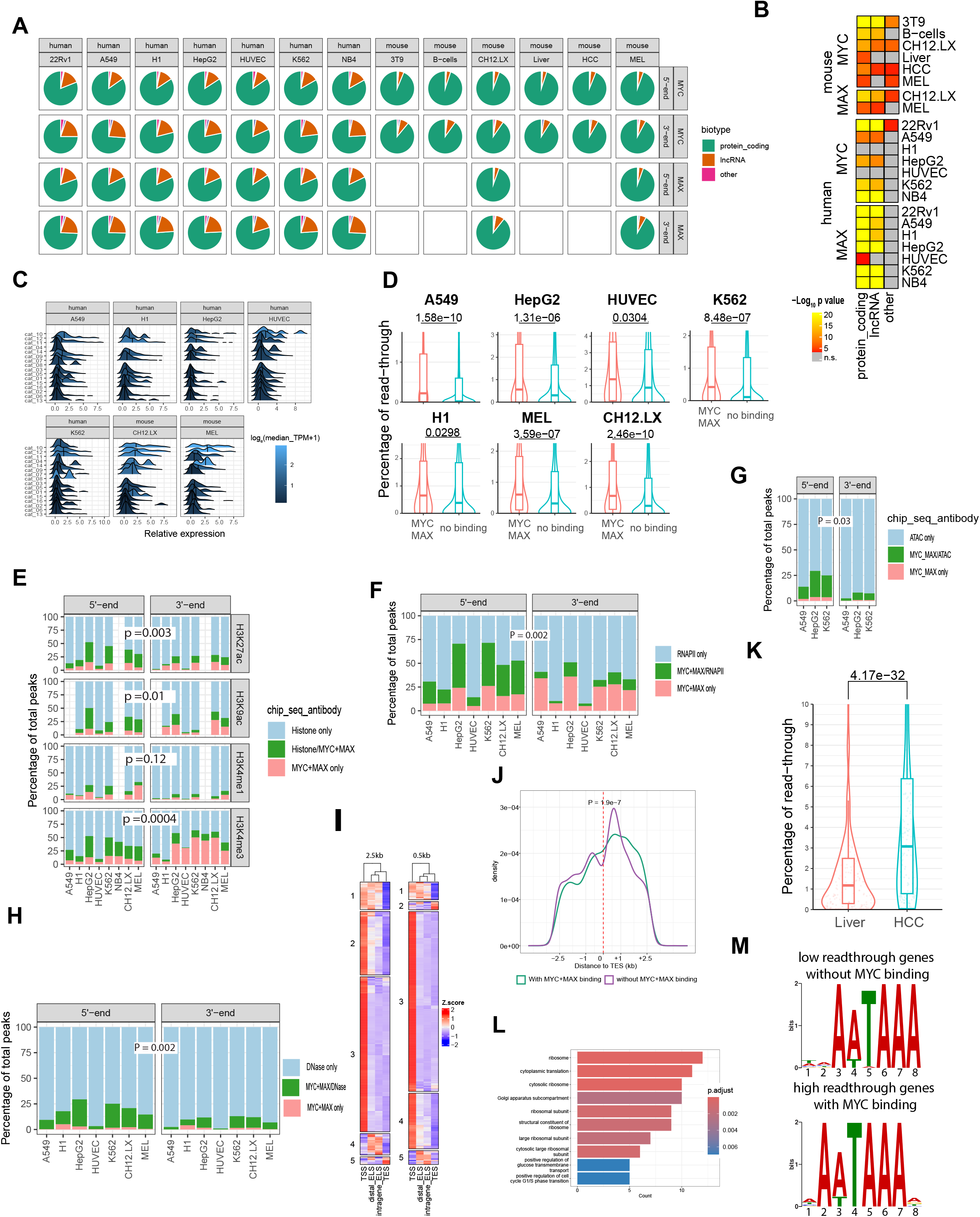
TES-associated MYC and MAX binding is enriched at genes encoding lncRNAs, participates in chromatin re-modeling and alters transcriptional read-through. (A). Fractional distribution of MYC and MAX binding around TSSs and TESs of genes encoding proteins, lncRNAs, miRNAs and other RNAs in the indicated cell lines. Blank boxes indicate data sets that were unavailable. (B). Comparisons of the fraction of genes from A binding MYC and MAX near TESs vs. TSSs in the indicated cell lines. (C). Ridge plots showing expression differences in lncRNAs for the 16 categories of MYC- and MAX-binding genes (Figure 1A). Blank panels indicate categories for which too few genes were available to allow meaningful distribution patterns to be shown. (D). MYC+MAX binding at and around TESs is associated with greater transcriptional read- through. For expressed genes with or without evidence for TES-associated MYC+MAX binding (Figure 1D), the ratio of transcripts containing sequences downstream of the TES to those originating from the last exon was determined in the indicated cell lines. (E). TSS- and TES-associated MYC+MAX binding site peaks coincide with those for the transcriptionally permissive histone modifications H3K27ac, H3K9ac, H3K4me1 and H3K4me3. ChIPseq data for annotated genes containing TSS- and TES-associated MYC+MAX binding sites within the domains demarcated by Figure 1B were examined for the indicated modified histones using ChIPseq results. ^18^ Each histogram shows the fraction of genes for which modified histone and/or MYC+MAX binding peaks were or were not coincident. (F). TSS- and TES-associated MYC+MAX binding site peak coincidence with those for RNAPII. ChIPseq studies were performed as described as in panel E. (G). ATACseq results showing susceptibilities of genes binding MYC+MAX in the vicinity of TSSs and TESs to Tn5 transposase cutting. (H). Susceptibilities to DNaseI digestion of genes binding MYC+MAX in the vicinity of TSSs and TESs. (I). Binding frequencies of 1210 TFs and co-factors (File S1) to TSSs, TESs and intragenic and distal enhancer-like sequences (ELS) associated with known MYC-binding human genes. The TSS- and TES-associated boundaries employed in the search were either the same as those used to define MYC and MAX binding (i.e. +/- 2.5 kb: left-most heatmap) (Figure 1B) or were narrowed to +/- 0.5 kb to minimize the inclusion of enhancer sequences (right-most heatmap). Numbers to the left of the panels (1-5) indicate subsets of factors whose binding patterns allowed the 4 different regions to be distinguished from one another. (J). Broader distribution of RNAPII around TESs that bind MYC and/or MAX than at those that do not. (K). Fractional DoG transcription of 190 genes expressed in livers and Myc-driven HCCs^11,12^ showing higher levels of TES-associated MYC+MAX binding in the latter. (L). Functional categories of the 190 genes shown in panel K as determined by IPA. (M). Consensus transcriptional termination/poly(A) sites in genes with the highest and lowest levels of DoG expression from K.

The confinement of MYC and MAX binding around TESs (Figure 1B) suggested a role in mRNA termination. The absence or mutation of cleavage/poly(A) signals, various cellular stresses and neoplastic transformation are among the factors associated with selective increases in transcriptional read-through efficiency of some genes at the expense of precise mRNA termination and polyadenylation. ^25–27^ Not uncommonly, these “downstream of gene” (DoG) transcripts extend 50-100 kb beyond canonical TESs. ^26,27^ We used ENCODE strand-specific RNA-seq data to obtain ratios of DoG transcript expression relative to that encoded by the closest upstream exon. In each cell line examined, MYC+MAX binding in the proximity of TESs was, on average, associated with higher fractional read through (Figure 4D). Thus, in addition to contributing to the physiologic and pathologic responses of certain transcripts, specific 3’-end binding by MYC and MAX is also associated with higher average fractional levels of basal transcriptional read-through.

TES-associated MYC+MAX binding sites, particularly those showing evidence of DoG transcription, might be predicted to possess a more relaxed and transcriptionally conducive local chromatin environment resembling that associated with promoters and enhancers. ^28–33^ Indeed, ChIP-seq results for acetylated and methylated histones H3K27-ac, H3K9-ac, H3K4- me1 and H3K4-me3 in the available cell lines from Table S1 showed that the binding signals for these transcriptionally permissive modifications around TSSs and TESs commonly overlapped those for MYC+MAX (Figure 4E). These results indicated that TES-associated MYC+MAX binding sites coincide with more extensive regions of transcriptionally permissive histone modifications and read through. This was further supported by demonstrating that many of the above 3’-end MYC+MAX binding sites also bound RNAPII and that the surrounding regions were susceptible to both ATAC seq and DNaseI digestion (Figure 4F-4H).

The above-described epigenetic modifications of the chromatin landscape recalled classical enhancer elements, which, like promoters, are also marked by numerous TFs, including MYC and RNAPII. ^28,29,31–34^ We therefore utilized the ReMap2022 to generate binding profiles for 1210 TFs and transcriptional co-factors that mapped to MYC-bound TESs, TSSs and both distal and intragenic enhancers (File S1). These were compiled from 737 human cell lines and tissues and 8103 Chip-seq data sets containing 182 million binding regions. Over a range of +/-2.5 kb, each of the above 4 Myc-bound regions showed distinct binding patterns for these factors that could be roughly categorized into at least 5 groups (Figure 4I, leftmost panel). Intragenic and distal enhancers, the most closely related with regard to their binding patterns, could barely be distinguished whereas TSSs and TESs could be distinguished by groups 1,2,3 and 5. Although TES binding profiles were more closely related overall to those of enhancers, they could nonetheless be readily identified by groups 1, 2, 4 and 5. These differences persisted when coverage was more narrowly restricted to include only factor binding footprints within +/-0.5 kb of TES although the number of factors belonging to each of the groups was altered (Figure 4I: rightmost panel). Thus, the local environments of MYC-associated TESs and enhancers, as defined by the binding of TFs and co-factors, were distinct from those of TESs. Finally, the distribution of RNAPII around MYC and/or MAX-associated sites was significantly broader than it was around sites unassociated with MYC and/or MAX binding (Figure 4J).

DoG transcription in tumors is often generated by stresses arising from nutrient deficiencies, redox imbalance, hypoxia and the de-regulated expression of oncogenes, including MYC.^26,35^. We therefore asked whether DoG transcription in Myc-induced HCCs ^11,12^ changed relative to that of normal livers due to more MYC binding near TESs. From 11 normal livers and 15 HCCs from the GEO database, we identified 190 genes with significantly higher levels of TES-associated Myc binding in the latter (Table S2). Though not necessarily expressed at higher levels in HCCs, these genes did demonstrate nearly 3-fold higher mean fractional levels of DoG-transcription (Figure 4K). Indeed, this was similar to that previously reported in stressed fibroblasts despite the latter results having been based upon the sequencing of nuclear RNA, which tends to be enriched for DoGs transcripts ^26^. These 190 transcripts comprised 34% of all DoG differences between livers and HCCs (total 590) indicating that factors other than MYC and MAX likely contribute to read-through transcription and/or that transient MYC binding can have long-term effects on transcription (Figures 3A-3F and 4D-4H).^26^ Unlike the >4800 DoG transcripts previously identified in stressed NIH3T3 cells, which reportedly did not comprise distinct functional categories,^26^ the above 190 transcripts were enriched for those encoding ribosomal proteins and translation factors, which represent major categories of Myc-regulated target genes (Figure 4 L). ^3,4,13,36^ 35% of them were also previously shown to have higher levels of DoG expression in NIH3T3 murine fibroblasts subjected to oxidative stress. ^26^

One proposed explanation for why certain genes are susceptible to DoG transcription is that their 3’-end cleavage/poly(A) motifs deviate from the consensus and thus represent “weaker” transcriptional termination signals.^26^ We therefore examined all genes with 3’-end-associated binding of MYC+MAX and above-average DoG transcription levels from the cell lines and HCCs mentioned above and compiled consensus HOMER plots of presumptive cleavage/poly(A) signals residing within 100 bp upstream of their assigned TESs. When compared to genes from the same sources with below average DoG transcription levels and without 3’-end MYC+MAX binding, we found there to be little, if any, difference in these motifs (Figure 4M). Thus, genes with TES-associated MYC+MAX binding and high levels of DoG transcription do not appear to contain non-consensus or weaker cleavage/poly(A) signals.

### MYC and MAX recognize similar DNA motifs at 5’- and 3’-ends of their target genes

Although MYC-MAX heterodimers bind E boxes and drive transcription directly, their suppression of target genes is indirect and mediated via inhibitory interactions with MIZ1, SP1 and SP3. ^14,15^ Using the same data sets employed for generating the heat map of TF binding sites and their relation to TSSs, TESs and enhancers (Figure 4I), we performed motif enrichment using MEME Suite to compile a rank list of the consensus DNA binding motifs that overlapped the MYC-MAX footprints. ^37^ Not unexpectedly, E boxes and SP1 sites were among the top 3 most prevalent motifs identified, with MIZ1 sites also being significantly enriched. The distribution of these sites was similar at 5’- and 3’-ends of both human and mouse genes (Figure 5A & 5B). The tight clustering of E boxes around MYC and MAX binding peaks likely reflected direct E box occupancy by these heterodimers whereas the more diffuse distribution of Sp1 and Miz1 sites indicated binding sites at which Sp1 and Miz1 binding was both independent of and associated with MYC-MAX.

**Figure 5.**
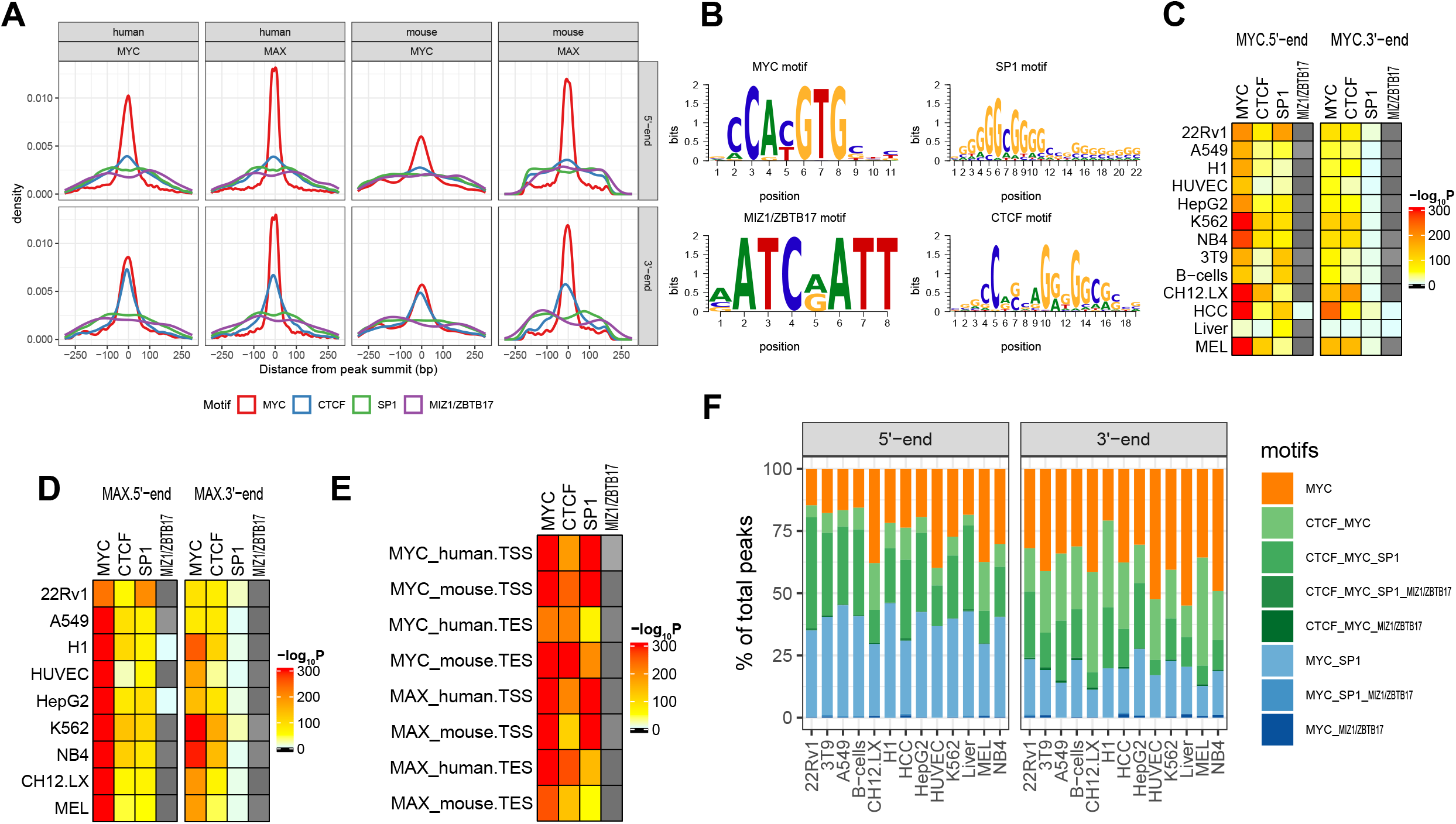
MYC and MAX binding around TSSs and TESs is associated with common DNA motifs. (A). Distribution of the top 4 TF binding sites in relation to MYC binding peaks around TSSs and TESs of all annotated genes in the human and mouse cell lines depicted in Figure 1A. Position “0” is defined as the peak of the MYC footprint. (B). HOMER plots showing consensus DNA binding motifs associated with ChIPseq peaks from panel A. (C). P values for the results in A indicating the non-random relationships of the indicated binding sites in relation to MYC ChIP signal peaks in TSS- and TES-associated sites. (D). P values for the results in A indicating the non-random relationships of the indicated binding sites in relation to MAX ChIP signal peaks in TSS- and TES-associated sites. (E). Combined results from all cell lines shown in C and D. (F). Association of MYC Chip footprints with other binding domains. ChIPseq results from the cell lines depicted in A were combined. All DNA sequences in which peak binding corresponded to an E box motif were then searched as in A for consensus elements for the additional indicated TFs. On average, <25% of the regions associated with MYC footprints were associated exclusively with E boxes.

Another top binding motif associated with 3’-ends was that for the multi-zinc-finger protein “CCCTC binding factor” (CTCF). Originally described as a suppressor of MYC-mediated transcription, CTCF was subsequently shown to possess more general roles in transcriptional insulation and repression as well as chromatin looping between promoters and enhancers.^31,38^. Some CTCF motifs contain loose approximations of E boxes embedded within their more complex consensus binding sequence (GCC^A^/_C_^C^/_T_CT^G^/ ^G^/),^31^ although they do not appear to comprise even low-affinity MYC+MAX binding sites ^39^. In general, CTCF motifs coincided more closely with those for MYC, particularly at TESs (Figure 5A). Thus, rather than being randomly distributed, all the above-described motifs and their respective bound factors clustered around MYC binding peaks in close proximity to TSSs and TESs (Figure 5C-5E).

### MYC-MAX binding at 5’- and 3’-ends of genes is associated with TSS-TES interactions

Direct contacts between proximal promoters and remote enhancers, mediated by CTCF-cohesin- facilitated DNA looping, allow other TFs bound at these sites, including MYC, to interact directly. ^17,40^ Similar contacts involving TSSs and TESs are believed to allow RNAPII to more efficiently return to promoters upon completing each round of transcription ^41,42^. Using ChIA-PET data from 5 evaluable cell lines in the ENCODE database ^43,44^, we assessed TSS-TES contacts (looping) among genes that bound RNAPII around both TSSs and TESs and MYC and/or MAX at TESs. In all cases examined, TSS-TES interactions were found to be more abundant when MYC and MAX co-bound at TES-proximal sites (Figure 6A). These genes were also expressed at significantly higher levels (Figure 6B).

**Figure 6.**
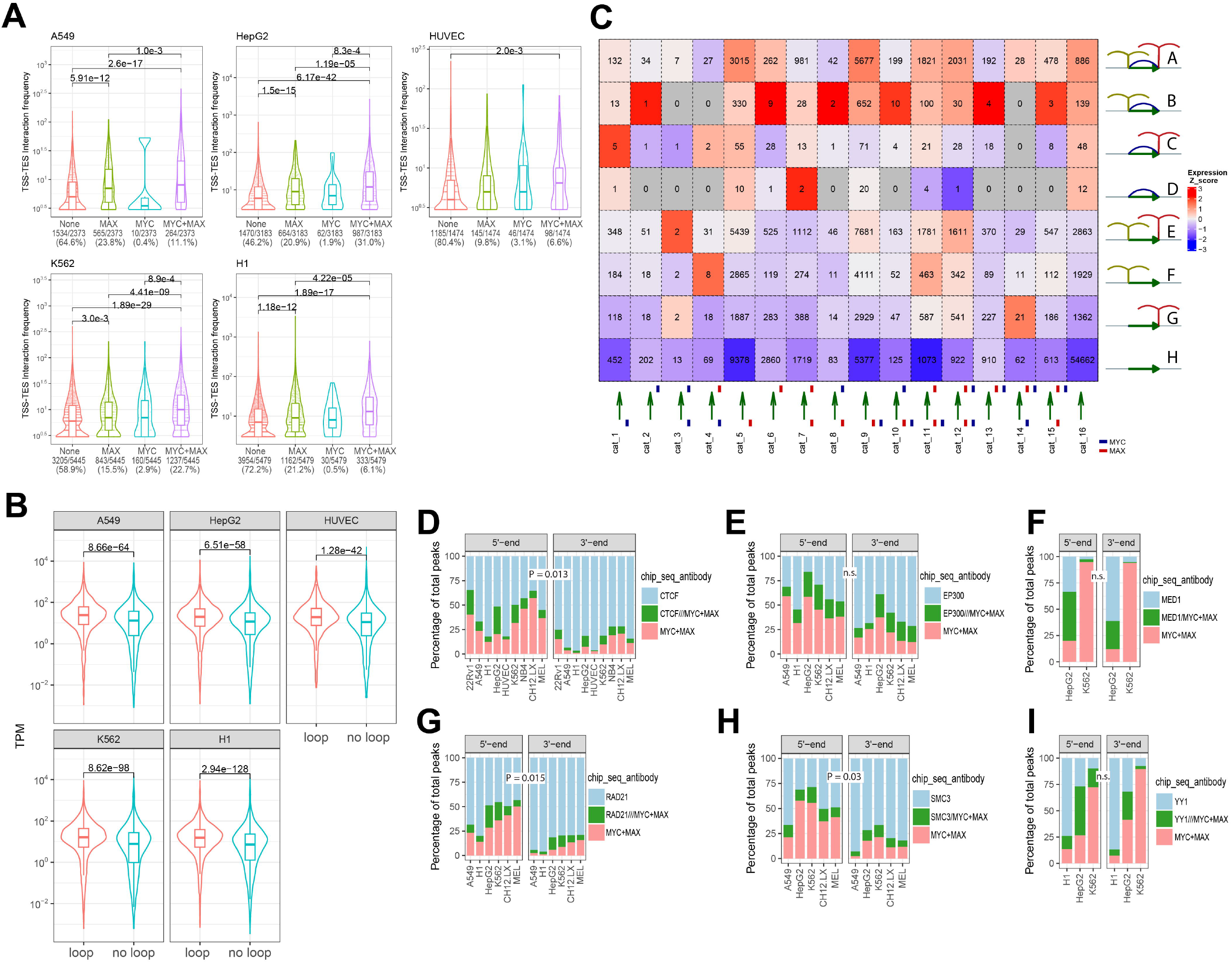
Intragenic contacts and expression are facilitated by TES-proximal MYC and MAX binding. (A) Gene interaction frequencies. For the indicated cell lines, all genes that bound RNAPII were divided into 4 groups based upon whether or not they also bound MYC and/or MAX at TES-proximal sites. Using ChIA-PET, TSS-TES interaction (looping) frequencies were then determined for each of these groups. Numbers below each panel indicate the number and percentage of genes in each category. (B). Using RNAseq data for the cell lines shown in A, the expression of genes with (loop) or without (no loop) TSS-TES looping was determined. (C). Combined gene expression heat maps and TSS-TES interactions for the cell lines shown in A. All genes that bound RNAPII at their 5’- and 3’-ends were divided into the 16 MYC and/or MAX binding categories originally shown in Figure 1A. These were further divided into the 8 groups shown at the right (A-H) according to types of TSS and TSS contacts that were identified based on ChIA-PET analyses ^18^. The blue curves shown in Groups A-D represent direct contacts between TSSs and TESs. Vertical ticks seen in all Groups except D and H indicate contacts between TSSs or TESs and additional intra- or extra-genic sites. Numbers within each of the 128 squares of the heat map indicate the total number of genes identified among the cell lines. (D) Binding of CTCF to regions +/-2.5 kb of TSSs and TESs. P value for differences in the total binding of these factors are indicated between the 2 sets of histograms. (E). Binding of p300 histone acetyl transferase to regions +/-2.5 kb of 5’-end TSSs and 3’-end TESs. (F). Binding of Mediator subunit MED1 to regions +/-2.5 kb of 5’-end TSSs and 3’-end TESs. (G,H). Binding of cohesion complex subunits RAD21 and SMC3, respectively, to regions +/-2.5 kb of 5’- end TSSs and 3’-end TESs. (I). Binding of the bi-functional transcription factor YY1 to regions +/-2.5 kb of 5’-end TSSs and 3’-end TESs.

For deeper insight into how MYC and MAX binding around TSSs and TESs influence DNA looping and transcription, all expressed genes in the above 5 cell lines were divided into the previously mentioned 16 categories (Figure 1A). ChIA-PET results were then used to further sub-classify these into 8 additional groups (A-H) based upon the types of contacts made by TSSs and TESs (Figure 6C). These included the TSS-TES contacts discussed above (Groups A-D) as well as other intra- and extra-genic interactions most likely involving enhancers. Combining these with the mean expression levels of these 128 individual gene sets yielded several observations. First, as already demonstrated (Figure 6B), gene Groups A-D were more highly expressed than those lacking TSS-TES contacts (Groups E-H); Second, expression among the 4 former groups was higher when their TSSs and TESs made additional intra- and extra-genic contacts (groups A-C vs. Group D). Third, MYC and MAX co-binding around TSSs and TESs was associated with higher looping frequencies both at TSSs and elsewhere. For example, twice as many contacts (81.6%) were seen among category 12 genes, with MYC+MAX bound at both ends, than among category 16 genes, with no MYC or MAX binding at either (40.2%). Fourth, significant levels of looping between TSSs and TESs could still occur when MYC-MAX binding occupied only one end or when binding at both ends was incomplete, i.e. 60.1% in the case of category 9 genes, 56.2% in the case of category 13 genes and 81.1% in the case of category 10 genes. Collectively, these findings indicate that the influence which MYC-MAX binding exerts upon gene expression is highly correlated with its TSS-TES contacts. However, this appears to involve additional and likely complex interactions with other TFs as well as intra- and extra-genic enhancers.

In addition to being enriched for RNAPII and CTCF, points of contact between promoters and enhancers also commonly contain the CTCF-cohesion complex, the EP300 transcriptional activator/acetyltransferase, the multi-subunit Mediator complex and the bi-functional transcription factor YY1 ^16,45,46^. Again using data from the ENCODE database for all or some of the cell lines, (Table S1)^18^ we mapped the binding sites for these factors or their key subunits.

Although somewhat less frequent at 3’-ends than at 5’-ends, these either directly overlapped MYC+MAX binding signal peaks or resided in close proximity to them (Figure 6D-6I). Thus, the protein complexes associated with MYC-MAX at sites of contact between TSS- and TES- containing regions resemble but are distinct from those associated with the points of contact between TSSs and enhancers.

### Loss of TES-associated MYC-MAX binding alters gene expression, basal and stress-induced DoG transcription and TSS-TES contacts

Many cell types contain genes with high levels of MYC-MAX binding to TSS- and TES-associated E boxes and robust DoG transcription (Figure 4D). ^18,26^ We identified 7 such genes in immortalized murine fibroblasts ^18^ and then used a Crispr/Cas9-based approach to delete/mutate their consensus TES E boxes (Figure 7A, Figure S8 and Table S3). We then asked how this impacted transcript levels during serum-starvation-mediated quiescence (0.1% FBS x 24 hr: “MYC-low”) and the subsequent response to a brief period of serum-stimulated endogenous Myc induction (10% FBS x 4 hr: “MYC-high”) ^47^. Relative to wild-type (WT) cells, transcript levels of 5 of the 7 genes were significantly reduced in MYC-low KO cells indicating that even minimal levels of TES-associated MYC binding are sufficient to maintain basal gene expression (Figure 7B). In contrast to this positive effect, transcript levels of most of the genes were down-regulated in MYC-high WT cells whereas their expression in MYC-high KO cells was either less affected by serum or, in 2 cases, up-regulated. Thus, for the limited number of genes examined, MYC binding to TES-proximal E boxes appeared to serve 2 purposes. First, it positively maintained gene expression when endogenous Myc levels were low as occurs during quiescence. Second, it determined both the direction and the magnitude of the gene’s response during periods of transiently high Myc expression as occurs after a growth-promoting stimulus such as serum stimulation.

**Figure 7.**
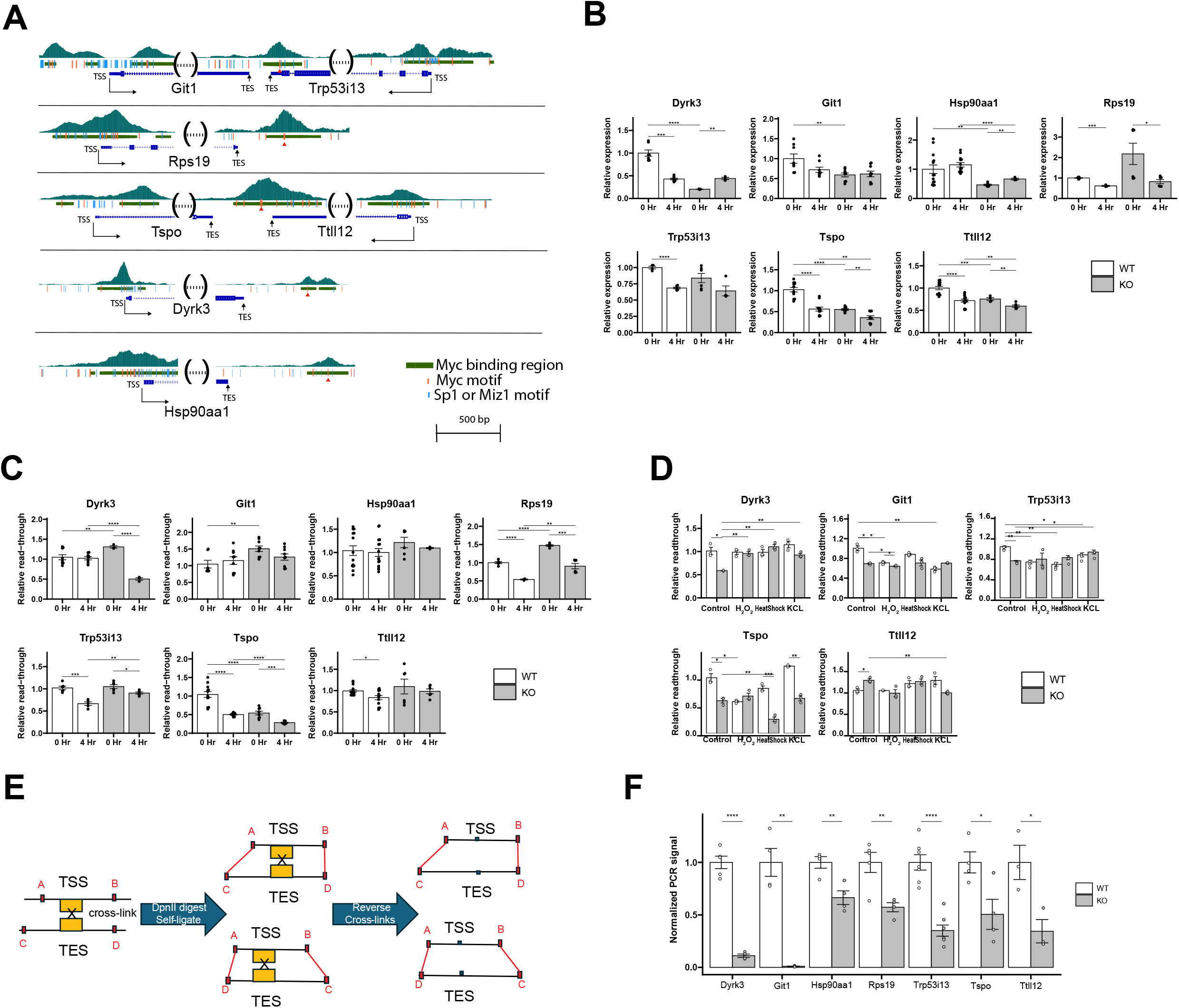
Loss of TES-proximal E boxes impacts total and read-through transcription and TSS-TES contacts. (A). Cartoons of the indicated 7 genes showing the locations of TSS- and TES-associated MYC binding sites in immortalized murine fibroblasts along with consensus E boxes and Sp1 and Miz1 sites.^18^ Note that the *Git1* and *Trpp53i13* and *Tspo* and *Ttll12* genes are arranged in tail-tail orientation and bind MYC-MAX at a single site that resides within <2.5 kb of each gene’s TES. (B). The indicated NIH 3T3 fibroblast lines containing WT or mutant TES-associated E boxes were grown to approximately 80% confluence and then placed for 24 hr in medium containing 0.1% FBS. One set of cells was then harvested at this point (time 0). Fresh medium containing 10% FBS was added to the remaining cells which were harvested 4 hr later. Transcript levels were determined in each clone by qRT-PCR using the primers indicated in Table S4 and normalized to those of transcripts for a control gene (TBP) as previously described. ^64^ (C). Basal read-through transcription is impacted by E box mutagenesis. Total RNAs were directionally reverse-transcribed using gene-specific forward primers, followed by qPCR. All primers used are listed in Table S4. (D). Changes in DoG transcription in response to a 4 hr exposure to heat shock (45° C), KCl (200 mM) and H_2_O_2_ (0.2 mM). DoG transcription was quantified as described in C. Control cells were grown under standard log-phase conditions. (E). General strategy for modified 3D chromatin conformation capture of WT and KO murine fibroblasts. The approach was designed to identify interactions between TSS- and TES-proximal regions regardless of their relative orientations as indicated in the upper and lower portions of the diagram. Black lines indicate the gene body. Step 1: formaldehyde cross-linking. Red boxes A-D indicate Dpn1 sites in closest proximity to TSSs, TESs and MYC-MAX binding sites. (orange boxes). Step 2: Cross-linked chromatin digested with DpnII and then self-ligated. Step 3: Cross-link reversal, SDS-proteinase K digestion and phenol-chloroform extraction. Step 4: PCR reactions designed to amplify fragments of defined sizes across all 4 possible ligated sites (Figure S9). Correct PCR products were confirmed based on their initial size and sizes of fragments observed following DpnII digestion and amplicon sequence. (F). Quantification of TSS-TES interactions. Using the primers shown in Figure S9 and Table S5, qPCR was performed on the indicated genes in WT and KO fibroblasts following formaldehyde cross-linking, DpnII digestion, re-ligation and reversal of cross-links as depicted in E. Results show the average of triplicate samples +/- 1 S.E. after DNA inputs were adjusted using a set of PCR primers that amplified a region of the *Mlx* gene. In some cases, PCR products from WT cells longer than those indicated were found to arise as a result of protection of one of the DpnII sites (panel E) by MYC-MAX. Instead, digestion occurred at the immediately adjacent site.

In the presence of low serum, DoG transcription of 3 genes (*Dyrk3, Git1* and *Rps19*) in KO cells was increased, implying that MYC binding near TESs normally suppresses this process (Figure 7C and Table S4). In contrast, the opposite effect was observed with the *Tspo* gene and no effects on DoG transcription were observed with *Hsp90aa1, Trp53i13* or *Ttll12*. In response to serum stimulation, DoG transcription in KO cells was decreased for 2 genes (*Dyrk3* and *Tspo*), increased for 2 genes (*Rps19* and *Trp53ai13*) and unchanged for the remaining 3 genes (*Git1, Hsp90aa1* and *Ttll12*). Thus, as was true with overall transcript expression, DoG responses in quiescent and serum-stimulated cells, and their regulation by TES-proximal MYC-MAX binding, were as varied as they were in genes that are driven solely by MYC binding near TSSs. 1,4,5,12,14,15

More so than serum deprivation, DoG transcription can also be impacted by other stresses such as heat, osmotic shock and reactive oxygen species, albeit in gene-specific ways.^26,27^ To determine whether TES-associated E box loss affected these responses, 5 of the above cell lines were exposed during log-phase growth to each of these stresses for 4 hr after which strand-specific qRT-PCR was performed to quantify the relative levels of DoG transcription (Table S4). In WT cells, 3 genes (*Git1, Trp53l13* and *Tspo*) were affected by at least one of the above stresses, although DoG transcription was reduced rather than increased (Figure 7D). In KO cells, DoG transcription was generally lower under control conditions, which largely replicated the results of serum-stimulated Myc induction. The responses of KO cells were both gene- and stress-specific and were not always decreased as they were in WT cells. Thus, TES-associated Myc binding can impact both the directionality, extent and specificity of stress-induced DoG transcription in ways that are highly gene-specific.

The above-noted changes in total and DoG transcription could have been facilitated via the formation of chromatin loops between MYC-MAX-bound TESs and TSSs in a manner analogous to that of promoter-enhancer contacts.^17,41,42,46,48,49^ To test this, we performed a modified and focused version of 3D chromatin conformation capture in which DNA bound by MYC-MAX was chemically cross-linked in WT and KO cell lines, digested with DpnII and then self-ligated prior to reversing the protein-DNA cross-links (Figure 7E). DpnII site junctions, in closest proximity to TSSs and TESs, were then amplified using PCR primer sets that flanked each of the 4 potential points of contact formed upon DpnII site re-ligation (Figure S9). The results of these studies showed that KO cells contained many fewer such re-ligated sites (Figure 7F). Thus, just as was true for basal and DoG transcription, chromatin looping and direct contact between TSSs and TESs appeared to be facilitated and maintained by the presence of MYC binding at the latter sites.

## Discussion

MYC and other TFs are traditionally portrayed as interacting with consensus DNA elements located in proximal promoters or enhancers, with the latter being classified as either intra- or extra-genic.^50,51^ The direct contacts between these sites occur via chromatin looping and are mediated by various post-translational histone modifications and protein-protein interactions ^16,28,29,32–34,45,46,51^. These facilitate access by more general components of the transcriptional machinery such as RNAPII, Mediator and their numerous co-factors that additively or synergistically promote mRNA initiation, capping, elongation and the reversal of pausing.^7,24,50,51^ Relying on data from the ENCODE Consortium, ^18^ we have identified heretofore unreported binding of MYC and/or MAX to the 3’-ends of over one-sixth of all annotated genes, with the vast majority of this binding centering around TESs (Figure 1A&1B and Figure S2). Its conservation over >700 million years of evolution thus precedes the bilaterian divergence and suggests important functional roles (Figure S7). ^22^ Typically, it involves co-binding by both MAX and MYC although binding by individual factors is not uncommon (Figure 1A). MAX binding in MYC’s absence is more frequent and probably reflects its additional MYC-independent interactions with various MXD members ^2,4^. In contrast, binding by MYC alone was 5 times less frequent and is consistent with the fact that both its direct binding to DNA and its inhibitory interactions with positively-acting TFs such as Miz1 and Sp1/3, occur in association with MAX.^2,4,14,15^ Overall, binding by MYC and/or Max at either or both 5’- and 3’-end of genes as defined here accounted for an average of 52.5% of all annotated genes (range 30.0-63.6%) across the 9 human and murine cell lines examined in this study (Figure 1A).

Initial comparisons of MYC and MAX binding revealed both similarities and differences with regard to 5’- and 3’-end binding profiles. For example, the binding of both factors centered around TSSs and TESs, although the footprints at the latter sites were broader (Figure 1 B and Figure S2). Both high- and low-affinity binding sites were observed with the latter being somewhat more prominent at 3’-ends (Figure 1E and 1G). In response to both physiologic and pathologic levels of MYC induction, changes in gene expression occurred within as little as 2 hr of MYC-MAX binding and in genes with multiple binding sites, those residing closest to TSSs and TESs tended to be of higher affinity and were occupied before those located more distally (Figures 1G and 3A-3I). Collectively, these findings indicate that MYC and MAX binding sites around TSSs and TESs have evolved and behave similarly with respect to their spatial organization, binding affinities and effects on gene expression in response to fluctuations in MYC protein levels.

Although MYC-regulated genes tend to oversee mitochondrial and ribosomal structure and function, cell cycle, redox regulation, senescence/aging and DNA damage recognition/repair, those with MYC-MAX binding sites in TES-proximal regions were somewhat more enriched for functions pertaining to differentiation, proliferation, cytokine/immune responses (Figure 3J and K).^2,4^ Genes encoding lncRNAs were also more likely to show evidence of TES-associated MYC/MAX binding (Figures 4A). The precise reasons for these differences remain to be determined, but nonetheless indicate that, like the temporal and spatial binding hierarchies of these sites, the functional preferences are also broadly conserved across cell types and species.

The persistence of MYC at both TSS- and TES-proximal sites in Myci975-treated 22Rv1 cells belied the more extensive changes in its total levels as previously determined by immuno- blotting (Figure 1E).^19^ Numerous studies have shown MYC protein half-life to be 15-30 min and that its levels decline by >90% within 12-24 hr of adding various Myc inhibitors.^19,21,52^ Reconciling these seemingly contradictory findings thus leads to the conclusion that, at any given time, a modest fraction of MYC is highly stabilized by virtue of being chromatin bound. This idea is consistent with the fact that newly synthesized MYC and MAX do not immediately associate since they are synthesized in and initially localize to different cytoplasmic compartments. ^53^ The stability of MYC-MAX at its DNA-bound sites may also partially reflect the fact that Myc inhibitors are much more effective at preventing MYC-MAX heterodimerization than in reversing it. ^54^ Furthermore, nuclear-localized DNA-bound MYC may be relatively under- phosphorylated at Thr_58_/Ser_62_ (a major determinant of MYC’s stability) and further shielded from proteasome-mediated decay. ^21,55,56^ The marked and rapid changes in gene expression following Myci975 treatment, despite the persistence of MYC-MAX binding, may thus reflect the tendency of many Myc targets to be exquisitely sensitive to even small changes in MYC levels (Figures 1E and 3A-3F).^4^ Cells may therefore contain 2 populations of MYC, the first of which is abundant, monomeric, inherently unstable and susceptible to MYC inhibitors. In contrast, the second population, while much less abundant, is highly stable by virtue of its association with Max, its sequestration by chromatin (possibly in an under-phosphorylated state) its protection from proteasomal degradation and its resistance to MYC inhibitors. Precisely how much DNA-bound Myc’s half life is extended, why gene expression is so sensitive to small changes in its levels and whether it serves non-canonical functions as a result of this persistence are questions that will be important to investigate in future work.

Both TSS- and TES-associated MYC and/or MAX binding and function occur in the context of numerous other TFs and co-factors that promote transcriptionally conducive changes in the local epigenetic landscape and an open chromatin network (Figures 4E-4I and 5 and File S1). This includes a high density of “paused” RNAPII that tends to accumulate around TESs where it plays important roles in transcriptional termination, mRNA cleavage and polyadenylation ^57,58^. Its differential binding pattern around MYC-MAX-bound sites versus those without MYC-MAX binding suggests that it also plays additional roles in gene regulation (Figure 4J). This might reduce the likelihood of mRNA termination and increase the probability of DoG transcription. Similarly, the distinct binding patterns of these various factors in the vicinity of TESs further suggests that their roles are different from those of either intragenic or distal enhancers (Figure 4I).

Four of the 7 genes whose 3’-end E boxes sites were mutated for the studies described above resided in tail-tail configurations and shared only single binding peaks of MYC-MAX in close proximity of one another’s TESs (*Git1-Trp53i13* and *Tspo-Ttl12*) (Figure 7A). In the latter case, the shared E box appeared to be necessary for the basal expression for both genes whereas in the former case, only the baseline levels of *Git1* was affected by E box mutation. These studies suggest that MYC-MAX binding to a single shared E box can differentially impact the expression of its adjacent genes. However, it remains possible that under different conditions or in different tissues, both qualitiative and quantitative variations in these patterns are possible.

Several features distinguish MYC/MAX-associated TESs from classical enhancers. The first and most notable of these is their predictable and nearly always solitary location relative to TESs in contrast to enhancers whose numbers and positions cannot be ascertained *a priori* (Figure 1B).^16,29,32,33,45,46,59^ Second is their association with a prominent landmark, namely the cleavage/poly(A) site. Third is a pattern of TF/co-factor binding that distinguishes them from both promoters and enhancers (Figure 4I). Fourth, unlike enhancers, which positively regulate transcription, MYC-MAX-bound TES regions can also be suppressive. Finally, they participate directly in both basal and stress-responsive DoG transcription Figures 1C, 4K and 7C & 7D). DoG transcripts may display novel properties that include their refractoriness to nuclear export, their maintenance of euchromatin structure and their read-through into and activation of adjacent downstream oncogenes that can impact cancer. ^27,60^

Despite the above differences, MYC-MAX associated TESs do appear to share some of the functions of classical enhancers as determined by their interactions with both TSSs and true enhancers in a manner that involves direct contacts that are mediated by DNA looping (Figures 6C and 7E and 7F). In the limited number of genes that we examined directly, we identified both positive and negative impacts of TES-associated MYC-MAX binding on total and DoG transcription. This binding appears to represent a somewhat more flexible and perhaps specific way of regulating total and DoG transcription than can be afforded solely by enhancers, particularly since it involves more intimate contacts between sites of mRNA initiation and termination. The ability of MYC-MAX heterodimers bound at one site to interact with those at another and facilitate looping (Figure 7F) is also in keeping with the long-known ability of MYC and MAX to heterotetramerize.^61^ At the same time, it should be stressed that, despite the importance of MYC-MAX in the formation of 5’- and 3’-end contacts, it seems likely that this represents only one of many factors that are important for establishing these loops, which are likely to be transient in nature, highly tissue-specific, and subject to a variety of both physiologic and stress-related factors and post-translational modifications (Figures 5A, 6D-6I and 7).

The studies reported here have focused on MYC and its role in regulating gene expression via binding to TES-proximal regions of genes. However, it is clear from our analyses that the chromatin epigenetic landscape is associated with and modified accordingly by many factors other than MYC-MAX (Figures 4I and 6D-6I). Precisely how these and the modifications they impart influence gene expression, contribute to or modify MYC-MAX DNA binding, and further enable DNA looping and contact with TSSs and enhancers will require individualized attention.

## Supporting information

Key Resource Table

## ACKNOWLEDGEMENTS

The analysis of ENCODE and GEO data was, in part, supported by the University of Pittsburgh Center for Research Computing. Specifically, work utilizing the HTC cluster was funded by NIH award number S10OD028483. Funding for this project was provided by grants from the NIH (RO1 CA174713), The Rally Foundation (no. 22N42) and a Hyundai Hope on Wheels Scholar grant (all to EVP). Additional support was provided by The UPMC Children’s Hospital Foundation.

## AUTHOR CONTRIBUTIONS

HW compiled and analyzed genomics data and performed ChIP studies; BM, TS, JK and JL generated and characterized cells lines and conducted qRT-PCR studies; EVP conceived the study and EVP and HW devised experiments, analyzed data and wrote the manuscript. All authors read and approved the final version of the manuscript prior to submission for publication.

## STAR**•**METHODS

Detailed methods and provided in the online version of this paper and include the following:

- KEY RESOURCES TABLE
- RESOURCE AVAILABILITY

- Lead contact
- Materials availability
- Data and code availability
- EXPERIMENTAL MODEL and subject DETAILS
- METHOD DETAILS

- Computational methods
- gRNA vectors
- Generation and characterization of murine fibroblasts lacking TES-proximal E boxes.
- Quantification of total transcripts, DoG transcription and TSS-TES contacts.
- PCR based assay for TES-TSS interactions.
- Quantification and statistical analysis
- ADDITIONAL REFERENCES

## STARfllMethods

### Resource availability

#### Lead contact

Further information and requests for resources and reagents should be directed to and will be fulfilled by the lead contact, Edward Prochownik.

### Materials availability

All unique reagents generated in this study will be made available from the Lead Contact (E.V.P.) and may require a completed Materials Transfer Agreement.

## Data and code availability

This paper does not report original code and new high throughput sequence data. ChIPseq, DNaseseq, ATACseq, ChIA-PET and RNAseq datasets utilized in this study were obtained from the ENCODE or GEO databases. A complete list of all metadata employed in the current study can be found in Table S1.

Any additional information required to reanalyze the data reported in this paper is available from the lead contact upon request.

## Experimental model and study participant details

### Materials and Methods

#### Computational methods

ChIPseq, DNaseseq, ATACseq, ChIA-PET and RNAseq datasets utilized in this study were obtained from the ENCODE or GEOdatabases. A complete list of all metadata employed in the current study can be found in Table S1.

All alignments were performed using GRCh38, mm10 and dm6 as the reference sequences for human, mouse and *D. melanogaster* genomes, respectively. The "gencode.vM21.primary_assembly.annotation_UCSC_names.gtf" (EMCODE accession: ENCSR884DHJ) and "gencode.v29.primary_assembly.annotation_UCSC_names.gtf" (ENCODE accession: ENCSR884DHJ) annotation files ^75^ were used as the reference annotations for mouse and human, respectively.

For the ENCODE ChIPseq datasets, the IDR (Irreproducible Discovery Rate) thresholded peaks provided by ENCODE were directly used as the processed data. For GEO ChIPseq datasets, fastq files were downloaded from the NCBI SRA (Sequence Read Archive) database. Subsequently, these files were analyzed using the ENCODE Transcription Factor and Histone ChIP-Seq processing pipeline (ENCODE-DCC/chip-seq-pipeline2-2.2.0)^66^, which incorporates the ENCODE integrative analysis approach.^18^

ChIP peak annotation was performed using the ChIPpeakAnno package (version 3.6.5) in the R/Bioconductor environment. IDR thresholded peaks, obtained in the "narrowPeak" format, were subjected to further analysis. To prepare IDR thresholded peaks for annotation, they were transformed from the "narrowPeak" format into GrangesList forms. The transformed peaks served as the input data for the subsequent annotation process. The annotatePeakInBatch function takes the transformed GrangesList forms of ChIPseq peaks as input and assigns functional annotations to each peak based on its genomic coordinates. In this analysis, the binding regions for all genes of interest were defined as ±2.5 kb (±1 kb for *D. melanogaster*) relative to each transcript’s TSS or TES. To annotate the ChIP peaks, the appropriate AnnotationData for the respective organisms was utilized. For human data, the TxDb.Hsapiens.UCSC.hg38.knownGene(3.17.0) database was used. For mouse data, the TxDb.Mmusculus.UCSC.mm10.knownGene(3.9.0) database was employed and for *D. melanogaster,* the TxDb.Dmelanogaster.UCSC.dm6.ensGene database was utilized.

For ENCODE RNAseq datasets, gene quantification files available from the ENCODE website were directly utilized as the processed data. For GEO RNAseq datasets, the raw FASTQ files were downloaded from the NCBI SRA database. These files were then analyzed using the nf-core-rnaseq-3.4 pipeline. ^68^

For gene set over-representation analyses, gene sets were obtained using the R package msigdbr (version 7.5.1) and subsequently refined to exclusively include sets categorized as “C5: Gene Ontology resource (GO)”. The R package ClusterProfiler (version 4.10.0) from Bioconductor ^70^ was employed to conduct over-representation analysis using the enrichGO function, with a filter applied to retain results featuring adjusted p-values less than 0.05.

To define transcriptional landscapes of promoters and enhancers Cis Regulatory Module (CRM) annotations were derived from the ReMap2022 database, which integrates ChIPseq results from multiple sources and biotypes and thereby provides a multi-cellular multi-tissue regulatory map. ^76^ ReMap2022 covers 1210 TFs and co-factors from 737 human cell lines and tissues, 8103 QC ChIPseq datasets and 182 million binding sites. CRM annotations were first filtered so as to include all regions with MYC binding in common (File S1). Based on mapping to the human genome (version GRCh38), these “Myc CRMs” were assigned to TSSs or TESs as previously noted (Figure 1A and 1B). For all Myc CRMs sits, CCRE annotation ^18^ was used to determine whether they met the criteria as enhancers which were then assigned to regions within genes (intra-genic enhancers) or to distal regions (extragenic enhancers) located >2.5 kb upstream of TSSs or >2.5 kb downstream of TESs. The percentage of other transcriptional regulators associated with MYC in these regions was also calculated and then transformed to z-score. Each factor’s z-score in each cCRE-genome region was used as a matrix for heatmaps, which were generated using the ComplexHeatmap R package.

Motif analyses to identify putative bind sites at 5’- and 3’-ends of their target genes were performed in R using memes 1.10.0/ MEME Suite 5.5.0 and universalmotif. Enriched motifs were identified using the runAME function from memes with motif Databases HOCOMOCO Human (v11 FULL) as known motif database and a control set to ’shuffle’ for the input sequences. The runFimo function was used to identify individual motif occurrences.

For read through/DoG analyses, we used ENCODE, strand-specific RNAseq datasets, which were also rRNA-depleted. The output BAM files were generated by aligning to either the human genome (assembly: GRCh38, genome annotation: V29) or the mouse genome (assembly: mm10, genome annotation: M21). GEO total RNAseq was aligned using the same genome assembly and annotation, with detailed information listed in Table S1. Additional analyses utilized DoGFinder software^74^ was used. Two different gene boundaries were considered, corresponding to stranded and unstranded RNAseq, respectively. Gene boundaries were set so as to be as inclusive as possible based on gene annotation V29 (human) or M21 (mouse), considering strand specificity or lack thereof. We defined the read count in the 500 bp downstream of the TES as read through and the 500 bp upstream of the TES as total gene expression. The Get_DoGs_rpkm function was used for read counting total and read through.

Chromatin Interaction Analysis Using Paired-End Tag Sequencing (ChIA-PET) data from ENCODE utilized RNA polymerase II subunit A (POLR2A) results.^44^ Interaction loop information files in bedpe format were downloaded from the ENCODE database, and comprehensive details are available in Table S1. The assignment of loop ends was performed by associating them with gene ends, employing the same approach as the binding assignment for MYC/MAX gene ends.

<u>gRNA vectors</u>. gRNA sequences were generated to allow targeting of MYC+MAX-binding E boxes of interest residing in proximity of TESs. The Crispr/spCas9 cloning vector pDG458, which encodes EGFP and can accommodate 2 gRNAs, was obtained from Addgene (Watertown, MA). Plasmid DNAs were purified using Qiagen columns according to the supplier’s protocol (Qiagen). gRNAs were directionally cloned into the vector using a single-step protocol (https://media.addgene.org/data/plasmids/100/100900/100900-attachment_Yl0i43bWJig3.pdf). Following introduction into chemically competent *E. coli* (Thermo Scientific DH5α competent cells, Thermo-Fisher, Pittsburgh, PA), plasmid DNAs from isolated bacterial colonies were purified and the identities and orientations of the gRNA-encoding inserts were confirmed by automated DNA sequencing. All gRNA-encoding oligonucleotides were chemically synthesized by IDT, Inc., Coralville, IA and are shown in Figure S8 and Table S3.

### Generation and characterization of murine fibroblasts lacking TES-proximal E boxes

NIH 3T3 murine fibroblasts were obtained from the ATCC (Manassas, VA) and routinely cultured in Dulbecco’s-modified minimal essential medium supplemented with 10% fetal bovine serum (FBS) (Atlanta Biological, Inc. Flowery Branch, GA) L-glutamine (200 μM), penicillin (100 units/ml) and streptomycin (10 μg/ml) (all from Sigma-Aldrich, Inc. St. Louis, MO). Cells were seeded the day before transfection into 6 well plates so as to be 30-50% confluent at the time of transfection. Transfections with gRNA-encoding pDG458 vectors (3 μg) were performed using Lipofectamine (Thermo-Fisher, Inc. Pittsburgh, PA) according to the directions of the supplier. Because each vector encoded gRNAs targeting 2 different genes, it could be used to eliminate or mutate the E boxes in 2 genes such that a single NIH3T3 clone could potentially harbor E box mutations in both genes. 2-3 days after transfection, single EGFP+ cells were sorted into 96 well plates using a FacsAria IIu Cell Sorter (BD Biosciences, San Jose, CA) and expanded into individual clones. Total cellular DNAs from randomly selected clones were then purified using DNeasy kits (Qiagen). PCR reactions (12 μl) using 100 ng of input genomic DNA and Pfu DNA Polymerase (Promega, Inc. Madison, WI) were performed to amplify regions flanking the targeted E boxes using the PCR primers listed in Table S3. Aliquots of PCR products from genes with consensus palindromic CACGTG E boxes were digested with 5 units the restriction enzyme PmlI to determine whether the product was fully or partially resistant to digestion and thereby indicating the presence of an E box mutation. PCR fragments from genes containing non-consensus E box elements were melted, re-annealed and digested with one unit of T7 endonuclease 1 according to the protocol of the supplier (New England Biolabs, Inc., Ipswich, MA). This permitted the identification of heteroduplexes between WT and mutant sequences or between two different mutant sequences. Upon confirming that all E boxes had been mutated or deleted, PCR products were subjected to amplicon sequencing to identify all mutations that were present (Azenta Life Sciences, Chelmsford, MA). Because NIH3T3 cells are known to be hypertriploid, more than 2 mutant alleles were often detected (Figure S8) (https://nih3t3.com).

### Quantification of total transcripts, DoG transcription and TSS-TES contacts

RNA extractions were performed using RNeasy columns (Qiagen) and cDNAs were synthesized by SuperScript™ IV First-Strand Synthesis System as previously described.^64^ Transcript levels were quantified by qRT-PCR using the Syber Green method using the primers listed in Table S4.^2,77^ These were normalized against a control qRT-PCR reaction for transcripts encoding TBP. DoG transcripts were quantified by first performing strand-specific reverse transcription for each gene followed by qRT-PCR as described above. DoG transcript levels were then normalized to those obtained for total transcript levels. Strand-specific RT primers and those for subsequent qPCR are listed in Table S4.

### PCR based assay for TES-TSS interactions

A qPCR-based assay was employed to quantify interactions between TES and TSS regions as indicated in Figure 7E. 1x10^7^ log-phase WT and E box mutant NIH3T3 cells were cross-linked with 1% formaldehyde (FA) for 20 minutes, quenched with 0.2 M glycine and washed with DPBS. The FA-crosslinked cells were further cross-linked with 2 mM EGS [ethylene glycol bis(succinimidyl succinate)] for 45 minutes, quenched again with 0.2 M glycine and washed with DPBS. The cross-linked cells were lysed with 0.1% SDS in the presence of protease inhibitors(Protease Inhibitor Cocktail, MilliporeSigma) at 4°C for 1 hr. Nuclei were further permeabilized with 0.55% SDS at 25°C for 10 min, followed by incubation at 62°C for 10 min, and then at 37°C for 10 min. SDS was quenched with Triton X-100 at 37°C for 15 min. The chromatin was then digested to completion with 400 U of DpnII (New England Biolabs, Inc., Ipswich, MA) in 1X DpnII NEBuffer™ at 37°C for 24 hours. The enzyme was inactivated at 65°C for 20 minutes and de-salted using a 3 kDa cutoff centrifugal filter (Millipore, Inc. Burlington, MA). The DpnII-digested chromatin was next ligated in 1X NEB ligase buffer with 4000 U of NEB T4 ligase overnight at 16°C and then at 4°C for an additional 48 hours. FA cross-links were then reversed by adding 1mg/ml proteinase K and incubating at 65°C overnight. Following extraction with phenol/chloroform/isoamyl alcohol and precipitated with ethanol, the success of the digestion and ligation steps were verified using agarose gel electrophoresis. Ligated DNAs were quantified using a NanoDrop spectrophotometer. SybrGreen-based PCR reactions were used to amplify the TES-TSS junctions, with primer sequences and positions shown in Figure S9 and Table S5. The amplicon from each PCR reaction was verified by amplicon sequencing (Azenta Life Sciences, Waltham, MA). Final PCR products were also checked by agarose gel electrophoresis at the end of each assay to ensure that only single bands had been obtained, and could be digested with DpnII to generate products of the predicted size. A *Mlx* gene region lacking DpnII site was used for normalization by TaqMan PCR assay (forward primer sequence: 5’-AGTCCGCTGGCTTGTTT- 3’, reverse primer sequence: 5’-TTGACCCAAGGGTCCTC-3’, probe sequence:c 5’-/56- FAM/CGGTTCGGT/ZEN/AGGTTCACGATGACG/3IABkFQ/-3’).

### Statistical analyses

Quantification and statistical analysis were performed using R software v4.3 (R Foundation for Statistical Computing, Vienna, Austria) and GraphPad Prism v9.00 (GraphPad Software Inc., USA). Data are shown as mean ± SEM. When comparing 2 groups, a 2-tailed unpaired Student’s t test or a Mann-Whitney U test was performed. When comparing >2 groups, one-way ANOVA or Kruskal–Wallis tests were used. Correlative analyses were performed using a Pearson correlation test. All experiments were performed on at least 3 biological replicates.

## SUPPLEMENTAL INFORMATION TITLES AND LEGENDS

Supplemental Tables Table S1-S5

Supplemental Figures ------- Figure S1-S9 Supplemental File File S1

Table S1. Cell lines and data sets used in the current study.

Table S2. 190 genes associated with higher levels of TES-associated MYC+MAX binding and read-through transcription in HCCs vs. livers.

Table S3. Oligonucleotides used to create pDG458 gRNA constructs for CRISPR-Cas9 targeting of TES-associated E boxes and PCR primers for amplifying regions flanking the targeted E boxes.

Table S4. qRT-PCR primers used to quantify gene expression and DoG expression in WT and KO fibroblasts (Figure 7B-7D).

Table S5. qRT-PCR primers used to quantify TES-TSS region interactions(Figure 7F).

Figure S1. **Distribution of MYC and/or MAX binding sites residing within +/- 2.5 kb of TSSs and TESs.** Results of MYC and MAX binding were obtained from the ENCODE and GEO databases (Table S1). AnnotatePeakInBatch (ChIPpeakAnno Version 3.6.5) was used to assign binding sites.

Figure S2. Binding of MYC and MAX around TSSs and TESs of genes for the individual cell lines shown in Figure 1B.

Figure S3. Circos plots showing the chromosomal locations of human genes that bind MYC only in the vicinity of TSSs and/or TESs in each of the 7 cell lines used in this study. Red dots: locations of genes associated with MYC binding near TSSs only; green dots: locations of genes associated with MYC binding near TESs only; blue dots: locations of genes associated with MYC binding near both sites.

Figure S4. Circos plots showing the chromosomal locations of human genes that bind MAX only in the vicinity of TSSs and/or TESs in each of the 7 cell lines used in this study. Red dots: locations of genes associated with MAX binding near TSSs only; green dots: locations of genes associated with MAX binding near TESs only; blue dots: locations of genes associated with MAX binding near both sites.

Figure S5. Circos plots showing the chromosomal locations of mouse genes that bind MYC only in the vicinity of TSSs and/or TESs in each of the tissues and cell lines used in this study. Red dots: locations of genes associated with MYC binding near TSSs only; green dots: locations of genes associated with MYC binding near TESs only; blue dots: locations of genes associated with MYC binding near both sites. The last diagram shows the combined results similar to that shown for human genes in Figure 1F.

Figure S6. Circos plots showing the chromosomal locations of mouse genes that bind MAX only in the vicinity of TSSs and/or TESs in each of the cell lines used in this study. Red dots: locations of genes associated with MAX binding near TSSs only; green dots: locations of genes associated with MAX binding near TESs only; blue dots: locations of genes associated with MAX binding near both sites. The last diagram shows the combined results similar to that shown for human genes in Figure 1F.

Figure S7. Circos plots showing the chromosomal locations of *D. melanogaster* genes that bind MAX only in the vicinity of TSSs and/or TESs. Gene locations are based on ChIP-seq results obtained from third instar larvae.^18^ Red dots: locations of genes associated with dMYC or dMAX binding around TSSs only; green dots: locations of genes associated with dMYC or dMAX binding around TESs only; blue dots: locations of genes associated with dMYC or dMAX binding at both TSSs and TESs.

Figure S8. Mutagenesis of TES-associated E boxes identified as sites of MYC+MAX binding in immortalized murine fibroblasts.

The top portion of each panel displays the WT gene sequence surrounding the TES-associated E box, and the gRNA used for Crispr-mediated editing. Beneath this are the mutant sequences identified subsequent to Crispr-Cas9 targeting of the sites. In keeping with the fact that NIH3T3 cells are hyperdiploid,^78^ all clones contained between 3 and 6 different mutant alleles.

Figure S9. Strategy for quantifying TSS-TES E box contacts in murine fibroblasts and list of qPCR primers used for this purpose.

Each circle shows the expected structure and size (in bps) of loops formed between TSS- and TES-proximal regions (green and brown thick lines, respectively) following DpnII digestion of formaldehyde cross-linked genomic DNA and re-ligation. Locations of ligated DpnII sites and E boxes (or their previous locations) are indicated. Sequences of PCR primers used to amplify ligated regions flanking the sites of DpnII digestion/re-ligation are indicated as are the predicted sizes of the amplified DNA products.

**File S1.** Rank of 1210 TFs and transcriptional co-factors and the frequency with which they bound to MYC-associated TSSs, TESs and intragenic and distal enhancer elements. The ReMap2022 database (https://remap2022.univ-amu.fr/) was used a source of binding profile.

