## Supplementary material for "MYC Binding Near Transcriptional End Sites Regulates Basal Gene Expression, Read-Through Transcription and Intragenic Contacts": Key Resource Table

**Key resources table**

| **REAGENT or RESOURCE** | **SOURCE** | **IDENTIFIER** |
| --- | --- | --- |
| **Chemicals, peptides, and recombinant proteins** | | |
| L-Glutamine | Gibco | Cat#25030081 |
| Trypsin-EDTA (0.25%) | Corning | Cat# 25-053-CI |
| Penicillin-Streptomycin | Corning | Cat# 30-002-CI |
| DMEM | HyClone | Cat#SH30243.01 |
| Fetal Bovine Serum | Biowest | Cat# S1560 |
| Formaldehyde solution | MilliporeSigma | Cat# 47608-250ml-F |
| DPBS | Gibco | Cat#14190250 |
| EGS (ethylene glycol bis(succinimidyl succinate)) | Thermo Scientific | Cat#21565 |
| DMSO | MilliporeSigma | Cat#D2650 |
| Protease Inhibitor Cocktail | MilliporeSigma | Cat# 11836170001 |
| Glycine | MilliporeSigma | Cat# 1005901000 |
| **Critical commercial assays** | | |
| Lipofectamine™ 3000 Transfection Reagent | Thermo Fisher | Cat#L3000015 |
| Taq DNA Polymerase | NEB | Cat#M0273 |
| DpnII | NEB | Cat#R0543S |
| PmlI | NEB | Cat#R0532S |
| T4 DNA Ligase | NEB | Cat#M0202L |
| Proteinase K | MilliporeSigma | Cat# 3115836001 |
| Protease Inhibitor Cocktail | MilliporeSigma | Cat# 11836170001 |
| Thiazole Green (SYBR® Green I) | Biotium | Cat# #40086 |
| DNeasy Blood & Tissue Kits | QIAGEN | Cat# 69504 |
| RNeasy Mini Kit | QIAGEN | Cat# 74104 |
| QIAGEN Plasmid Maxi Kit | QIAGEN | Cat# 12162 |
| SuperScript™ IV First-Strand Synthesis System | Fisher Scientific | Cat# 18091050 |
| T7 Endonuclease I | NEB | Cat# M0689S |
| MAX Efficiency™ DH5α Competent Cells | Fisher Scientific | Cat# 18258012 |
| Triton™ X-100 | MilliporeSigma | Cat# X100-5ML |
| Amicon ® Ultra Centrifugal Filter, 3 kDa MWCO | MilliporeSigma | Cat#UFC500308 |
| **Deposited data** | | |
| ENCODE database | ENCODE Project Consortium^18^ | https://www.encodeproject.org/ |
| Datasets identifiers from ENCODE | In this paper | Table S1 |
| GEO database | NCBI GEO^20^ | https://www.ncbi.nlm.nih.gov/geo/ |
| Datasets identifiers from GEO | In this paper | Table S1 |
| Experimental models: Cell lines | | |
| NIH/3T3 | ATCC | Cat# CRL-1658; RRID:CVCL_0594 |
| **Oligonucleotides** | | |
| gRNA Oligonucleotides for pDG458 constructs | IDT | Table S3 |
| PCR primers for flanking region of the targeted E boxes | IDT | Table S3 |
| qRT-PCR primers for Figure 7B-7D | IDT | Table S4 |
| qRT-PCR primers for Figure 7E-7F | IDT | Table S5 |
| **Recombinant DNA** | | |
| pDG458 | Adikusuma et al ^65^ | RRID: Addgene_100900 |
| **Software and algorithms** | | |
| ENCODE-DCC/chip-seq-pipeline2-2.2.0 | Hitz et al.^66^ | https://github.com/ENCODE-DCC/chip-seq-pipeline2 |
| ChIPpeakAnno (version 3.6.5) | Zhu et al.^67^ | https://bioconductor.org/packages/release/bioc/html/ChIPpeakAnno.html; RRID:SCR_012828 |
| TxDb.Hsapiens.UCSC.hg38.knownGene(3.17.0) | Bioconductor Core Team | https://bioconductor.org/packages/release/data/annotation/html/TxDb.Hsapiens.UCSC.hg38.knownGene.html |
| TxDb.Mmusculus.UCSC.mm10.knownGene(3.9.0) | Bioconductor Core Team | https://bioconductor.org/packages/release/data/annotation/html/TxDb.Mmusculus.UCSC.mm10.knownGene.html |
| nf-core-rnaseq-3.4 pipeline | Ewels et al^68^ | https://nf-co.re/rnaseq/3.4 |
| msigdbr(version 7.5.1) | Subramanian et al ^69^ | https://CRAN.R-project.org/package=msigdbr; RRID:SCR_022870 |
| ClusterProfiler (version 4.10.0) | Wu et al ^70^ | <https://www.bioconductor.org/packages/release/bioc/html/clusterProfiler.html>;  RRID:SCR_016884 |
| ComplexHeatmap | Gu et al ^71^ | <https://www.bioconductor.org/packages/release/bioc/html/ComplexHeatmap.html>;  RRID:SCR_017270 |
| memes | Nystrom et al ^72^ | https://www.bioconductor.org/packages/release/bioc/html/memes.html |
| MEME | Bailey et al ^73^ | <https://meme-suite.org/meme/index.html>;RRID:SCR_001783 |
| universalmotif | Tremblay et al | https://www.bioconductor.org/packages/release/bioc/html/universalmotif.html |
| DoGFinder | Wiesel et al ^74^ | https://github.com/shalgilab/DoGFinder |
| R software v4.3 | R Core Team (2021) | https://www.R-project.org/ |
| R studio | Posit | https://posit.co/download/rstudio-desktop/;  RRID:SCR_000432 |
| ggplot2 | CRAN | https://ggplot2.tidyverse.org ;  RRID:SCR_014601 |
| GraphPad Prism v9.00 | GraphPad Software | [www.graphpad.com](http://www.graphpad.com);  RRID:SCR_002798 |
