## Supplemental information for "MYC Binding Near Transcriptional End Sites Regulates Basal Gene Expression, Read-Through Transcription and Intragenic Contacts"

Supplemental Tables  ------- Table S1-S5

Supplemental Figures ------- Figure S1-S9

**Table S1.** Cell lines and data sets used in the current study

| Assay type | Database | File accession ID | Experiment accession | Genome assembly | Biosample term name | Biosample organism | Experiment target | Biosample treatments |
|---|---|---|---|---|---|---|---|---|
| ChIP-seq | ENCODE | ENCFF598BZD | ENCSR000DYC | GRCh38 | A549 | Homo sapiens | MYC-human | |
| ChIP-seq | ENCODE | ENCFF713RHL | ENCSR000BTJ | GRCh38 | A549 | Homo sapiens | MAX-human | |
| ChIP-seq | ENCODE | ENCFF342ASE | ENCSR000ERN | mm10 | CH12.LX | Mus musculus | MYC-mouse | |
| ChIP-seq | ENCODE | ENCFF349LDZ | ENCSR000ERL | mm10 | CH12.LX | Mus musculus | MAX-mouse | |
| ChIP-seq | ENCODE | ENCFF270GMO | ENCSR000EBY | GRCh38 | H1 | Homo sapiens | MYC-human | |
| ChIP-seq | ENCODE | ENCFF460LNS | ENCSR000BSJ | GRCh38 | H1 | Homo sapiens | MAX-human | |
| ChIP-seq | ENCODE | ENCFF239IMY | ENCSR784BVD | GRCh38 | HepG2 | Homo sapiens | MYC-human | |
| ChIP-seq | ENCODE | ENCFF254ZDA | ENCSR168DYA | GRCh38 | HepG2 | Homo sapiens | MAX-human | |
| ChIP-seq | ENCODE | ENCFF608CXN | ENCSR000EGJ | GRCh38 | K562 | Homo sapiens | MYC-human | |
| ChIP-seq | ENCODE | ENCFF822FKQ | ENCSR000BLP | GRCh38 | K562 | Homo sapiens | MAX-human | |
| ChIP-seq | ENCODE | ENCFF152JNC | ENCSR000EUA | mm10 | MEL | Mus musculus | MYC-mouse | |
| ChIP-seq | ENCODE | ENCFF262ITC | ENCSR000ETX | mm10 | MEL | Mus musculus | MAX-mouse | |
| ChIP-seq | ENCODE | ENCFF858ZYN | ENCSR000EHR | GRCh38 | NB4 | Homo sapiens | MYC-human | |
| ChIP-seq | ENCODE | ENCFF793GVV | ENCSR000EHS | GRCh38 | NB4 | Homo sapiens | MAX-human | |
| ChIP-seq | ENCODE | ENCFF562ZOV | ENCSR000EEZ | GRCh38 | endothelial cell of umbilical vein (HUVEC) | Homo sapiens | MAX-human | |
| ChIP-seq | ENCODE | ENCFF459QFK | ENCSR000DLU | GRCh38 | endothelial cell of umbilical vein (HUVEC) | Homo sapiens | MYC-human | |
| ChIP-seq | ENCODE | ENCFF327QOB | ENCSR999ZCR | dm6 | whole organism | Drosophila melanogaster | Myc-dmelanogaster | |
| ChIP-seq | ENCODE | ENCFF629XNY | ENCSR672KPS | dm6 | whole organism | Drosophila melanogaster | Max-dmelanogaster | |
| ChIP-seq | ENCODE | ENCFF341QUK | ENCSR191VCQ | dm6 | whole organism | Drosophila melanogaster | Myc-dmelanogaster | |
| ChIP-seq | ENCODE | ENCFF409EHG | ENCSR949RDZ | dm6 | Kc167 | Drosophila melanogaster | Myc-dmelanogaster | |
| ATAC-seq | ENCODE | ENCFF735UWS | ENCSR220ASC | GRCh38 | A549 | Homo sapiens | | |
| ATAC-seq | ENCODE | ENCFF333TAT | ENCSR868FGK | GRCh38 | K562 | Homo sapiens | | |
| ATAC-seq | ENCODE | ENCFF439EIO | ENCSR291GJU | GRCh38 | HepG2 | Homo sapiens | | |
| DNase-seq | ENCODE | ENCFF291OQF | ENCSR000CNN | mm10 | MEL | Mus musculus | | |
| DNase-seq | ENCODE | ENCFF855RCO | ENCSR000CMQ | mm10 | CH12.LX | Mus musculus | | |
| DNase-seq | ENCODE | ENCFF983UCL | ENCSR000EMU | GRCh38 | H1 | Homo sapiens | | |
| DNase-seq | ENCODE | ENCFF128ZVL | ENCSR000ELW | GRCh38 | A549 | Homo sapiens | | |
| DNase-seq | ENCODE | ENCFF274YGF | ENCSR000EKS | GRCh38 | K562 | Homo sapiens | | |
| DNase-seq | ENCODE | ENCFF080VVT | ENCSR000EOQ | GRCh38 | endothelial cell of umbilical vein (HUVEC) | Homo sapiens | | |
| DNase-seq | ENCODE | ENCFF218WIP | ENCSR000EPL | GRCh38 | NB4 | Homo sapiens | | |
| DNase-seq | ENCODE | ENCFF897NME | ENCSR149XIL | GRCh38 | HepG2 | Homo sapiens | | |
| ChIP-seq | ENCODE | ENCFF398DMC | ENCSR000ERI | mm10 | CH12.LX | Mus musculus | EP300-mouse | |
| ChIP-seq | ENCODE | ENCFF185UAQ | ENCSR000ETV | mm10 | MEL | Mus musculus | EP300-mouse | |
| ChIP-seq | ENCODE | ENCFF702XPO | ENCSR000EGE | GRCh38 | K562 | Homo sapiens | EP300-human | |
| ChIP-seq | ENCODE | ENCFF678QTH | ENCSR886OEO | GRCh38 | A549 | Homo sapiens | EP300-human | |
| ChIP-seq | ENCODE | ENCFF244VKF | ENCSR000BKK | GRCh38 | H1 | Homo sapiens | EP300-human | |
| ChIP-seq | ENCODE | ENCFF827LSX | ENCSR271XMW | GRCh38 | HepG2 | Homo sapiens | EP300-human | |
| ChIP-seq | ENCODE | ENCFF125SFM | ENCSR000CGL | mm10 | CH12.LX | Mus musculus | H3K9ac-mouse | |
| ChIP-seq | ENCODE | ENCFF392DCV | ENCSR000ALD | GRCh38 | endothelial cell of umbilical vein | Homo sapiens | H3K9ac-human | |
| ChIP-seq | ENCODE | ENCFF568IPQ | ENCSR000ALD | GRCh38 | endothelial cell of umbilical vein | Homo sapiens | H3K9ac-human | |
| ChIP-seq | ENCODE | ENCFF077LGZ | ENCSR000ALB | GRCh38 | endothelial cell of umbilical vein | Homo sapiens | H3K27ac-human | |
| ChIP-seq | ENCODE | ENCFF317QGQ | ENCSR000ANP | GRCh38 | H1 | Homo sapiens | H3K27ac-human | |
| ChIP-seq | ENCODE | ENCFF148UQI | ENCSR000EVZ | GRCh38 | K562 | Homo sapiens | H3K9ac-human | |
| ChIP-seq | ENCODE | ENCFF044QMC | ENCSR000AMD | GRCh38 | HepG2 | Homo sapiens | H3K9ac-human | |
| ChIP-seq | ENCODE | ENCFF392KDI | ENCSR000AMO | GRCh38 | HepG2 | Homo sapiens | H3K27ac-human | |
| ChIP-seq | ENCODE | ENCFF395EHX | ENCSR000CGJ | mm10 | CH12.LX | Mus musculus | H3K27ac-mouse | |
| ChIP-seq | ENCODE | ENCFF544LXB | ENCSR000AKP | GRCh38 | K562 | Homo sapiens | H3K27ac-human | |
| ChIP-seq | ENCODE | ENCFF747IZX | ENCSR778NQS | GRCh38 | A549 | Homo sapiens | H3K27ac-human | |
| ChIP-seq | ENCODE | ENCFF900QYB | ENCSR000CEU | mm10 | MEL | Mus musculus | H3K9ac-mouse | |
| ChIP-seq | ENCODE | ENCFF972YIT | ENCSR000CEV | mm10 | MEL | Mus musculus | H3K27ac-mouse | |
| ChIP-seq | ENCODE | ENCFF679LHF | ENCSR441UHO | GRCh38 | H1 | Homo sapiens | H3K9ac-human | |
| ChIP-seq | ENCODE | ENCFF354VWZ | ENCSR000EEM | GRCh38 | HepG2 | Homo sapiens | POLR2A-human | |
| ChIP-seq | ENCODE | ENCFF322DAE | ENCSR000BHN | GRCh38 | H1 | Homo sapiens | POLR2A-human | |
| ChIP-seq | ENCODE | ENCFF004KQA | ENCSR000EUC | mm10 | MEL | Mus musculus | POLR2A-mouse | |
| ChIP-seq | ENCODE | ENCFF269DMJ | ENCSR000ERQ | mm10 | CH12.LX | Mus musculus | POLR2A-mouse | |
| ChIP-seq | ENCODE | ENCFF730QNU | ENCSR000FAL | GRCh38 | NB4 | Homo sapiens | POLR2A-human | |
| ChIP-seq | ENCODE | ENCFF229KAY | ENCSR000BQB | GRCh38 | endothelial cell of umbilical vein | Homo sapiens | POLR2A-human | |
| ChIP-seq | ENCODE | ENCFF681MRC | ENCSR000DMZ | GRCh38 | A549 | Homo sapiens | POLR2A-human | |
| ChIP-seq | ENCODE | ENCFF355MNE | ENCSR388QZF | GRCh38 | K562 | Homo sapiens | POLR2A-human | |
| ChIA-PET | ENCODE | ENCFF364UNM | ENCSR857MYZ | GRCh38 | HepG2 | Homo sapiens | POLR2A-human | |
| ChIA-PET | ENCODE | ENCFF421KYP | ENCSR138NSW | GRCh38 | A549 | Homo sapiens | POLR2A-human | |
| ChIA-PET | ENCODE | ENCFF753NSM | ENCSR782EKZ | GRCh38 | H1 | Homo sapiens | POLR2A-human | |
| ChIA-PET | ENCODE | ENCFF308LJF | ENCSR080OMN | GRCh38 | endothelial cell of umbilical vein | Homo sapiens | POLR2A-human | |
| ChIA-PET | ENCODE | ENCFF511QFN | ENCSR880DSH | GRCh38 | K562 | Homo sapiens | POLR2A-human | |
| ChIP-seq | GEO | SRR8555222 | GSM3596839 | mm10 | primary naive mouse B-cells | Mus musculus | none(input control) | LPS (50 ug/ml) 0h |

**Table S1.** Cell lines and data sets used in the current study(cont'd)

| Assay type | Database | File accession ID | Experiment accession | Genome assembly | Biosample term name | Biosample organism | Experiment target | Biosample treatments |
|---|---|---|---|---|---|---|---|---|
| ChIP-seq | GEO | SRR8555223 | GSM3596840 | mm10 | primary naive mouse B-cells | Mus musculus | MYC-mouse | LPS (50 ug/ml) 0h |
| ChIP-seq | GEO | SRR8555224 | GSM3596841 | mm10 | primary naive mouse B-cells | Mus musculus | MYC-mouse | LPS (50 ug/ml) 2h |
| ChIP-seq | GEO | SRR8555225 | GSM3596842 | mm10 | primary naive mouse B-cells | Mus musculus | MYC-mouse | LPS (50 ug/ml) 4h |
| ChIP-seq | GEO | SRR8555226 | GSM3596843 | mm10 | primary naive mouse B-cells | Mus musculus | MYC-mouse | LPS (50 ug/ml) 8h |
| ChIP-seq | GEO | SRR5493408 | GSM2595123 | mm10 | 3T9 | Mus musculus | none(input control) | MYC-ER activation by 4-hydroxytamoxifen (400nM) |
| ChIP-seq | GEO | SRR5493409 | GSM2595124 | mm10 | 3T9 | Mus musculus | MYC-mouse | 0 min MYC-ER activation by 4-hydroxytamoxifen (400nM) |
| ChIP-seq | GEO | SRR5493410 | GSM2595125 | mm10 | 3T9 | Mus musculus | MYC-mouse | 10 min MYC-ER activation by 4-hydroxytamoxifen (400nM) |
| ChIP-seq | GEO | SRR5493411 | GSM2595126 | mm10 | 3T9 | Mus musculus | MYC-mouse | 20 min MYC-ER activation by 4-hydroxytamoxifen (400nM) |
| ChIP-seq | GEO | SRR5493412 | GSM2595127 | mm10 | 3T9 | Mus musculus | MYC-mouse | 30 min MYC-ER activation by 4-hydroxytamoxifen (400nM) |
| ChIP-seq | GEO | SRR5493413 | GSM2595128 | mm10 | 3T9 | Mus musculus | MYC-mouse | 2 h MYC-ER activation by 4-hydroxytamoxifen (400nM) |
| ChIP-seq | GEO | SRR5493414 | GSM2595129 | mm10 | 3T9 | Mus musculus | MYC-mouse | 4 h MYC-ER activation by 4-hydroxytamoxifen (400nM) |
| ChIP-seq | GEO | SRR3020030 | GSM1973344 | mm10 | normal liver | Mus musculus | MYC-mouse | |
| ChIP-seq | GEO | SRR3020031 | GSM1973345 | mm10 | normal liver | Mus musculus | MYC-mouse | |
| ChIP-seq | GEO | SRR3020032 | GSM1973346 | mm10 | normal liver | Mus musculus | MYC-mouse | |
| ChIP-seq | GEO | SRR3020033 | GSM1973347 | mm10 | normal liver | Mus musculus | MYC-mouse | |
| ChIP-seq | GEO | SRR3020034 | GSM1973348 | mm10 | tet-Myc liver tumor | Mus musculus | MYC-mouse | |
| ChIP-seq | GEO | SRR3020035 | GSM1973349 | mm10 | tet-Myc liver tumor | Mus musculus | MYC-mouse | |
| ChIP-seq | GEO | SRR3020036 | GSM1973350 | mm10 | tet-Myc liver tumor | Mus musculus | MYC-mouse | |
| ChIP-seq | GEO | SRR3020037 | GSM1973351 | mm10 | tet-Myc liver tumor | Mus musculus | MYC-mouse | |
| ChIP-seq | GEO | SRR3020038 | GSM1973352 | mm10 | tet-Myc liver tumor | Mus musculus | MYC-mouse | |
| ChIP-seq | GEO | SRR3020039 | GSM1973353 | mm10 | tet-Myc liver tumor | Mus musculus | MYC-mouse | |
| ChIP-seq | GEO | SRR3020040 | GSM1973354 | mm10 | tet-Myc liver tumor | Mus musculus | MYC-mouse | |
| ChIP-seq | GEO | SRR3020041 | GSM1973355 | mm10 | tet-Myc liver tumor | Mus musculus | MYC-mouse | |
| ChIP-seq | GEO | SRR3020042 | GSM1973356 | mm10 | tet-Myc liver tumor | Mus musculus | MYC-mouse | |
| ChIP-seq | GEO | SRR3020076 | GSM1973390 | mm10 | normal liver | Mus musculus | none(input control) | |
| ChIP-seq | GEO | SRR14907854 | GSM5399514 | GRCh38 | 22Rv1 | Homo sapiens | MYC-human | 1hr MYCi - 10 µM |
| ChIP-seq | GEO | SRR14907855 | GSM5399515 | GRCh38 | 22Rv1 | Homo sapiens | MYC-human | 1hr MYCi - 10 µM |
| ChIP-seq | GEO | SRR14907856 | GSM5399516 | GRCh38 | 22Rv1 | Homo sapiens | MYC-human | 1hr MYCi - 10 µM |
| ChIP-seq | GEO | SRR14907991 | GSM5399513 | GRCh38 | 22Rv1 | Homo sapiens | MYC-human | 1hr MYCi - 10 µM |
| ChIP-seq | GEO | SRR14907857 | GSM5399517 | GRCh38 | 22Rv1 | Homo sapiens | MAX-human | 1hr MYCi - 10 µM |
| ChIP-seq | GEO | SRR14907858 | GSM5399518 | GRCh38 | 22Rv1 | Homo sapiens | MAX-human | 1hr MYCi - 10 µM |
| ChIP-seq | GEO | SRR14907859 | GSM5399519 | GRCh38 | 22Rv1 | Homo sapiens | MAX-human | 1hr MYCi - 10 µM |
| ChIP-seq | GEO | SRR14907860 | GSM5399520 | GRCh38 | 22Rv1 | Homo sapiens | MAX-human | 1hr MYCi - 10 µM |
| ChIP-seq | GEO | SRR14907863 | GSM5399523 | GRCh38 | 22Rv1 | Homo sapiens | none(input control) | 1hr MYCi - 10 µM |
| ChIP-seq | GEO | SRR14907864 | GSM5399524 | GRCh38 | 22Rv1 | Homo sapiens | none(input control) | 1hr MYCi - 10 µM |
| ChIP-seq | GEO | SRR14907865 | GSM5399525 | GRCh38 | 22Rv1 | Homo sapiens | none(input control) | 1hr MYCi - 10 µM |
| ChIP-seq | GEO | SRR14907866 | GSM5399526 | GRCh38 | 22Rv1 | Homo sapiens | none(input control) | 1hr MYCi - 10 µM |
| ChIP-seq | GEO | SRR14907896 | GSM5399555 | GRCh38 | 22Rv1 | Homo sapiens | MYC-human | 24hr MYCi - 10 µM |
| ChIP-seq | GEO | SRR14907897 | GSM5399556 | GRCh38 | 22Rv1 | Homo sapiens | MYC-human | 24hr MYCi - 10 µM |
| ChIP-seq | GEO | SRR14907898 | GSM5399557 | GRCh38 | 22Rv1 | Homo sapiens | MYC-human | 24hr MYCi - 10 µM |
| ChIP-seq | GEO | SRR14907899 | GSM5399558 | GRCh38 | 22Rv1 | Homo sapiens | MYC-human | 24hr MYCi - 10 µM |
| ChIP-seq | GEO | SRR14907900 | GSM5399559 | GRCh38 | 22Rv1 | Homo sapiens | MAX-human | 24hr MYCi - 10 µM |
| ChIP-seq | GEO | SRR14907901 | GSM5399560 | GRCh38 | 22Rv1 | Homo sapiens | MAX-human | 24hr MYCi - 10 µM |
| ChIP-seq | GEO | SRR14907902 | GSM5399561 | GRCh38 | 22Rv1 | Homo sapiens | MAX-human | 24hr MYCi - 10 µM |
| ChIP-seq | GEO | SRR14907903 | GSM5399562 | GRCh38 | 22Rv1 | Homo sapiens | MAX-human | 24hr MYCi - 10 µM |
| ChIP-seq | GEO | SRR14907908 | GSM5399567 | GRCh38 | 22Rv1 | Homo sapiens | none(input control) | 24hr MYCi - 10 µM |
| ChIP-seq | GEO | SRR14907909 | GSM5399568 | GRCh38 | 22Rv1 | Homo sapiens | none(input control) | 24hr MYCi - 10 µM |
| ChIP-seq | GEO | SRR14907910 | GSM5399569 | GRCh38 | 22Rv1 | Homo sapiens | none(input control) | 24hr MYCi - 10 µM |
| ChIP-seq | GEO | SRR14907911 | GSM5399570 | GRCh38 | 22Rv1 | Homo sapiens | none(input control) | 24hr MYCi - 10 µM |
| ChIP-seq | GEO | SRR14907912 | GSM5399571 | GRCh38 | 22Rv1 | Homo sapiens | MYC-human | 48hr MYCi - 10 µM |
| ChIP-seq | GEO | SRR14907913 | GSM5399572 | GRCh38 | 22Rv1 | Homo sapiens | MYC-human | 48hr MYCi - 10 µM |
| ChIP-seq | GEO | SRR14907914 | GSM5399573 | GRCh38 | 22Rv1 | Homo sapiens | MYC-human | 48hr MYCi - 10 µM |
| ChIP-seq | GEO | SRR14907915 | GSM5399574 | GRCh38 | 22Rv1 | Homo sapiens | MYC-human | 48hr MYCi - 10 µM |
| ChIP-seq | GEO | SRR14907916 | GSM5399575 | GRCh38 | 22Rv1 | Homo sapiens | MAX-human | 48hr MYCi - 10 µM |
| ChIP-seq | GEO | SRR14907917 | GSM5399576 | GRCh38 | 22Rv1 | Homo sapiens | MAX-human | 48hr MYCi - 10 µM |
| ChIP-seq | GEO | SRR14907918 | GSM5399577 | GRCh38 | 22Rv1 | Homo sapiens | MAX-human | 48hr MYCi - 10 µM |
| ChIP-seq | GEO | SRR14907919 | GSM5399578 | GRCh38 | 22Rv1 | Homo sapiens | MAX-human | 48hr MYCi - 10 µM |
| ChIP-seq | GEO | SRR14907922 | GSM5399581 | GRCh38 | 22Rv1 | Homo sapiens | none(input control) | 48hr MYCi - 10 µM |
| ChIP-seq | GEO | SRR14907923 | GSM5399582 | GRCh38 | 22Rv1 | Homo sapiens | none(input control) | 48hr MYCi - 10 µM |
| ChIP-seq | GEO | SRR14907924 | GSM5399583 | GRCh38 | 22Rv1 | Homo sapiens | none(input control) | 48hr MYCi - 10 µM |
| ChIP-seq | GEO | SRR14907925 | GSM5399584 | GRCh38 | 22Rv1 | Homo sapiens | none(input control) | 48hr MYCi - 10 µM |
| ChIP-seq | GEO | SRR14907867 | GSM5399527 | GRCh38 | 22Rv1 | Homo sapiens | MYC-human | 4hr MYCi - 10 µM |
| ChIP-seq | GEO | SRR14907868 | GSM5399528 | GRCh38 | 22Rv1 | Homo sapiens | MYC-human | 4hr MYCi - 10 µM |

**Table S1.** Cell lines and data sets used in the current study(cont'd..)

| Assay type | Database | File accession ID | Experiment accession | Genome assembly | Biosample term name | Biosample organism | Experiment target | Biosample treatments |
|---|---|---|---|---|---|---|---|---|
| ChIP-seq | GEO | SRR14907869 | GSM5399529 | GRCh38 | 22Rv1 | Homo sapiens | MYC-human | 4hr MYCi - 10 µM |
| ChIP-seq | GEO | SRR14907870 | GSM5399530 | GRCh38 | 22Rv1 | Homo sapiens | MYC-human | 4hr MYCi - 10 µM |
| ChIP-seq | GEO | SRR14907871 | GSM5399531 | GRCh38 | 22Rv1 | Homo sapiens | MAX-human | 4hr MYCi - 10 µM |
| ChIP-seq | GEO | SRR14907872 | GSM5399532 | GRCh38 | 22Rv1 | Homo sapiens | MAX-human | 4hr MYCi - 10 µM |
| ChIP-seq | GEO | SRR14907873 | GSM5399533 | GRCh38 | 22Rv1 | Homo sapiens | MAX-human | 4hr MYCi - 10 µM |
| ChIP-seq | GEO | SRR14907874 | GSM5399534 | GRCh38 | 22Rv1 | Homo sapiens | MAX-human | 4hr MYCi - 10 µM |
| ChIP-seq | GEO | SRR14907877 | GSM5399537 | GRCh38 | 22Rv1 | Homo sapiens | none(input control) | 4hr MYCi - 10 µM |
| ChIP-seq | GEO | SRR14907878 | GSM5399538 | GRCh38 | 22Rv1 | Homo sapiens | none(input control) | 4hr MYCi - 10 µM |
| ChIP-seq | GEO | SRR14907879 | GSM5399539 | GRCh38 | 22Rv1 | Homo sapiens | none(input control) | 4hr MYCi - 10 µM |
| ChIP-seq | GEO | SRR14907881 | GSM5399540 | GRCh38 | 22Rv1 | Homo sapiens | none(input control) | 4hr MYCi - 10 µM |
| ChIP-seq | GEO | SRR14907882 | GSM5399541 | GRCh38 | 22Rv1 | Homo sapiens | MYC-human | 8hr MYCi - 10 µM |
| ChIP-seq | GEO | SRR14907883 | GSM5399542 | GRCh38 | 22Rv1 | Homo sapiens | MYC-human | 8hr MYCi - 10 µM |
| ChIP-seq | GEO | SRR14907884 | GSM5399543 | GRCh38 | 22Rv1 | Homo sapiens | MYC-human | 8hr MYCi - 10 µM |
| ChIP-seq | GEO | SRR14907885 | GSM5399544 | GRCh38 | 22Rv1 | Homo sapiens | MYC-human | 8hr MYCi - 10 µM |
| ChIP-seq | GEO | SRR14907886 | GSM5399545 | GRCh38 | 22Rv1 | Homo sapiens | MAX-human | 8hr MYCi - 10 µM |
| ChIP-seq | GEO | SRR14907887 | GSM5399546 | GRCh38 | 22Rv1 | Homo sapiens | MAX-human | 8hr MYCi - 10 µM |
| ChIP-seq | GEO | SRR14907888 | GSM5399547 | GRCh38 | 22Rv1 | Homo sapiens | MAX-human | 8hr MYCi - 10 µM |
| ChIP-seq | GEO | SRR14907889 | GSM5399548 | GRCh38 | 22Rv1 | Homo sapiens | MAX-human | 8hr MYCi - 10 µM |
| ChIP-seq | GEO | SRR14907892 | GSM5399551 | GRCh38 | 22Rv1 | Homo sapiens | none(input control) | 8hr MYCi - 10 µM |
| ChIP-seq | GEO | SRR14907893 | GSM5399552 | GRCh38 | 22Rv1 | Homo sapiens | none(input control) | 8hr MYCi - 10 µM |
| ChIP-seq | GEO | SRR14907894 | GSM5399553 | GRCh38 | 22Rv1 | Homo sapiens | none(input control) | 8hr MYCi - 10 µM |
| ChIP-seq | GEO | SRR14907895 | GSM5399554 | GRCh38 | 22Rv1 | Homo sapiens | none(input control) | 8hr MYCi - 10 µM |
| ChIP-seq | GEO | SRR14907974 | GSM5399497 | GRCh38 | 22Rv1 | Homo sapiens | MYC-human | DMSO (0.2%) |
| ChIP-seq | GEO | SRR14907975 | GSM5399498 | GRCh38 | 22Rv1 | Homo sapiens | MYC-human | DMSO (0.2%) |
| ChIP-seq | GEO | SRR14907976 | GSM5399499 | GRCh38 | 22Rv1 | Homo sapiens | MYC-human | DMSO (0.2%) |
| ChIP-seq | GEO | SRR14907977 | GSM5399500 | GRCh38 | 22Rv1 | Homo sapiens | MYC-human | DMSO (0.2%) |
| ChIP-seq | GEO | SRR14907978 | GSM5399501 | GRCh38 | 22Rv1 | Homo sapiens | MAX-human | DMSO (0.2%) |
| ChIP-seq | GEO | SRR14907979 | GSM5399502 | GRCh38 | 22Rv1 | Homo sapiens | MAX-human | DMSO (0.2%) |
| ChIP-seq | GEO | SRR14907980 | GSM5399503 | GRCh38 | 22Rv1 | Homo sapiens | MAX-human | DMSO (0.2%) |
| ChIP-seq | GEO | SRR14907981 | GSM5399504 | GRCh38 | 22Rv1 | Homo sapiens | MAX-human | DMSO (0.2%) |
| ChIP-seq | GEO | SRR14907986 | GSM5399509 | GRCh38 | 22Rv1 | Homo sapiens | none(input control) | DMSO (0.2%) |
| ChIP-seq | GEO | SRR14907988 | GSM5399510 | GRCh38 | 22Rv1 | Homo sapiens | none(input control) | DMSO (0.2%) |
| ChIP-seq | GEO | SRR14907989 | GSM5399511 | GRCh38 | 22Rv1 | Homo sapiens | none(input control) | DMSO (0.2%) |
| ChIP-seq | GEO | SRR14907990 | GSM5399512 | GRCh38 | 22Rv1 | Homo sapiens | none(input control) | DMSO (0.2%) |
| polyA plus RNA-seq | ENCODE | ENCFF472HFI | ENCSR000AEP | GRCh38 | K562 | Homo sapiens | | |
| polyA plus RNA-seq | ENCODE | ENCFF628SMT | ENCSR000AEP | GRCh38 | K562 | Homo sapiens | | |
| polyA plus RNA-seq | ENCODE | ENCFF486DGZ | ENCSR000AEQ | GRCh38 | K562 | Homo sapiens | | |
| polyA plus RNA-seq | ENCODE | ENCFF088LCK | ENCSR000AEQ | GRCh38 | K562 | Homo sapiens | | |
| polyA plus RNA-seq | ENCODE | ENCFF298KDC | ENCSR000CID | mm10 | MEL | Mus musculus | | |
| polyA plus RNA-seq | ENCODE | ENCFF952RYM | ENCSR000CID | mm10 | MEL | Mus musculus | | |
| polyA plus RNA-seq | ENCODE | ENCFF806WJZ | ENCSR937WIG | GRCh38 | A549 | Homo sapiens | | |
| polyA plus RNA-seq | ENCODE | ENCFF203NNS | ENCSR937WIG | GRCh38 | A549 | Homo sapiens | | |
| polyA plus RNA-seq | ENCODE | ENCFF715WBR | ENCSR937WIG | GRCh38 | A549 | Homo sapiens | | |
| polyA plus RNA-seq | ENCODE | ENCFF842EIL | ENCSR000AEO | GRCh38 | K562 | Homo sapiens | | |
| polyA plus RNA-seq | ENCODE | ENCFF322EYC | ENCSR000AEO | GRCh38 | K562 | Homo sapiens | | |
| polyA plus RNA-seq | ENCODE | ENCFF831QQF | ENCSR000CPE | GRCh38 | HepG2 | Homo sapiens | | |
| polyA plus RNA-seq | ENCODE | ENCFF168QKW | ENCSR000CPE | GRCh38 | HepG2 | Homo sapiens | | |
| polyA plus RNA-seq | ENCODE | ENCFF355TDA | ENCSR985KAT | GRCh38 | HepG2 | Homo sapiens | | |
| polyA plus RNA-seq | ENCODE | ENCFF073RKC | ENCSR985KAT | GRCh38 | HepG2 | Homo sapiens | | |
| polyA plus RNA-seq | ENCODE | ENCFF179CNW | ENCSR000EYO | GRCh38 | K562 | Homo sapiens | | |
| polyA plus RNA-seq | ENCODE | ENCFF679BCM | ENCSR000EYO | GRCh38 | K562 | Homo sapiens | | |
| polyA plus RNA-seq | ENCODE | ENCFF068NRZ | ENCSR000CPH | GRCh38 | K562 | Homo sapiens | | |
| polyA plus RNA-seq | ENCODE | ENCFF928YLB | ENCSR000CPH | GRCh38 | K562 | Homo sapiens | | |
| polyA plus RNA-seq | ENCODE | ENCFF321HCT | ENCSR000COU | GRCh38 | H1 | Homo sapiens | | |
| polyA plus RNA-seq | ENCODE | ENCFF736CCO | ENCSR000COU | GRCh38 | H1 | Homo sapiens | | |
| polyA plus RNA-seq | ENCODE | ENCFF443ZUL | ENCSR000CHM | mm10 | MEL | Mus musculus | | |
| polyA plus RNA-seq | ENCODE | ENCFF838AMX | ENCSR000CHM | mm10 | MEL | Mus musculus | | |
| polyA plus RNA-seq | ENCODE | ENCFF742CVV | ENCSR000AEM | GRCh38 | K562 | Homo sapiens | | |
| polyA plus RNA-seq | ENCODE | ENCFF222UVT | ENCSR000AEM | GRCh38 | K562 | Homo sapiens | | |
| polyA plus RNA-seq | ENCODE | ENCFF006IHP | ENCSR000BZU | GRCh38 | H1 | Homo sapiens | | |
| polyA plus RNA-seq | ENCODE | ENCFF779OCC | ENCSR000COZ | GRCh38 | endothelial cell of umbilical vein (HUVEC) | Homo sapiens | | |
| polyA plus RNA-seq | ENCODE | ENCFF770MAV | ENCSR000COZ | GRCh38 | endothelial cell of umbilical vein (HUVEC) | Homo sapiens | | |
| polyA plus RNA-seq | ENCODE | ENCFF855AKQ | ENCSR000CON | GRCh38 | A549 | Homo sapiens | | |
| polyA plus RNA-seq | ENCODE | ENCFF244DNJ | ENCSR000CON | GRCh38 | A549 | Homo sapiens | | |
| polyA plus RNA-seq | ENCODE | ENCFF082MCE | ENCSR000CHR | mm10 | CH12.LX | Mus musculus | | |
| polyA plus RNA-seq | ENCODE | ENCFF795QMY | ENCSR000CHR | mm10 | CH12.LX | Mus musculus | | |
| polyA plus RNA-seq | ENCODE | ENCFF881WXC | ENCSR000EYS | GRCh38 | endothelial cell of umbilical vein | Homo sapiens | | |
| polyA plus RNA-seq | ENCODE | ENCFF237XJC | ENCSR000EYS | GRCh38 | endothelial cell of umbilical vein | Homo sapiens | | |
| polyA plus RNA-seq | ENCODE | ENCFF233CZT | ENCSR632DQP | GRCh38 | A549 | Homo sapiens | | |
| polyA plus RNA-seq | ENCODE | ENCFF285BTU | ENCSR632DQP | GRCh38 | A549 | Homo sapiens | | |
| polyA plus RNA-seq | ENCODE | ENCFF904PCS | ENCSR632DQP | GRCh38 | A549 | Homo sapiens | | |
| polyA plus RNA-seq | ENCODE | ENCFF360GXF | ENCSR632DQP | GRCh38 | A549 | Homo sapiens | | |
| polyA plus RNA-seq | ENCODE | ENCFF190XYJ | ENCSR962TBJ | GRCh38 | H1 | Homo sapiens | | |
| polyA plus RNA-seq | ENCODE | ENCFF296UGP | ENCSR000EYP | GRCh38 | H1 | Homo sapiens | | |
| polyA plus RNA-seq | ENCODE | ENCFF247KYR | ENCSR000EYP | GRCh38 | H1 | Homo sapiens | | |

**Table S1.** Cell lines and data sets used in the current study(cont'd)

| Assay type | Database | File accession ID | Experiment accession | Genome assembly | Biosample term name | Biosample organism | Experiment target | Biosample treatments |
|---|---|---|---|---|---|---|---|---|
| polyA plus RNA-seq | ENCODE | ENCFF131NEX | ENCSR000EYP | GRCh38 | H1 | Homo sapiens | | |
| polyA plus RNA-seq | ENCODE | ENCFF562ECY | ENCSR000EYP | GRCh38 | H1 | Homo sapiens | | |
| polyA plus RNA-seq | ENCODE | ENCFF121UFS | ENCSR637VLS | GRCh38 | K562 | Homo sapiens | | |
| polyA plus RNA-seq | ENCODE | ENCFF938DAF | ENCSR637VLS | GRCh38 | K562 | Homo sapiens | | |
| polyA plus RNA-seq | ENCODE | ENCFF432RPO | ENCSR043RSE | GRCh38 | H1 | Homo sapiens | | |
| polyA plus RNA-seq | ENCODE | ENCFF695GMA | ENCSR000EYR | GRCh38 | HepG2 | Homo sapiens | | |
| polyA plus RNA-seq | ENCODE | ENCFF292KIL | ENCSR000EYR | GRCh38 | HepG2 | Homo sapiens | | |
| polyA plus RNA-seq | ENCODE | ENCFF806WAH | ENCSR643QIZ | GRCh38 | H1 | Homo sapiens | | |
| polyA plus RNA-seq | ENCODE | ENCFF582HOU | ENCSR643QIZ | GRCh38 | H1 | Homo sapiens | | |
| polyA plus RNA-seq | ENCODE | ENCFF243GKW | ENCSR670WQY | GRCh38 | H1 | Homo sapiens | | |
| polyA plus RNA-seq | ENCODE | ENCFF119KXQ | ENCSR561FEE | GRCh38 | HepG2 | Homo sapiens | | |
| polyA plus RNA-seq | ENCODE | ENCFF308OYV | ENCSR561FEE | GRCh38 | HepG2 | Homo sapiens | | |
| polyA plus RNA-seq | ENCODE | ENCFF416LVG | ENCSR545DKY | GRCh38 | K562 | Homo sapiens | | |
| polyA plus RNA-seq | ENCODE | ENCFF556ISR | ENCSR545DKY | GRCh38 | K562 | Homo sapiens | | |
| total RNA-seq | ENCODE | ENCFF263YGK | ENCSR000AJV | mm10 | CH12.LX | Mus musculus | | |
| total RNA-seq | ENCODE | ENCFF525WDQ | ENCSR000AJV | mm10 | CH12.LX | Mus musculus | | |
| total RNA-seq | ENCODE | ENCFF465NLB | ENCSR000AJV | mm10 | CH12.LX | Mus musculus | | |
| total RNA-seq | ENCODE | ENCFF445CEF | ENCSR000AJV | mm10 | CH12.LX | Mus musculus | | |
| total RNA-seq | ENCODE | ENCFF160NVZ | ENCSR414IGI | GRCh38 | A549 | Homo sapiens | | |
| total RNA-seq | ENCODE | ENCFF017ELJ | ENCSR414IGI | GRCh38 | A549 | Homo sapiens | | |
| total RNA-seq | ENCODE | ENCFF292XOH | ENCSR414IGI | GRCh38 | A549 | Homo sapiens | | |
| total RNA-seq | ENCODE | ENCFF823SJE | ENCSR414IGI | GRCh38 | A549 | Homo sapiens | | |
| total RNA-seq | ENCODE | ENCFF537IEO | ENCSR414IGI | GRCh38 | A549 | Homo sapiens | | |
| total RNA-seq | ENCODE | ENCFF507RJZ | ENCSR000CWD | mm10 | CH12.LX | Mus musculus | | |
| total RNA-seq | ENCODE | ENCFF469ZCH | ENCSR000CWD | mm10 | CH12.LX | Mus musculus | | |
| total RNA-seq | ENCODE | ENCFF203XTH | ENCSR000CWD | mm10 | CH12.LX | Mus musculus | | |
| total RNA-seq | ENCODE | ENCFF796PCV | ENCSR000CWD | mm10 | CH12.LX | Mus musculus | | |
| total RNA-seq | ENCODE | ENCFF472HPD | ENCSR000CWE | mm10 | MEL | Mus musculus | | |
| total RNA-seq | ENCODE | ENCFF933SWB | ENCSR000CWE | mm10 | MEL | Mus musculus | | |
| total RNA-seq | ENCODE | ENCFF475KFH | ENCSR000CWE | mm10 | MEL | Mus musculus | | |
| total RNA-seq | ENCODE | ENCFF082TDF | ENCSR000CWE | mm10 | MEL | Mus musculus | | |
| total RNA-seq | ENCODE | ENCFF067CVP | ENCSR181ZGR | GRCh38 | HepG2 | Homo sapiens | | |
| total RNA-seq | ENCODE | ENCFF343TKA | ENCSR181ZGR | GRCh38 | HepG2 | Homo sapiens | | |
| total RNA-seq | ENCODE | ENCFF286SDY | ENCSR181ZGR | GRCh38 | HepG2 | Homo sapiens | | |
| total RNA-seq | ENCODE | ENCFF140GVE | ENCSR181ZGR | GRCh38 | HepG2 | Homo sapiens | | |
| total RNA-seq | ENCODE | ENCFF379NOY | ENCSR895ZTB | GRCh38 | H1 | Homo sapiens | | |
| total RNA-seq | ENCODE | ENCFF675NTU | ENCSR895ZTB | GRCh38 | H1 | Homo sapiens | | |
| total RNA-seq | ENCODE | ENCFF948VKA | ENCSR993JMV | GRCh38 | endothelial cell of umbilical vein | Homo sapiens | | |
| total RNA-seq | ENCODE | ENCFF140CNM | ENCSR993JMV | GRCh38 | endothelial cell of umbilical vein | Homo sapiens | | |
| total RNA-seq | ENCODE | ENCFF932MJL | ENCSR792OIJ | GRCh38 | K562 | Homo sapiens | | |
| total RNA-seq | ENCODE | ENCFF170FKF | ENCSR792OIJ | GRCh38 | K562 | Homo sapiens | | |
| RNA-seq | GEO | SRR3020294 | GSM1973514 | mm10 | normal liver | Mus musculus | | |
| RNA-seq | GEO | SRR3020296 | GSM1973515 | mm10 | normal liver | Mus musculus | | |
| RNA-seq | GEO | SRR3020297 | GSM1973516 | mm10 | normal liver | Mus musculus | | |
| RNA-seq | GEO | SRR3020299 | GSM1973517 | mm10 | normal liver | Mus musculus | | |
| RNA-seq | GEO | SRR3020300 | GSM1973518 | mm10 | normal liver | Mus musculus | | |
| RNA-seq | GEO | SRR3020302 | GSM1973519 | mm10 | normal liver | Mus musculus | | |
| RNA-seq | GEO | SRR3020304 | GSM1973520 | mm10 | normal liver | Mus musculus | | |
| RNA-seq | GEO | SRR3020305 | GSM1973521 | mm10 | normal liver | Mus musculus | | |
| RNA-seq | GEO | SRR3020306 | GSM1973522 | mm10 | normal liver | Mus musculus | | |
| RNA-seq | GEO | SRR3020307 | GSM1973523 | mm10 | normal liver | Mus musculus | | |
| RNA-seq | GEO | SRR3020308 | GSM1973524 | mm10 | normal liver | Mus musculus | | |
| RNA-seq | GEO | SRR3020309 | GSM1973525 | mm10 | tet-Myc liver tumor | Mus musculus | | |
| RNA-seq | GEO | SRR3020310 | GSM1973526 | mm10 | tet-Myc liver tumor | Mus musculus | | |
| RNA-seq | GEO | SRR3020311 | GSM1973527 | mm10 | tet-Myc liver tumor | Mus musculus | | |
| RNA-seq | GEO | SRR3020312 | GSM1973528 | mm10 | tet-Myc liver tumor | Mus musculus | | |
| RNA-seq | GEO | SRR3020313 | GSM1973529 | mm10 | tet-Myc liver tumor | Mus musculus | | |
| RNA-seq | GEO | SRR3020314 | GSM1973530 | mm10 | tet-Myc liver tumor | Mus musculus | | |
| RNA-seq | GEO | SRR3020315 | GSM1973531 | mm10 | tet-Myc liver tumor | Mus musculus | | |
| RNA-seq | GEO | SRR3020316 | GSM1973532 | mm10 | tet-Myc liver tumor | Mus musculus | | |
| RNA-seq | GEO | SRR3020317 | GSM1973533 | mm10 | tet-Myc liver tumor | Mus musculus | | |
| RNA-seq | GEO | SRR3020318 | GSM1973534 | mm10 | tet-Myc liver tumor | Mus musculus | | |
| RNA-seq | GEO | SRR3020319 | GSM1973535 | mm10 | tet-Myc liver tumor | Mus musculus | | |
| RNA-seq | GEO | SRR3020320 | GSM1973536 | mm10 | tet-Myc liver tumor | Mus musculus | | |
| RNA-seq | GEO | SRR3020321 | GSM1973537 | mm10 | tet-Myc liver tumor | Mus musculus | | |
| RNA-seq | GEO | SRR3020322 | GSM1973538 | mm10 | tet-Myc liver tumor | Mus musculus | | |
| RNA-seq | GEO | SRR3020323 | GSM1973539 | mm10 | tet-Myc liver tumor | Mus musculus | | |
| RNA-seq | GEO | SRR3020324 | GSM1973540 | mm10 | tet-Myc liver tumor | Mus musculus | | |
| RNA-Seq | GEO | SRR8555186 | GSM3596803 | mm10 | Primary B-cells_WT | Mus musculus | | untreated |
| RNA-Seq | GEO | SRR8555187 | GSM3596804 | mm10 | Primary B-cells_WT | Mus musculus | | untreated |
| RNA-Seq | GEO | SRR8555188 | GSM3596805 | mm10 | Primary B-cells_WT | Mus musculus | | untreated |
| RNA-Seq | GEO | SRR8555189 | GSM3596806 | mm10 | Primary B-cells_WT | Mus musculus | | untreated |
| RNA-Seq | GEO | SRR8555190 | GSM3596807 | mm10 | Primary B-cells_WT | Mus musculus | | untreated |
| RNA-Seq | GEO | SRR8555191 | GSM3596808 | mm10 | Primary B-cells_WT | Mus musculus | | untreated |
| RNA-Seq | GEO | SRR8555192 | GSM3596809 | mm10 | Primary B-cells_MycKO | Mus musculus | | untreated |
| RNA-Seq | GEO | SRR8555193 | GSM3596810 | mm10 | Primary B-cells_MycKO | Mus musculus | | untreated |
| RNA-Seq | GEO | SRR8555194 | GSM3596811 | mm10 | Primary B-cells_MycKO | Mus musculus | | untreated |

#### **Table S1.** Cell lines and data sets used in the current study(cont'd)

| Assay type | Database | File accession ID | Experiment accession | Genome assembly | Biosample term name | Biosample organism | Experiment target | Biosample treatments |
|---|---|---|---|---|---|---|---|---|
| RNA-Seq | GEO | SRR8555195 | GSM3596812 | mm10 | Primary B-cells_MycKO | Mus musculus | | untreated |
| RNA-Seq | GEO | SRR8555196 | GSM3596813 | mm10 | Primary B-cells_MycKO | Mus musculus | | untreated |
| RNA-Seq | GEO | SRR8555197 | GSM3596814 | mm10 | Primary B-cells_MycKO | Mus musculus | | untreated |
| RNA-Seq | GEO | SRR8555198 | GSM3596815 | mm10 | Primary B-cells_WT | Mus musculus | | LPS (50 ug/ml) 2h |
| RNA-Seq | GEO | SRR8555199 | GSM3596816 | mm10 | Primary B-cells_WT | Mus musculus | | LPS (50 ug/ml) 2h |
| RNA-Seq | GEO | SRR8555200 | GSM3596817 | mm10 | Primary B-cells_WT | Mus musculus | | LPS (50 ug/ml) 2h |
| RNA-Seq | GEO | SRR8555201 | GSM3596818 | mm10 | Primary B-cells_WT | Mus musculus | | LPS (50 ug/ml) 2h |
| RNA-Seq | GEO | SRR8555202 | GSM3596819 | mm10 | Primary B-cells_MycKO | Mus musculus | | LPS (50 ug/ml) 2h |
| RNA-Seq | GEO | SRR8555203 | GSM3596820 | mm10 | Primary B-cells_MycKO | Mus musculus | | LPS (50 ug/ml) 2h |
| RNA-Seq | GEO | SRR8555204 | GSM3596821 | mm10 | Primary B-cells_MycKO | Mus musculus | | LPS (50 ug/ml) 2h |
| RNA-Seq | GEO | SRR8555205 | GSM3596822 | mm10 | Primary B-cells_MycKO | Mus musculus | | LPS (50 ug/ml) 2h |
| RNA-Seq | GEO | SRR8555206 | GSM3596823 | mm10 | Primary B-cells_WT | Mus musculus | | LPS (50 ug/ml) 4h |
| RNA-Seq | GEO | SRR8555207 | GSM3596824 | mm10 | Primary B-cells_WT | Mus musculus | | LPS (50 ug/ml) 4h |
| RNA-Seq | GEO | SRR8555208 | GSM3596825 | mm10 | Primary B-cells_WT | Mus musculus | | LPS (50 ug/ml) 4h |
| RNA-Seq | GEO | SRR8555209 | GSM3596826 | mm10 | Primary B-cells_WT | Mus musculus | | LPS (50 ug/ml) 4h |
| RNA-Seq | GEO | SRR8555210 | GSM3596827 | mm10 | Primary B-cells_MycKO | Mus musculus | | LPS (50 ug/ml) 4h |
| RNA-Seq | GEO | SRR8555211 | GSM3596828 | mm10 | Primary B-cells_MycKO | Mus musculus | | LPS (50 ug/ml) 4h |
| RNA-Seq | GEO | SRR8555212 | GSM3596829 | mm10 | Primary B-cells_MycKO | Mus musculus | | LPS (50 ug/ml) 4h |
| RNA-Seq | GEO | SRR8555213 | GSM3596830 | mm10 | Primary B-cells_MycKO | Mus musculus | | LPS (50 ug/ml) 4h |
| RNA-Seq | GEO | SRR8555214 | GSM3596831 | mm10 | Primary B-cells_WT | Mus musculus | | LPS (50 ug/ml) 8h |
| RNA-Seq | GEO | SRR8555215 | GSM3596832 | mm10 | Primary B-cells_WT | Mus musculus | | LPS (50 ug/ml) 8h |
| RNA-Seq | GEO | SRR8555216 | GSM3596833 | mm10 | Primary B-cells_WT | Mus musculus | | LPS (50 ug/ml) 8h |
| RNA-Seq | GEO | SRR8555217 | GSM3596834 | mm10 | Primary B-cells_WT | Mus musculus | | LPS (50 ug/ml) 8h |
| RNA-Seq | GEO | SRR8555218 | GSM3596835 | mm10 | Primary B-cells_MycKO | Mus musculus | | LPS (50 ug/ml) 8h |
| RNA-Seq | GEO | SRR8555219 | GSM3596836 | mm10 | Primary B-cells_MycKO | Mus musculus | | LPS (50 ug/ml) 8h |
| RNA-Seq | GEO | SRR8555220 | GSM3596837 | mm10 | Primary B-cells_MycKO | Mus musculus | | LPS (50 ug/ml) 8h |
| RNA-Seq | GEO | SRR8555221 | GSM3596838 | mm10 | Primary B-cells_MycKO | Mus musculus | | LPS (50 ug/ml) 8h |
| RNA-Seq | GEO | SRR5493432 | GSM2595068 | mm10 | 3T9 | Mus musculus | | 0 min MYC-ER activation by 4-hydroxytamoxifen (400nM) |
| RNA-Seq | GEO | SRR5493433 | GSM2595069 | mm10 | 3T9 | Mus musculus | | 10 min MYC-ER activation by 4-hydroxytamoxifen (400nM) |
| RNA-Seq | GEO | SRR5493434 | GSM2595070 | mm10 | 3T9 | Mus musculus | | 20 min MYC-ER activation by 4-hydroxytamoxifen (400nM) |
| RNA-Seq | GEO | SRR5493435 | GSM2595071 | mm10 | 3T9 | Mus musculus | | 30 min MYC-ER activation by 4-hydroxytamoxifen (400nM) |
| RNA-Seq | GEO | SRR5493436 | GSM2595072 | mm10 | 3T9 | Mus musculus | | 1h MYC-ER activation by 4-hydroxytamoxifen (400nM) |
| RNA-Seq | GEO | SRR5493437 | GSM2595073 | mm10 | 3T9 | Mus musculus | | 1.5h MYC-ER activation by 4-hydroxytamoxifen (400nM) |
| RNA-Seq | GEO | SRR5493438 | GSM2595074 | mm10 | 3T9 | Mus musculus | | 2 h MYC-ER activation by 4-hydroxytamoxifen (400nM) |
| RNA-Seq | GEO | SRR5493439 | GSM2595075 | mm10 | 3T9 | Mus musculus | | 4 h MYC-ER activation by 4-hydroxytamoxifen (400nM) |
| RNA-Seq | GEO | SRR5493440 | GSM2595076 | mm10 | 3T9 | Mus musculus | | 8 h MYC-ER activation by 4-hydroxytamoxifen (400nM) |
| RNA-Seq | GEO | SRR5493441 | GSM2595077 | mm10 | 3T9 | Mus musculus | | 12 h MYC-ER activation by 4-hydroxytamoxifen (400nM) |
| RNA-Seq | GEO | SRR5493442 | GSM2595078 | mm10 | 3T9 | Mus musculus | | 16 h MYC-ER activation by 4-hydroxytamoxifen (400nM) |
| RNA-Seq | GEO | SRR5493454 | GSM2595090 | mm10 | 3T9 | Mus musculus | | 0 min MYC-ER activation by 4-hydroxytamoxifen (400nM) |
| RNA-Seq | GEO | SRR5493455 | GSM2595091 | mm10 | 3T9 | Mus musculus | | 10 min MYC-ER activation by 4-hydroxytamoxifen (400nM) |
| RNA-Seq | GEO | SRR5493456 | GSM2595092 | mm10 | 3T9 | Mus musculus | | 20 min MYC-ER activation by 4-hydroxytamoxifen (400nM) |
| RNA-Seq | GEO | SRR5493457 | GSM2595093 | mm10 | 3T9 | Mus musculus | | 30 min MYC-ER activation by 4-hydroxytamoxifen (400nM) |
| RNA-Seq | GEO | SRR5493458 | GSM2595094 | mm10 | 3T9 | Mus musculus | | 1h MYC-ER activation by 4-hydroxytamoxifen (400nM) |
| RNA-Seq | GEO | SRR5493459 | GSM2595095 | mm10 | 3T9 | Mus musculus | | 1.5h MYC-ER activation by 4-hydroxytamoxifen (400nM) |
| RNA-Seq | GEO | SRR5493460 | GSM2595096 | mm10 | 3T9 | Mus musculus | | 2 h MYC-ER activation by 4-hydroxytamoxifen (400nM) |
| RNA-Seq | GEO | SRR5493461 | GSM2595097 | mm10 | 3T9 | Mus musculus | | 4 h MYC-ER activation by 4-hydroxytamoxifen (400nM) |
| RNA-Seq | GEO | SRR5493462 | GSM2595098 | mm10 | 3T9 | Mus musculus | | 8 h MYC-ER activation by 4-hydroxytamoxifen (400nM) |
| RNA-Seq | GEO | SRR5493463 | GSM2595099 | mm10 | 3T9 | Mus musculus | | 12 h MYC-ER activation by 4-hydroxytamoxifen (400nM) |
| RNA-Seq | GEO | SRR5493464 | GSM2595100 | mm10 | 3T9 | Mus musculus | | 16 h MYC-ER activation by 4-hydroxytamoxifen (400nM) |
| RNA-Seq | GEO | SRR5493476 | GSM2595112 | mm10 | 3T9 | Mus musculus | | 0 min MYC-ER activation by 4-hydroxytamoxifen (400nM) |
| RNA-Seq | GEO | SRR5493477 | GSM2595113 | mm10 | 3T9 | Mus musculus | | 10 min MYC-ER activation by 4-hydroxytamoxifen (400nM) |
| RNA-Seq | GEO | SRR5493478 | GSM2595114 | mm10 | 3T9 | Mus musculus | | 20 min MYC-ER activation by 4-hydroxytamoxifen (400nM) |
| RNA-Seq | GEO | SRR5493479 | GSM2595115 | mm10 | 3T9 | Mus musculus | | 30 min MYC-ER activation by 4-hydroxytamoxifen (400nM) |
| RNA-Seq | GEO | SRR5493480 | GSM2595116 | mm10 | 3T9 | Mus musculus | | 1h MYC-ER activation by 4-hydroxytamoxifen (400nM) |
| RNA-Seq | GEO | SRR5493481 | GSM2595117 | mm10 | 3T9 | Mus musculus | | 1.5h MYC-ER activation by 4-hydroxytamoxifen (400nM) |
| RNA-Seq | GEO | SRR5493482 | GSM2595118 | mm10 | 3T9 | Mus musculus | | 2 h MYC-ER activation by 4-hydroxytamoxifen (400nM) |
| RNA-Seq | GEO | SRR5493483 | GSM2595119 | mm10 | 3T9 | Mus musculus | | 4 h MYC-ER activation by 4-hydroxytamoxifen (400nM) |

**Table S1.** Cell lines and data sets used in the current study(cont'd)

| Assay type | Database | File accession ID | Experiment accession | Genome assembly | Biosample term name | Biosample organism | Experiment target | Biosample treatments |
|---|---|---|---|---|---|---|---|---|
| RNA-Seq | GEO | SRR5493484 | GSM2595120 | mm10 | 3T9 | Mus musculus | | 8 h MYC-ER activation by 4-hydroxytamoxifen (400nM) |
| RNA-Seq | GEO | SRR5493485 | GSM2595121 | mm10 | 3T9 | Mus musculus | | 12 h MYC-ER activation by 4-hydroxytamoxifen (400nM) |
| RNA-Seq | GEO | SRR5493486 | GSM2595122 | mm10 | 3T9 | Mus musculus | | 16 h MYC-ER activation by 4-hydroxytamoxifen (400nM) |
| ENCODE cCREs (encodeCcreCombined) | The UCSC Genome Browser database | encodeCcreCombined.bb | | GRCh38 | | Homo sapiens | | |
| remap2022_crm_macs2_hg38_v1_0.bed.gz | ReMAP2022 | https://remap.univ-amu.fr/storage/remap2022/hg38/MACS2/remap2022_crm_macs2_hg38_v1_0.bed.gz | | GRCh38 | | Homo sapiens | | |

**Table S2.** 190 genes associated with higher levels of TES-associated MYC+MAX binding and read-through transcription in HCCs vs. livers.

| gene_name | HCC | | | Liver | | | P value |
|---|---|---|---|---|---|---|---|
| | Gene_RPKM | DoG_RPKM | DoG_RPKM/Gene_RPKM | Gene_RPKM | DoG_RPKM | DoG_RPKM/Gene_RPKM | |
| 2410002F23Rik | 11.34078 | 0.01426 | 0.00123 | 1.53105 | 0.00000 | 0.00000 | 4.60E-03 |
| 9530068E07Rik | 16.66992 | 0.09146 | 0.00542 | 16.55367 | 0.04270 | 0.00275 | 1.72E-02 |
| Aars | 12.45808 | 0.18122 | 0.01526 | 27.31433 | 0.01128 | 0.00056 | 6.49E-06 |
| Acot4 | 7.93994 | 0.01391 | 0.00177 | 12.61325 | 0.00805 | 0.00033 | 9.17E-03 |
| Adk | 49.88304 | 0.39532 | 0.00877 | 114.30119 | 0.63074 | 0.00559 | 3.31E-02 |
| Afmid | 1.64394 | 0.08979 | 0.05635 | 20.98295 | 0.20805 | 0.00993 | 4.80E-05 |
| Akap13 | 2.89655 | 0.11263 | 0.03765 | 3.11615 | 0.04768 | 0.01667 | 3.26E-03 |
| Akt1 | 16.53958 | 0.10946 | 0.00674 | 5.80536 | 0.01966 | 0.00336 | 4.18E-02 |
| Alkbh6 | 4.22233 | 0.09751 | 0.02420 | 1.84113 | 0.01381 | 0.00778 | 5.98E-04 |
| Alkbh8 | 5.70713 | 0.17645 | 0.03061 | 1.26941 | 0.02068 | 0.01529 | 8.81E-03 |
| Ank | 1.91810 | 0.53853 | 0.28245 | 2.14291 | 0.41421 | 0.19531 | 4.91E-03 |
| Apoh | 65.95812 | 0.37588 | 0.00614 | 474.81591 | 0.66272 | 0.00141 | 8.10E-04 |
| Asb13 | 11.60138 | 0.05771 | 0.00495 | 9.43275 | 0.01455 | 0.00158 | 4.65E-03 |
| Aspg | 13.95176 | 0.08926 | 0.00656 | 33.03140 | 0.07748 | 0.00227 | 6.34E-04 |
| Atp5b | 308.36277 | 0.54225 | 0.00187 | 142.65346 | 0.12272 | 0.00089 | 9.26E-03 |
| Atp9b | 4.57492 | 0.13873 | 0.03073 | 3.97628 | 0.06805 | 0.01871 | 1.01E-03 |
| Atrn | 3.03428 | 0.11464 | 0.03838 | 8.15124 | 0.09634 | 0.01334 | 3.81E-04 |
| BC004004 | 8.80189 | 0.03951 | 0.00466 | 6.37933 | 0.00184 | 0.00027 | 3.20E-05 |
| Bcar3 | 3.52909 | 0.15310 | 0.04288 | 7.02469 | 0.09266 | 0.01282 | 1.43E-04 |
| C3 | 214.34285 | 0.16265 | 0.00079 | 1306.22436 | 0.26365 | 0.00021 | 3.02E-03 |
| C330022C24Rik | 5.41556 | 0.03617 | 0.00734 | 23.94696 | 0.02068 | 0.00095 | 1.93E-03 |
| Chd9 | 3.67687 | 0.24589 | 0.06760 | 3.51365 | 0.16014 | 0.04969 | 2.51E-02 |
| Chid1 | 7.37203 | 12.74452 | 1.73190 | 4.44363 | 6.41796 | 1.45363 | 6.15E-03 |
| Cluap1 | 6.19966 | 0.59598 | 0.09627 | 1.95130 | 0.11533 | 0.05899 | 3.67E-03 |
| Commd5 | 8.00275 | 0.37842 | 0.05099 | 4.21332 | 0.14459 | 0.03725 | 4.60E-02 |
| Coq8b | 2.31412 | 0.01259 | 0.00496 | 1.82505 | 0.00169 | 0.00084 | 4.94E-02 |
| Cox17 | 28.25153 | 0.01105 | 0.00042 | 9.81616 | 0.00000 | 0.00000 | 4.33E-02 |
| Cpb2 | 31.14404 | 0.52962 | 0.01805 | 57.64553 | 0.35572 | 0.00624 | 2.85E-06 |
| Crebbp | 4.19352 | 0.13751 | 0.03409 | 2.33239 | 0.03921 | 0.01723 | 8.74E-03 |
| Crebl2 | 1.87426 | 0.08986 | 0.04779 | 6.62354 | 0.07758 | 0.01154 | 2.91E-04 |
| Csf1r | 5.77080 | 3.61617 | 0.66203 | 3.84525 | 1.49284 | 0.44190 | 5.13E-03 |
| Ctdsp1 | 10.54117 | 0.12940 | 0.01232 | 15.37208 | 0.09037 | 0.00571 | 5.07E-04 |
| Cul4a | 20.57131 | 0.95883 | 0.04685 | 15.69491 | 0.46734 | 0.03282 | 2.62E-02 |
| Cxxc5 | 8.06895 | 0.21770 | 0.02656 | 5.72521 | 0.08850 | 0.01579 | 1.75E-02 |
| Dbt | 9.20462 | 0.04349 | 0.00483 | 13.41807 | 0.02117 | 0.00153 | 4.67E-03 |
| Dedd2 | 4.86472 | 0.07345 | 0.01552 | 3.10817 | 0.01042 | 0.00344 | 3.88E-04 |
| Dmpk | 1.32293 | 0.17604 | 0.14341 | 1.15845 | 0.09748 | 0.08097 | 8.59E-03 |
| Dnaja1 | 37.95137 | 16.98592 | 0.45325 | 46.72701 | 5.94671 | 0.13216 | 2.17E-11 |
| Dnajc14 | 4.22559 | 0.25865 | 0.06123 | 4.50589 | 0.06918 | 0.01635 | 5.56E-06 |
| Dnttip1 | 3.13726 | 0.23756 | 0.07471 | 1.37440 | 0.02180 | 0.01671 | 5.52E-08 |
| Dpm1 | 4.85415 | 1.43699 | 0.29933 | 4.65783 | 0.61344 | 0.13543 | 2.64E-07 |
| Dst | 3.42364 | 0.05714 | 0.01683 | 5.81082 | 0.01816 | 0.00365 | 2.79E-03 |
| Dyrk1a | 6.37261 | 0.21686 | 0.03375 | 2.55418 | 0.05166 | 0.02182 | 1.75E-02 |
| Eef1akmt4 | 7.09084 | 0.02480 | 0.00353 | 2.63222 | 0.00000 | 0.00000 | 1.82E-05 |

**Table S2.** 190 genes associated with higher levels of TES-associated MYC+MAX binding and read-through transcription in HCCs vs. livers(cont'd).

| gene_name | HCC | | | Liver | | | P value |
|---|---|---|---|---|---|---|---|
| | Gene_RPKM | DoG_RPKM | DoG_RPKM/Gene_RPKM | Gene_RPKM | DoG_RPKM | DoG_RPKM/Gene_RPKM | |
| Eef1b2 | 107.33352 | 1.36229 | 0.01287 | 16.52691 | 0.11634 | 0.00723 | 9.05E-04 |
| Eef2 | 263.48819 | 0.20858 | 0.00081 | 130.94803 | 0.04154 | 0.00033 | 2.01E-03 |
| Egfl7 | 13.05827 | 1.37701 | 0.10741 | 5.52801 | 0.06731 | 0.01292 | 1.65E-09 |
| Eif4a2 | 58.20868 | 15.18731 | 0.26087 | 35.78269 | 0.78064 | 0.02161 | 3.24E-17 |
| Eif5 | 58.69608 | 2.27125 | 0.03867 | 29.67156 | 0.73348 | 0.02488 | 1.07E-04 |
| Elk4 | 6.09982 | 0.34530 | 0.05666 | 6.46943 | 0.15819 | 0.02490 | 1.80E-06 |
| Eng | 2.78456 | 22.19393 | 8.61980 | 7.24490 | 20.10352 | 2.83641 | 6.58E-07 |
| Epb41 | 12.40302 | 1.16592 | 0.09631 | 10.90067 | 0.36703 | 0.03476 | 5.94E-04 |
| Erbb3 | 5.86840 | 0.05741 | 0.01001 | 14.82948 | 0.05558 | 0.00388 | 9.96E-03 |
| Etfb | 68.97742 | 0.18590 | 0.00287 | 115.15766 | 0.01786 | 0.00018 | 8.42E-04 |
| Evi5 | 11.80213 | 0.31051 | 0.02680 | 13.48888 | 0.19742 | 0.01511 | 2.38E-04 |
| Fads1 | 13.53383 | 0.07673 | 0.00578 | 103.77382 | 0.09996 | 0.00098 | 6.04E-03 |
| Farp1 | 1.10453 | 0.81479 | 0.78054 | 2.43819 | 0.45461 | 0.19314 | 1.23E-10 |
| Fbxl5 | 6.56279 | 0.56256 | 0.08741 | 2.93184 | 0.08717 | 0.03033 | 3.81E-08 |
| Foxn3 | 7.55283 | 0.04824 | 0.00650 | 6.34670 | 0.01978 | 0.00313 | 2.25E-02 |
| Gabarapl1 | 11.90741 | 0.41434 | 0.03571 | 60.42976 | 0.40099 | 0.00715 | 7.55E-06 |
| Gas5 | 117.35393 | 2.29247 | 0.02083 | 17.92110 | 0.23462 | 0.01387 | 3.21E-02 |
| Gcat | 56.06024 | 4.66023 | 0.08710 | 15.25562 | 0.02215 | 0.00141 | 1.56E-08 |
| Gnas | 75.01851 | 0.03658 | 0.00047 | 62.06142 | 0.01234 | 0.00019 | 2.46E-02 |
| Gng12 | 10.23798 | 0.36515 | 0.03543 | 4.20731 | 0.10122 | 0.02399 | 2.45E-02 |
| Golt1a | 3.82895 | 0.01121 | 0.00330 | 4.98266 | 0.00147 | 0.00027 | 3.44E-02 |
| Grb2 | 8.07495 | 0.09715 | 0.01182 | 4.24263 | 0.02837 | 0.00649 | 2.98E-02 |
| Grina | 8.62048 | 0.05344 | 0.00594 | 14.94226 | 0.02145 | 0.00156 | 6.60E-04 |
| H13 | 18.63147 | 0.13058 | 0.00717 | 22.45215 | 0.06245 | 0.00321 | 8.26E-03 |
| Herc4 | 5.73196 | 0.21062 | 0.03722 | 5.07393 | 0.13059 | 0.02522 | 3.13E-02 |
| Hmgcs2 | 7.21698 | 0.25264 | 0.03700 | 76.77754 | 0.46113 | 0.00642 | 2.28E-06 |
| Hook3 | 3.17220 | 0.10103 | 0.03153 | 4.50197 | 0.07941 | 0.01751 | 1.76E-02 |
| Hsd11b1 | 4.01656 | 0.02421 | 0.00687 | 168.72190 | 0.14974 | 0.00086 | 3.44E-03 |
| Hsd17b2 | 9.78746 | 0.06320 | 0.00660 | 41.05146 | 0.12799 | 0.00292 | 3.32E-03 |
| Ifnar2 | 2.45068 | 0.11209 | 0.04547 | 1.08456 | 0.02361 | 0.02459 | 4.32E-02 |
| Ip6k1 | 5.10926 | 0.01592 | 0.00314 | 6.70319 | 0.00545 | 0.00083 | 1.77E-02 |
| Irf6 | 2.33426 | 0.13290 | 0.06128 | 5.64007 | 0.04310 | 0.00771 | 1.34E-05 |
| Jam2 | 1.27642 | 0.45734 | 0.37189 | 1.27520 | 0.23896 | 0.19733 | 5.56E-07 |
| Jmjd8 | 18.86097 | 2.25497 | 0.12040 | 13.99545 | 1.25238 | 0.08701 | 2.59E-04 |
| Kansl3 | 3.98085 | 0.10719 | 0.02743 | 2.50058 | 0.04090 | 0.01595 | 2.49E-02 |
| Kif13b | 3.96629 | 0.28026 | 0.07017 | 5.94631 | 0.29351 | 0.05037 | 3.47E-02 |
| Klf9 | 6.33103 | 0.42799 | 0.06843 | 9.89090 | 0.28537 | 0.02948 | 3.64E-07 |
| Lipc | 12.17822 | 0.14504 | 0.01278 | 39.45775 | 0.03881 | 0.00097 | 1.42E-04 |
| Litaf | 17.44704 | 0.27388 | 0.01576 | 12.08337 | 0.08777 | 0.00746 | 6.17E-05 |
| Lrba | 2.60069 | 0.08241 | 0.03225 | 1.68085 | 0.02382 | 0.01485 | 9.12E-03 |
| Lrrc8a | 3.75985 | 0.26126 | 0.07116 | 4.48840 | 0.13945 | 0.03127 | 2.47E-05 |
| Malat1 | 239.30118 | 0.85539 | 0.00365 | 450.23848 | 0.29439 | 0.00066 | 4.27E-08 |
| Matr3 | 32.40263 | 1.14871 | 0.03515 | 7.52704 | 0.16688 | 0.02237 | 2.52E-03 |
| Mbnl1 | 18.78336 | 0.74499 | 0.03964 | 12.69409 | 0.33937 | 0.02569 | 4.39E-03 |

**Table S2.** 190 genes associated with higher levels of TES-associated MYC+MAX binding and read-through transcription in HCCs vs. livers(cont'd)).

| gene_name | HCC | | | Liver | | | P value |
|---|---|---|---|---|---|---|---|
| | Gene_RPKM | DoG_RPKM | DoG_RPKM/Gene_RPKM | Gene_RPKM | DoG_RPKM | DoG_RPKM/Gene_RPKM | |
| Mccc1 | 2.78989 | 0.66021 | 0.23775 | 3.45600 | 0.47701 | 0.13678 | 1.18E-04 |
| Mgll | 13.41916 | 0.08420 | 0.00633 | 31.22481 | 0.06074 | 0.00200 | 2.39E-04 |
| Mmd | 3.02775 | 0.02138 | 0.00683 | 11.59133 | 0.03091 | 0.00266 | 2.59E-02 |
| Mrpl15 | 3.59307 | 0.83807 | 0.23513 | 1.10338 | 0.12251 | 0.11614 | 3.82E-04 |
| Msl1 | 4.62548 | 0.10253 | 0.02159 | 3.13617 | 0.02948 | 0.00987 | 8.52E-03 |
| Mtor | 5.57465 | 0.02817 | 0.00509 | 4.10461 | 0.00720 | 0.00156 | 1.69E-02 |
| Ndufb9 | 84.82118 | 0.32195 | 0.00413 | 108.30192 | 0.21495 | 0.00214 | 1.36E-02 |
| Nek7 | 9.81215 | 0.18031 | 0.01830 | 8.61428 | 0.08129 | 0.00932 | 1.44E-02 |
| Nfib | 4.49735 | 0.34477 | 0.07773 | 9.66122 | 0.26678 | 0.02808 | 2.79E-09 |
| Nipsnap1 | 89.87719 | 0.52639 | 0.00595 | 53.24608 | 0.07275 | 0.00137 | 4.63E-07 |
| Nr3c2 | 1.40740 | 0.22085 | 0.16221 | 1.90801 | 0.07397 | 0.04326 | 7.58E-07 |
| Nsmf | 2.68291 | 0.14348 | 0.05482 | 14.08314 | 0.19722 | 0.01420 | 2.48E-06 |
| Nucks1 | 34.49483 | 0.36951 | 0.01071 | 10.74348 | 0.07056 | 0.00684 | 4.73E-03 |
| Numb | 2.30352 | 1.32581 | 0.57393 | 2.62278 | 0.25995 | 0.10042 | 5.44E-10 |
| Oaz1 | 68.36889 | 0.19938 | 0.00300 | 69.67492 | 0.03563 | 0.00052 | 1.77E-06 |
| Oaz1-ps | 50.00989 | 0.38097 | 0.00823 | 53.12362 | 0.10726 | 0.00235 | 2.12E-04 |
| Pcsk6 | 5.07658 | 0.62130 | 0.12419 | 12.94952 | 0.09845 | 0.00829 | 1.38E-09 |
| Pex6 | 9.37149 | 2.14509 | 0.23413 | 11.67859 | 1.32602 | 0.13515 | 1.20E-04 |
| Pfdn2 | 10.74426 | 0.23371 | 0.02203 | 3.94839 | 0.05461 | 0.01360 | 1.43E-02 |
| Phkb | 3.99376 | 0.40147 | 0.10088 | 3.46376 | 0.26239 | 0.07712 | 4.98E-03 |
| Pick1 | 6.35381 | 0.02885 | 0.00466 | 1.88852 | 0.00000 | 0.00000 | 2.22E-03 |
| Pik3c2a | 14.05468 | 0.95924 | 0.06835 | 5.52838 | 0.16583 | 0.03036 | 7.95E-05 |
| Pik3c2g | 1.25602 | 0.03832 | 0.03939 | 1.78056 | 0.02488 | 0.01348 | 3.08E-02 |
| Pknox1 | 2.56655 | 0.10710 | 0.04366 | 1.69561 | 0.03590 | 0.02060 | 1.84E-02 |
| Pnkd | 4.30315 | 0.03632 | 0.00776 | 7.49645 | 0.01116 | 0.00138 | 6.90E-04 |
| Pnpla7 | 4.59443 | 0.11631 | 0.02528 | 38.07515 | 0.17683 | 0.00489 | 1.21E-06 |
| Ppp1r21 | 3.64693 | 0.19434 | 0.05279 | 2.16074 | 0.04693 | 0.02067 | 3.48E-05 |
| Prr14 | 2.89118 | 0.11486 | 0.03902 | 4.52120 | 0.10021 | 0.02341 | 2.20E-02 |
| Ptbp1 | 67.99520 | 0.36709 | 0.00537 | 11.42549 | 0.02239 | 0.00192 | 2.08E-05 |
| Ptprf | 10.99600 | 0.67180 | 0.06108 | 18.08228 | 0.62899 | 0.03515 | 1.96E-07 |
| Rabggtb | 25.31230 | 1.40123 | 0.05613 | 6.02275 | 0.13963 | 0.02355 | 2.80E-06 |
| Ramp2 | 16.76655 | 0.16949 | 0.01019 | 12.79984 | 0.04335 | 0.00331 | 3.45E-04 |
| Rara | 3.33304 | 0.02708 | 0.00774 | 3.06419 | 0.00079 | 0.00025 | 1.69E-03 |
| Rc3h1 | 6.12920 | 0.35182 | 0.05679 | 2.95602 | 0.10572 | 0.03390 | 2.76E-03 |
| Repin1 | 4.91507 | 0.09606 | 0.01979 | 5.67922 | 0.01218 | 0.00208 | 9.12E-09 |
| Rhou | 2.65160 | 0.07441 | 0.02784 | 10.88760 | 0.06135 | 0.00704 | 8.72E-06 |
| Rit1 | 4.22707 | 0.15809 | 0.03819 | 2.89265 | 0.07743 | 0.02661 | 3.77E-02 |
| Rnf130 | 2.96437 | 0.07620 | 0.02647 | 8.95353 | 0.08341 | 0.00925 | 2.01E-03 |
| Rnf14 | 5.63259 | 0.14671 | 0.02640 | 7.30376 | 0.09435 | 0.01320 | 7.28E-04 |
| Rpl17 | 341.86199 | 1.92865 | 0.00577 | 68.48556 | 0.26236 | 0.00389 | 8.60E-03 |
| Rpl21 | 115.56349 | 0.86557 | 0.00810 | 22.87647 | 0.11202 | 0.00513 | 2.83E-02 |
| Rpl27a | 290.31906 | 2.07393 | 0.00744 | 57.03651 | 0.27439 | 0.00479 | 6.44E-04 |
| Rpl37 | 82.75293 | 2.29689 | 0.03039 | 15.86345 | 0.29958 | 0.02032 | 2.18E-02 |
| Rpl4 | 547.56298 | 0.70036 | 0.00132 | 94.61602 | 0.03237 | 0.00035 | 9.64E-07 |

**Table S2.** 190 genes associated with higher levels of TES-associated MYC+MAX binding and read-through transcription in HCCs vs. livers(cont'd).

| gene_name | HCC | | | Liver | | | P value |
|---|---|---|---|---|---|---|---|
| | Gene_RPKM | DoG_RPKM | DoG_RPKM/Gene_RPKM | Gene_RPKM | DoG_RPKM | DoG_RPKM/Gene_RPKM | |
| Rpl6 | 159.99727 | 0.99485 | 0.00676 | 32.42205 | 0.09607 | 0.00306 | 4.53E-03 |
| Rps20 | 232.91263 | 2.00933 | 0.00902 | 36.81532 | 0.14260 | 0.00402 | 6.87E-05 |
| Rpsa | 299.98263 | 2.07477 | 0.00735 | 52.01091 | 0.16088 | 0.00353 | 7.20E-04 |
| Rusf1 | 6.58994 | 0.23601 | 0.03614 | 2.89391 | 0.05266 | 0.01879 | 8.19E-04 |
| Ryk | 12.97168 | 0.44937 | 0.03535 | 4.22666 | 0.07271 | 0.01782 | 8.65E-04 |
| Sap18 | 3.08975 | 4.14444 | 1.34454 | 1.86405 | 0.10423 | 0.05812 | 1.57E-13 |
| Serpinc1 | 52.59892 | 3.45409 | 0.06733 | 169.05848 | 2.25578 | 0.01346 | 1.40E-09 |
| Sft2d1 | 5.47567 | 0.00773 | 0.00138 | 4.16691 | 0.00000 | 0.00000 | 1.86E-02 |
| Sgms1 | 4.93866 | 0.15574 | 0.03141 | 1.88629 | 0.03267 | 0.01812 | 3.48E-02 |
| Shb | 1.09890 | 0.04153 | 0.03557 | 3.78296 | 0.05010 | 0.01423 | 2.45E-02 |
| Sirt7 | 3.91522 | 0.11369 | 0.02884 | 6.66898 | 0.12695 | 0.01808 | 1.73E-02 |
| Slc16a1 | 69.20914 | 0.30102 | 0.00451 | 22.53791 | 0.01485 | 0.00074 | 2.93E-08 |
| Slc25a16 | 2.98584 | 0.07428 | 0.02503 | 8.10769 | 0.02770 | 0.00356 | 2.02E-07 |
| Slc25a47 | 10.09411 | 0.21148 | 0.02354 | 76.27969 | 0.45370 | 0.00581 | 3.97E-05 |
| Slc35e2 | 4.64409 | 0.48176 | 0.10549 | 12.69505 | 0.24359 | 0.02025 | 4.56E-09 |
| Slc6a6 | 15.13141 | 0.14463 | 0.00946 | 10.86325 | 0.04475 | 0.00388 | 5.92E-05 |
| Smap2 | 4.30959 | 0.22924 | 0.05299 | 3.58705 | 0.12563 | 0.03596 | 1.33E-02 |
| Smo | 5.68280 | 0.14750 | 0.02590 | 1.94513 | 0.02003 | 0.01096 | 3.20E-03 |
| Smurf1 | 1.10130 | 0.03195 | 0.03122 | 1.44364 | 0.01544 | 0.01210 | 1.38E-02 |
| Snhg12 | 27.24115 | 3.63577 | 0.13356 | 15.14413 | 0.40515 | 0.02927 | 2.03E-13 |
| Snora16a | 27.44810 | 2.47587 | 0.10015 | 9.24959 | 0.22397 | 0.03005 | 1.13E-05 |
| Snora20 | 2.69603 | 2.50515 | 0.96650 | 8.78256 | 1.03801 | 0.12886 | 1.27E-09 |
| Snora3 | 8.16171 | 3.25818 | 0.41059 | 2.00327 | 0.45206 | 0.25707 | 2.89E-03 |
| Snora44 | 26.90402 | 2.43713 | 0.10762 | 5.25847 | 0.18551 | 0.04038 | 1.71E-03 |
| Snora61 | 112.60862 | 2.47001 | 0.02408 | 24.14663 | 0.20694 | 0.01037 | 2.31E-03 |
| Snord22 | 688.30769 | 2.39465 | 0.00476 | 368.46463 | 0.06908 | 0.00024 | 3.01E-03 |
| Srd5a1 | 2.98289 | 0.04570 | 0.01572 | 14.29234 | 0.05513 | 0.00373 | 2.39E-04 |
| Srebf1 | 11.87667 | 2.22253 | 0.18983 | 50.59409 | 0.57400 | 0.01370 | 7.13E-10 |
| Srf | 3.89285 | 0.92646 | 0.24268 | 1.37170 | 0.21125 | 0.14842 | 6.57E-03 |
| Ssh2 | 3.15768 | 0.08937 | 0.02855 | 3.02354 | 0.05123 | 0.01726 | 1.42E-02 |
| Stat1 | 4.52154 | 0.24369 | 0.05411 | 4.66229 | 0.14029 | 0.03095 | 1.63E-04 |
| Taok3 | 2.48144 | 0.06538 | 0.02731 | 1.97975 | 0.02714 | 0.01513 | 1.19E-02 |
| Tapbp | 4.38145 | 0.23626 | 0.05621 | 7.55369 | 0.10315 | 0.01344 | 6.68E-06 |
| Tapt1 | 7.77002 | 0.30905 | 0.03989 | 11.72817 | 0.30574 | 0.02634 | 2.40E-03 |
| Thtpa | 3.29474 | 3.11874 | 0.97613 | 3.44360 | 0.63914 | 0.20642 | 1.98E-09 |
| Tlcd1 | 14.71093 | 4.76327 | 0.32753 | 7.23593 | 1.65309 | 0.23545 | 9.65E-04 |
| Tmem123 | 9.29884 | 0.44816 | 0.04934 | 5.93812 | 0.02881 | 0.00514 | 2.28E-07 |
| Tmem184b | 2.94674 | 0.21777 | 0.07342 | 2.13348 | 0.10680 | 0.04903 | 1.62E-02 |
| Tmem38b | 5.75361 | 0.07184 | 0.01263 | 15.11911 | 0.07983 | 0.00577 | 1.68E-03 |
| Tmem62 | 1.22568 | 0.01106 | 0.00875 | 2.33360 | 0.00385 | 0.00209 | 1.96E-02 |
| Tnpo2 | 7.84798 | 0.34012 | 0.04403 | 2.84040 | 0.04374 | 0.01913 | 2.26E-03 |
| Tpt1 | 399.02910 | 1.26237 | 0.00323 | 171.18316 | 0.25973 | 0.00159 | 9.11E-06 |
| Traf7 | 5.01921 | 0.68765 | 0.13724 | 1.06676 | 0.05643 | 0.05802 | 2.69E-05 |
| Tymp | 12.93516 | 17.19967 | 1.32454 | 24.56404 | 4.72316 | 0.21049 | 9.72E-16 |

**Table S2.** 190 genes associated with higher levels of TES-associated MYC+MAX binding and read-through transcription in HCCs vs. livers(cont'd).

| gene_name | HCC | | | Liver | | | P value |
|---|---|---|---|---|---|---|---|
| | Gene_RPKM | DoG_RPKM | DoG_RPKM/Gene_RPKM | Gene_RPKM | DoG_RPKM | DoG_RPKM/Gene_RPKM | |
| Uba7 | 1.45551 | 0.01877 | 0.01524 | 1.55472 | 0.00244 | 0.00154 | 4.51E-02 |
| Ubqln1 | 19.62581 | 0.36670 | 0.01831 | 13.86032 | 0.09122 | 0.00658 | 1.07E-05 |
| Ugt1a10 | 12.09358 | 0.73280 | 0.06371 | 168.72442 | 1.09888 | 0.00694 | 3.06E-08 |
| Ugt1a6a | 12.09358 | 0.73280 | 0.06371 | 168.72442 | 1.09888 | 0.00694 | 3.06E-08 |
| Ugt1a7c | 12.09358 | 0.73280 | 0.06371 | 168.72442 | 1.09888 | 0.00694 | 3.06E-08 |
| Uso1 | 4.87062 | 0.07419 | 0.01566 | 8.28451 | 0.05000 | 0.00648 | 2.27E-02 |
| Vps13b | 5.33378 | 0.99298 | 0.18464 | 3.42649 | 0.25932 | 0.07595 | 4.57E-07 |
| Vps35l | 4.80230 | 13.80961 | 2.90414 | 1.85689 | 2.83377 | 1.55445 | 3.32E-10 |
| Whamm | 2.89285 | 0.28749 | 0.10000 | 11.92157 | 0.28676 | 0.02388 | 2.43E-08 |
| Zfand3 | 4.94925 | 0.23571 | 0.04804 | 7.86967 | 0.16858 | 0.02103 | 9.48E-05 |
| Zfas1 | 43.26804 | 1.72950 | 0.04148 | 2.97412 | 0.03629 | 0.01174 | 4.48E-05 |
| Zfp444 | 4.31142 | 0.02822 | 0.00697 | 1.79141 | 0.00460 | 0.00251 | 4.57E-02 |
| Zfp524 | 1.95369 | 0.19389 | 0.09868 | 1.81303 | 0.09346 | 0.05310 | 5.56E-03 |
| Zfp710 | 3.24251 | 1.51147 | 0.47205 | 1.15344 | 0.21484 | 0.18745 | 3.38E-08 |

**Table S3.** Oligonucleotides used to create pDG458 gRNA constructs for CRISPR-Cas9 targeting of TES-associated E boxes and PCR primers for amplifying regions flanking the targeted E boxes.

| Crispr-Cas9 targeting site | Purpose | oligo Name | Sequance |
|---|---|---|---|
| Rps19_TES | for pDG458 construct | Rps19_Fw_P2_gR | accgCTAAGGCTTTGAGAATCACAgt |
| | | Rps19_Fw_P2_gR_rc | taaaacTGTGATTCTCAAAGCCTTAG |
| | amplify E box flanking region | Rps19_TES_fwd | AGTGCGGTTGAGCAGATTTA |
| | | Rps19_TES_rev | CAGTTGGCAGTTCCTGAAGA |
| Tspo/Ttll12_TES | for pDG458 construct | Tspo_fw_P1_gR | caccgTGCCCACCTTGAGCACGTGC |
| | | Tspo_fw_P1_gR_rc | aaacGCACGTGCTCAAGGTGGGCAc |
| | amplify E box flanking region | Tspo_TES_fw | GGTCTCTGGCTTGTGCTTAT |
| | | Tspo_TES_rev | CAAGACAGCTCACCGAGTAAA |
| Git1/Trp53i13_TES | for pDG458 construct | Git1_rv_P1_gR | caccgCTCTGCGACCGCTGCCACGT |
| | | Git1_rv_P1_gR_rc | aaacACGTGGCAGCGGTCGCAGAGc |
| | amplify E box flanking region | Git1_TES_fw | TTCCTTGCCCTTGACTTCTC |
| | | Git1_TES_rev | CCATGTCCTCTCTCTGCAATC |
| Hsp90aa1_TES | for pDG458 construct | Hsp90aa1_fw_P1_gR | caccgCATGAATCAGGCACGTGCCT |
| | | Hsp90aa1_fw_P1_gR_rc | aaacAGGCACGTGCCTGATTCATGc |
| | amplify E box flanking region | Hsp90aa1_TES_fw | CCCGAAACAAGTGCTTTGATAC |
| | | Hsp90aa1_TES_rev | CTACTTTCCTTCCCTCCCTTTG |
| Dyrk3_TES | for pDG458 construct | Dyrk3_fw_P1_gR | caccgAAAAACTGAACTGGGTCACG |
| | | Dyrk3_fw_P1_gR_rc | aaacCGTGACCCAGTTCAGTTTTTc |
| | amplify E box flanking region | Dyrk3_TES_fw | CAGGGCTTCCACACTGATAA |
| | | Dyrk3_TES_rev | TGACTCCTGTGGGCATTAAG |

**Table S4.** qRT-PCR primers used to quantify gene expression and DoG expression in WT and KO fibroblasts (Figure 7B-7D).

| Name of Gene | NCBI Gene ID | Assay | Reverse transcription primer | RT-PCR primers |
|---|---|---|---|---|
| Tspo | 12257 | Total gene expression | Random hexamers | Fwd: 5 '-CGCTTGCTGTACCCTTACC-3'<br>Rev: 5'-CCAGAGTTATCACGCCATACAT-3' |
| | | Expression of DoG | TES_RT: 5'-CAAGACAGCTCACCGAGTAAA-3' | Fwd: 5 '-CGCTTGCTGTACCCTTACC-3'<br>Rev: 5'-CCAGAGTTATCACGCCATACAT-3' |
| Ttll12 | 223723 | Total gene expression | Random hexamers | Fwd: 5'-CCAAATCCTGGAGGTGAACTT-3'<br>Rev: 5'-TCAGTCTCGTCCAGAAACAAAG-3' |
| | | Expression of DoG | TES_RT:5'-GGTCTCTGGCTTGTGCTTAT-3' | Fwd: 5'-CCAAATCCTGGAGGTGAACTT-3'<br>Rev: 5'-TCAGTCTCGTCCAGAAACAAAG-3' |
| Git1 | 216963 | Total gene expression | Random hexamers | Fwd: 5'-CGGCTTCAGAGCGAGTG-3'<br>Rev: 5'-CAGCCTTGGCGATGTCATA-3' |
| | | Expression of DoG | TES_RT: 5'-TTCCTTGCCCTTGACTTCTC-3' | Fwd: 5'-CGGCTTCAGAGCGAGTG-3'<br>Rev: 5'-CAGCCTTGGCGATGTCATA-3' |
| Trp53i13 | 216964 | Total gene expression | Random hexamers | Fwd: 5'-CCTCGATCAGTGTTGTGAAGAG-3'<br>Rev: 5'-CTGTGAAGAGGCCAAGGAAA-3' |
| | | Expression of DoG | TES_RT: 5'-CCATGTCCTCTCTCTGCAATC-3' | Fwd: 5'-CCTCGATCAGTGTTGTGAAGAG-3'<br>Rev: 5'-CTGTGAAGAGGCCAAGGAAA-3' |
| Dyrk3 | 226419 | Total gene expression | Random hexamers | Fwd: 5'-CCCACCCTATTCGGACACATT-3'<br>Rev: 5'-TGAAACAGTTGTTCCACCTTCAT-3' |
| | | Expression of DoG | TES_RT: 5'-CAGGGCTTCCACACTGATAA-3' | Fwd: 5'-CCCACCCTATTCGGACACATT-3'<br>Rev: 5'-TGAAACAGTTGTTCCACCTTCAT-3' |
| Rps19 | 20085 | Total gene expression | Random hexamers | Fwd: 5'-CAGCAGGAGTTCGTCAGAGC-3'<br>Rev: 5'-CACCCATTCGGGGACTTTCA-3' |
| | | Expression of DoG | TES_RT: 5'-CAGTTGGCAGTTCCTGAAGA-3' | Fwd: 5'-CAGCAGGAGTTCGTCAGAGC-3'<br>Rev: 5'-CACCCATTCGGGGACTTTCA-3' |
| Hsp90aa1 | 15519 | Total gene expression | Random hexamers | Fwd: 5'-TGTTGCGGTACTACACATCTGC-3'<br>Rev: 5'-GTCCTTGGTCTCACCTGTGATA-3' |
| | | Expression of DoG | TES_RT: 5'-CTACTTTCCTTCCCTCCCTTTG-3' | Fwd: 5'-TGTTGCGGTACTACACATCTGC-3'<br>Rev: 5'-GTCCTTGGTCTCACCTGTGATA-3' |
| Tbp | 21374 | Internal conrol for total gene expression | Random hexamers | Fwd: 5’-CCCCACAACTCTTCCATTCT-3’<br>Rev: 5'-5’-GCAGGAGTGATAGGGGTCAT-3’ |
| | | Internal conrol for expression of DoG | TES_RT: 5'-GTGGTCTTCCTGAATCCCTTTA-3' | Fwd: 5’-CCCCACAACTCTTCCATTCT-3’<br>Rev: 5'-5’-GCAGGAGTGATAGGGGTCAT-3’ |

**Table S5.** qRT-PCR primers used to quantify TES-TSS region interactions(Figure 7F).

| Name of Gene | NCBI Gene ID | Amplicon length | RT-PCR primers |
|---|---|---|---|
| Tspo | 12257 | 58 bp | 5'-CCTTGGGTTGGTAGTGTGGA-3' |
| | | | 5'-AAGAGAGACAGCCTGTGGAC-3' |
| Ttll12 | 223723 | 90 bp | 5'-CCGTCCGCTTAAATCCTGC-3' |
| | | | 5'-CCTTGGGTTGGTAGTGTGGA-3' |
| Git1 | 216963 | 205 bp | 5'-TCCGAATCAGGGCCACTATC-3' |
| | | | 5'-CAGAGCAGCAGTGACTTGTG-3' |
| Trp53i13 | 216964 | 158 bp | 5'-TCACAGACATCCACTTGCCT-3' |
| | | | 5'-TCCGAATCAGGGCCACTATC-3' |
| Dyrk3 | 226419 | 144 bp | 5'-GCCCAGCCTCGTCTTGTA-3' |
| | | | 5'-TCACAAGCTTTTCCTGTGGC-3' |
| Rps19 | 20085 | 102 bp | 5'-CAAAATGGGCGGGTCTGTAA-3' |
| | | | 5'-ATGCACTTCACACTGGGAGA-3' |
| Hsp90aa1 | 15519 | 104 bp | 5'-TGAGTACACTGTCCCTGTCT-3' |
| | | | 5'-GGGGCATTAAGTAGAAACAGTTT-3' |
| Mlx (Control) | 21428 | 76 bp | 5'-AGTCCGCTGGCTTGTTT-3' |
| | | | 5'-TTGACCCAAGGGTCCTC-3' |
| | | | prob: 56-FAM/CGGTTCGGT/ZEN/AGGTTCACGATGACG/3IAB kFQ/ |

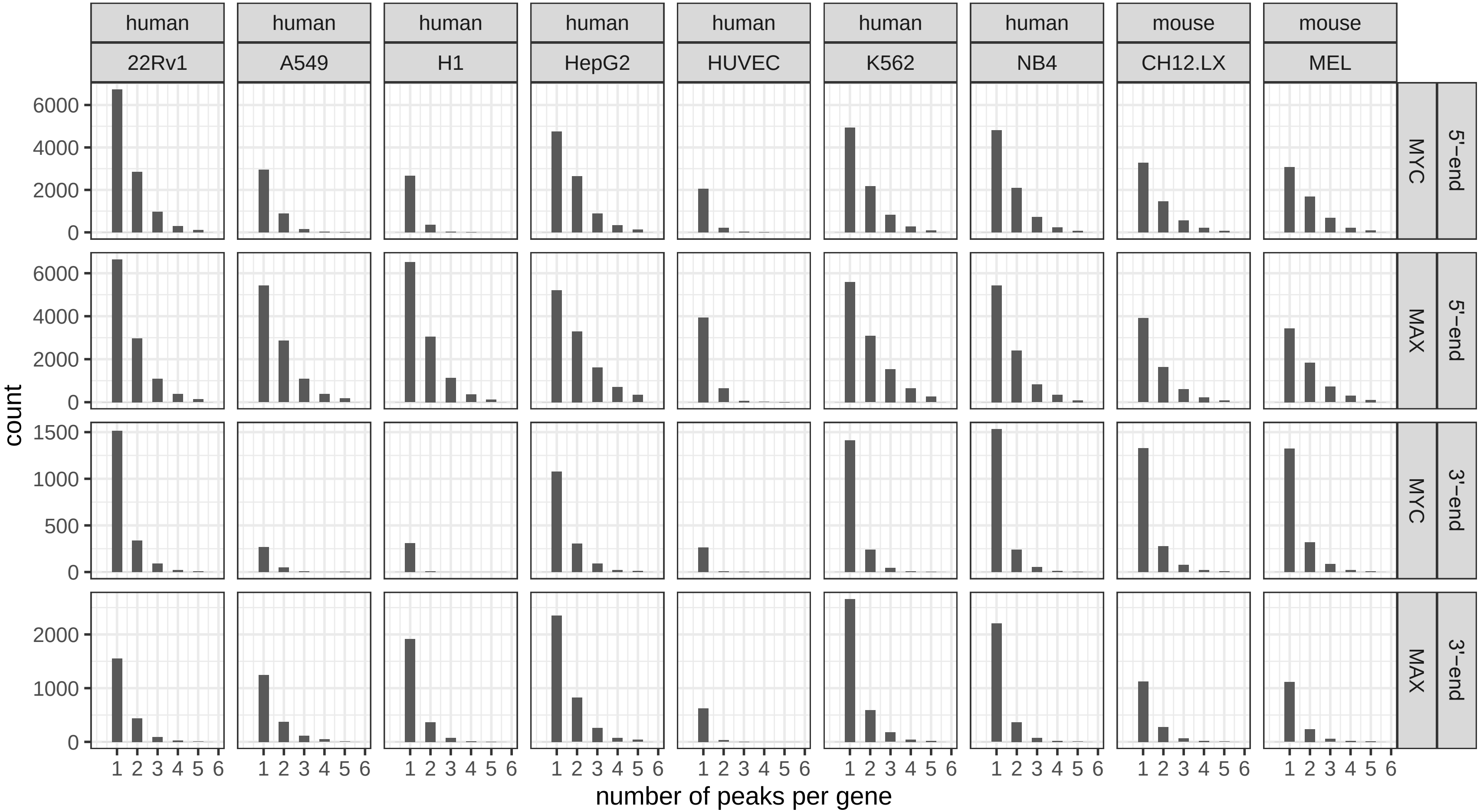


**Figure S1. Distribution of MYC and/or MAX binding sites residing within +/- 2.5 kb of TSSs and TESs.** Results of MYC and MAX binding were obtained from the ENCODE and GEO databases (Table S1). AnnotatePeakInBatch (ChIPpeakAnno Version 3.6.5) was used to assign binding sites.

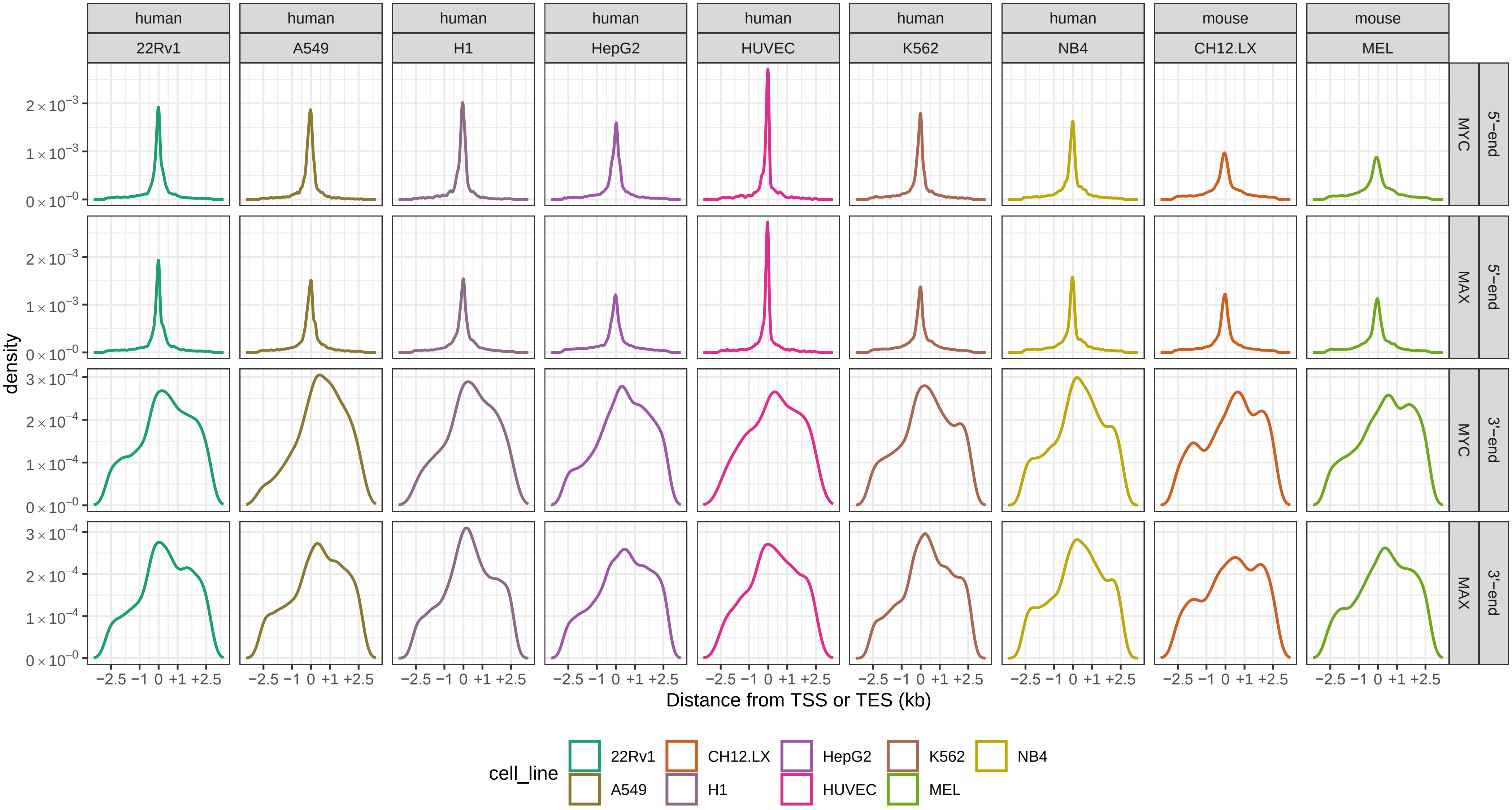


**Figure S2. Binding of MYC and MAX around TSSs and TESs of genes for the individual cell lines shown in** Figure 1B.

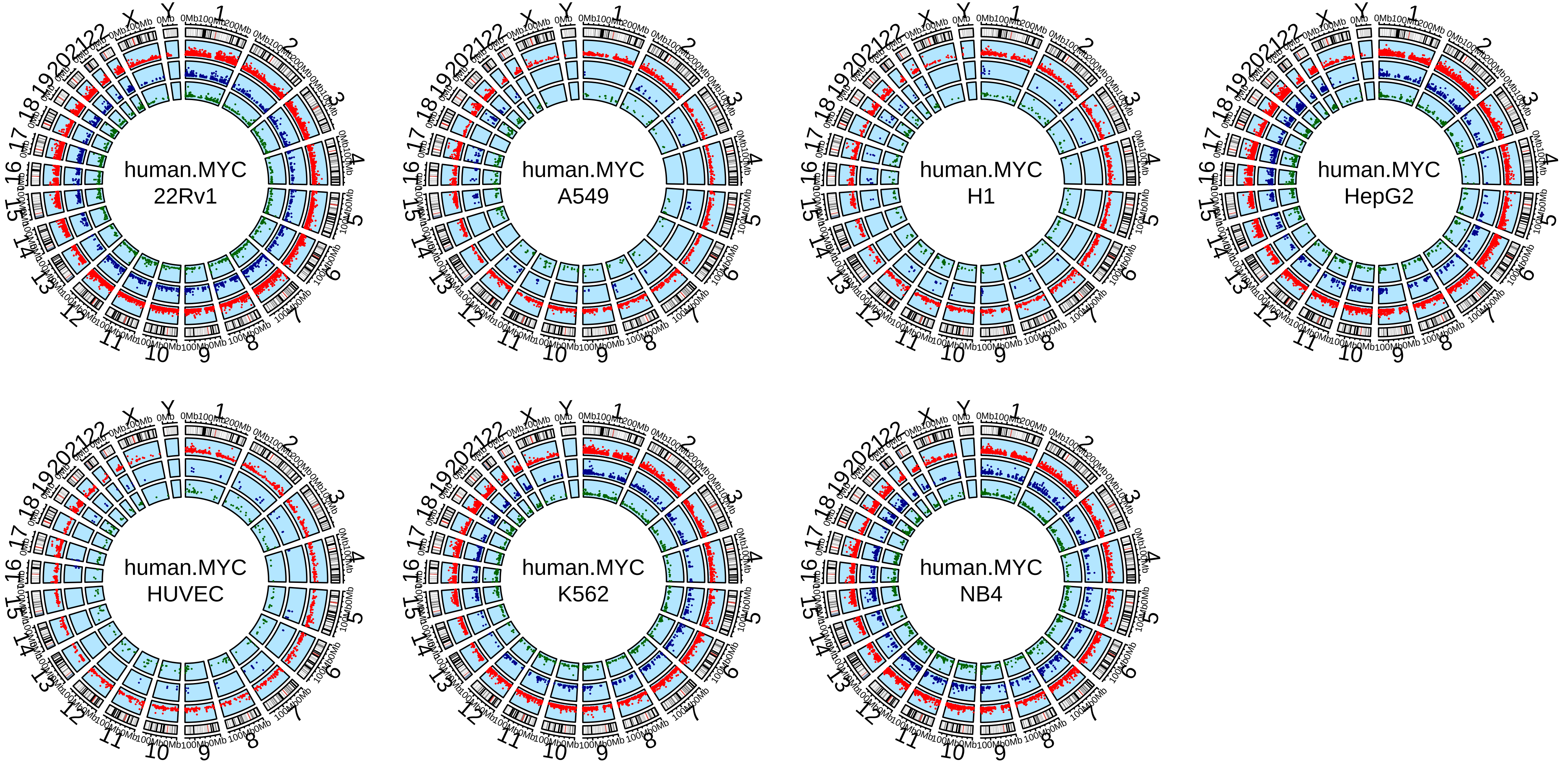


**Figure S3. Circos plots showing the chromosomal locations of human genes that bind MYC only in the vicinity of TSSs and/or TESs in each of the 7 cell lines used in this study.** Red dots: locations of genes associated with MYC binding near TSSs only; green dots: locations of genes associated with MYC binding near TESs only; blue dots: locations of genes associated with MYC binding near both sites.

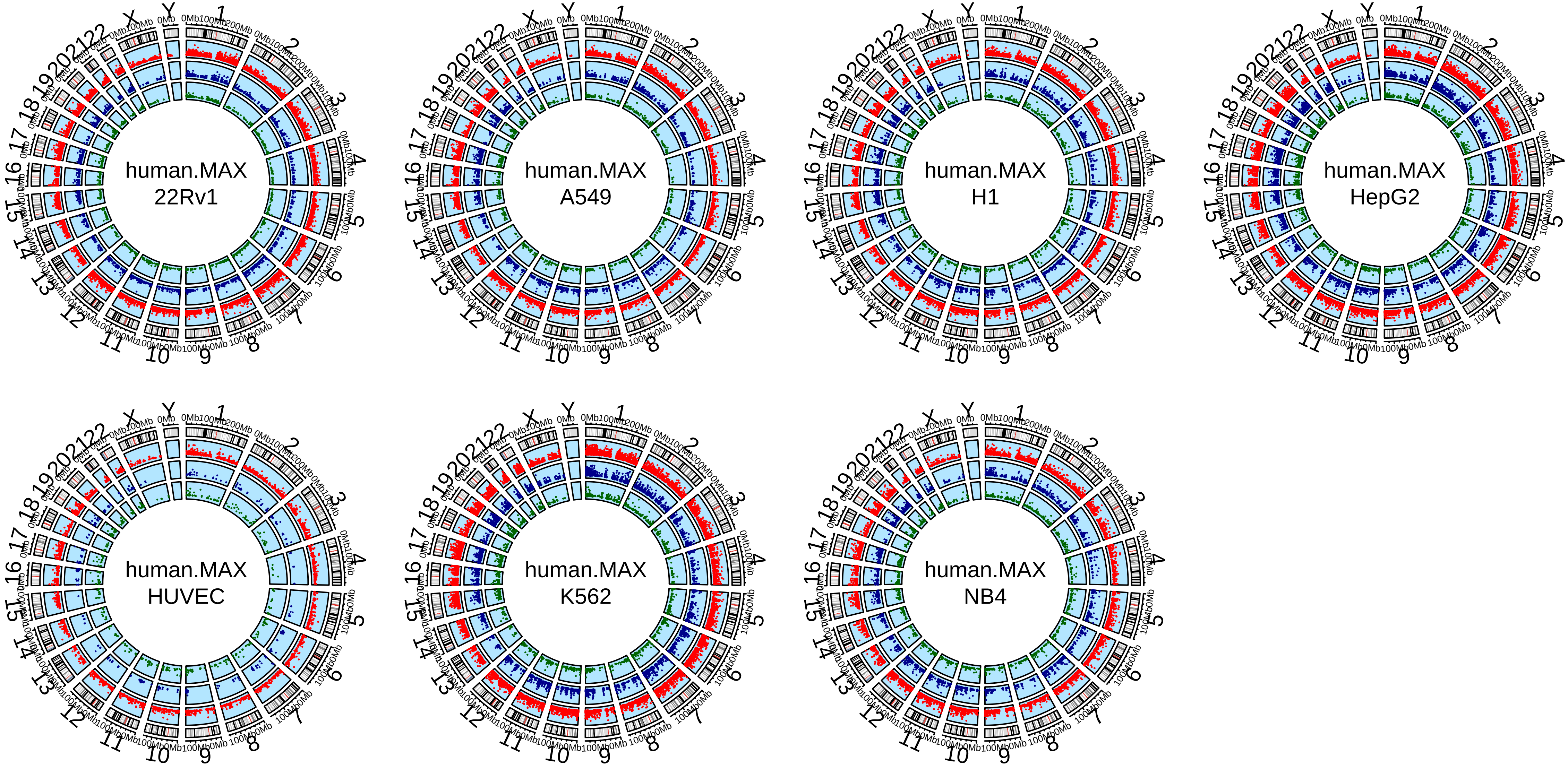


**Figure S4. Circos plots showing the chromosomal locations of human genes that bind MAX only in the vicinity of TSSs and/or TESs in each of the 7 cell lines used in this study.** Red dots: locations of genes associated with MAX binding near TSSs only; green dots: locations of genes associated with MAX binding near TESs only; blue dots: locations of genes associated with MAX binding near both sites.

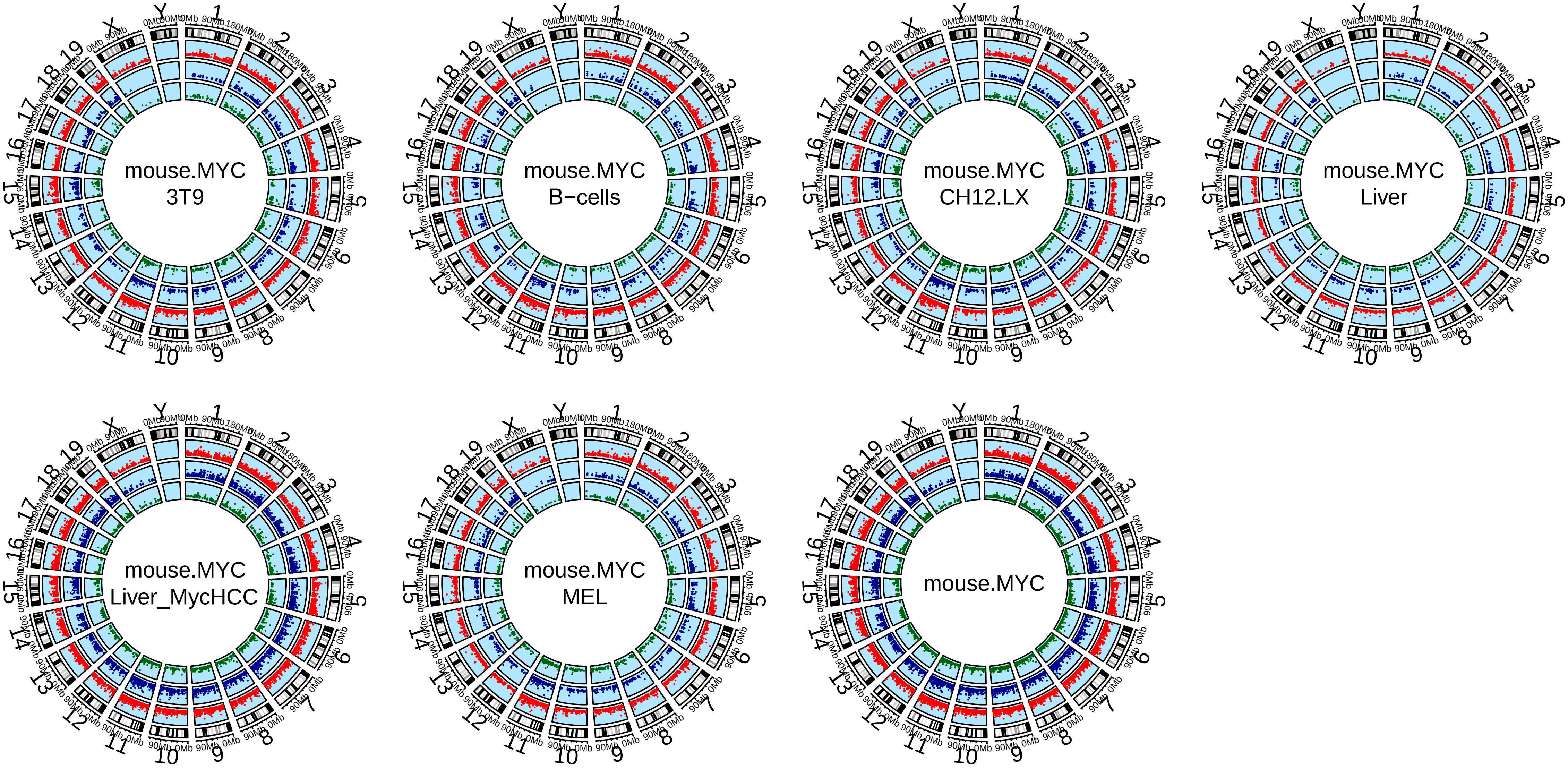


**Figure S5. Circos plots showing the chromosomal locations of mouse genes that bind MYC only in the vicinity of TSSs and/or TESs in each of the tissues and cell lines used in this study.** Red dots: locations of genes associated with MYC binding near TSSs only; green dots: locations of genes associated with MYC binding near TESs only; blue dots: locations of genes associated with MYC binding near both sites. The last diagram shows the combined results similar to that shown for human genes in Figure 1F.

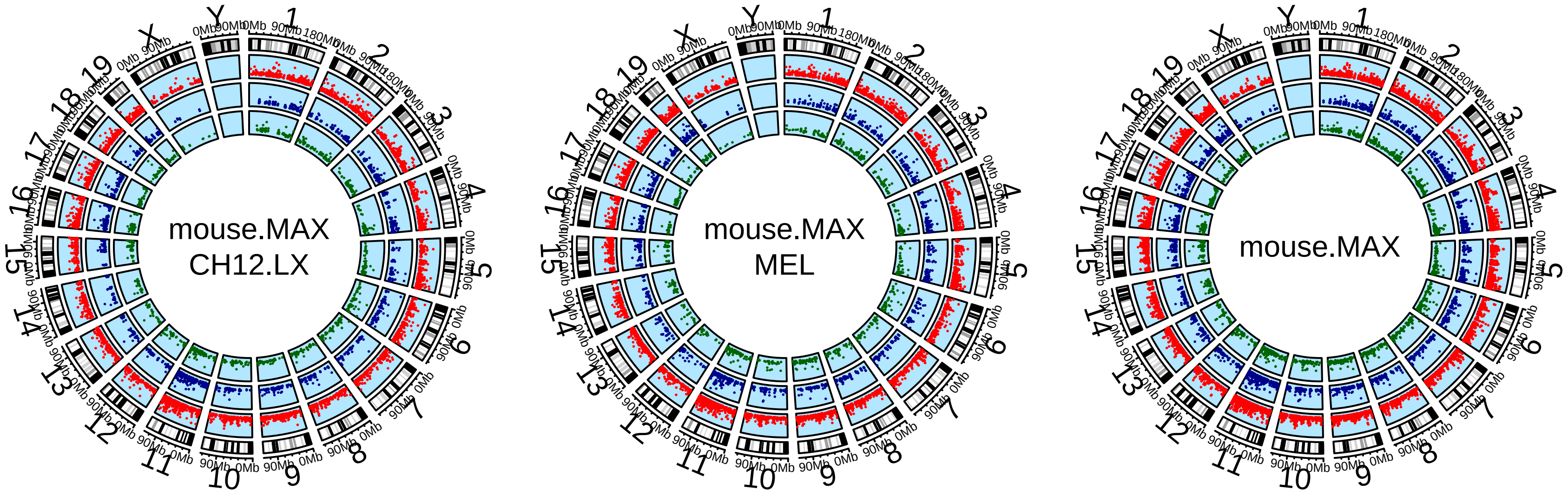


**Figure S6. Circos plots showing the chromosomal locations of mouse genes that bind MAX only in the vicinity of TSSs and/or TESs in each of the cell lines used in this study.** Red dots: locations of genes associated with MAX binding near TSSs only; green dots: locations of genes associated with MAX binding near TESs only; blue dots: locations of genes associated with MAX binding near both sites. The last diagram shows the combined results similar to that shown for human genes in Figure 1F.

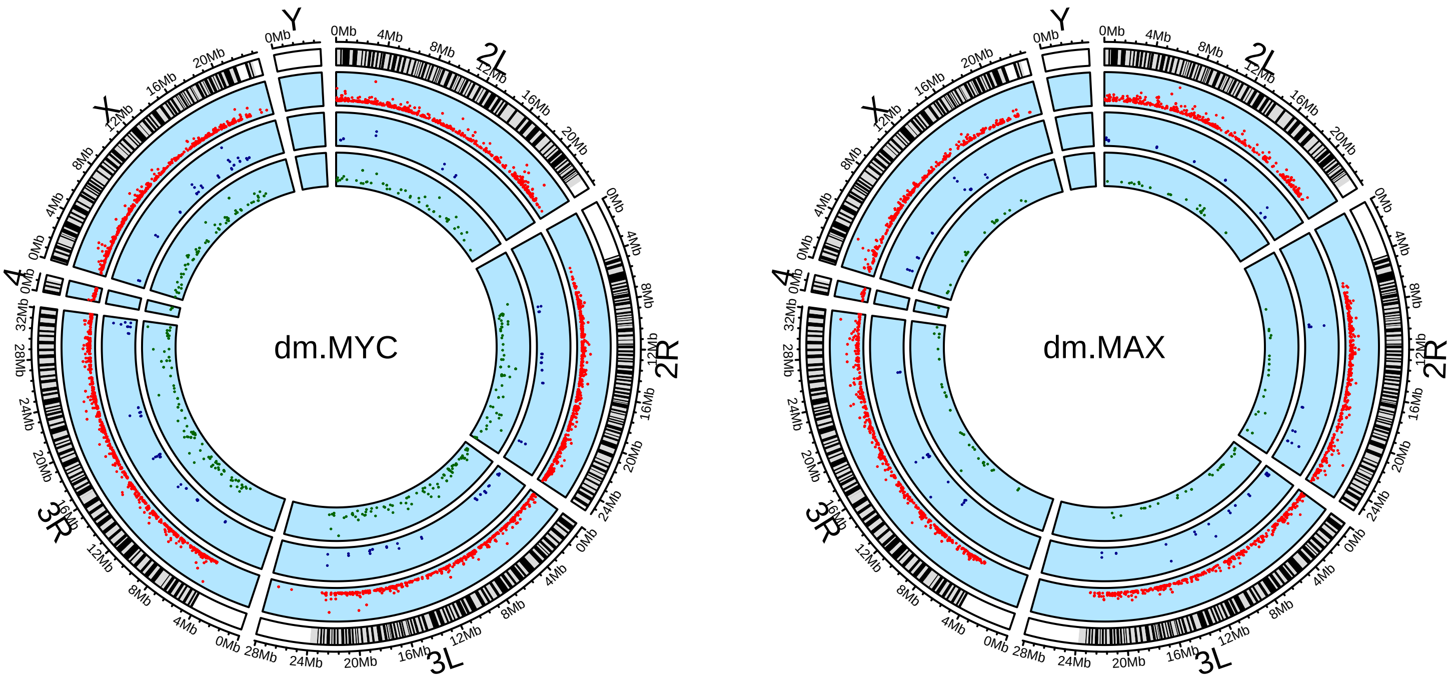


**Figure S7. Circos plots showing the chromosomal locations of *D. melanogaster* genes that bind MAX only in the vicinity of TSSs and/or TESs.** Gene locations are based on ChIP-seq results obtained from third instar larvae.[18] Red dots: locations of genes associated with dMYC or dMAX binding around TSSs only; green dots: locations of genes associated with dMYC or dMAX binding around TESs only; blue dots: locations of genes associated with dMYC or dMAX binding at both TSSs and TESs.

### Figure S8.

#### Git1/Trp53i13_TES_E-box

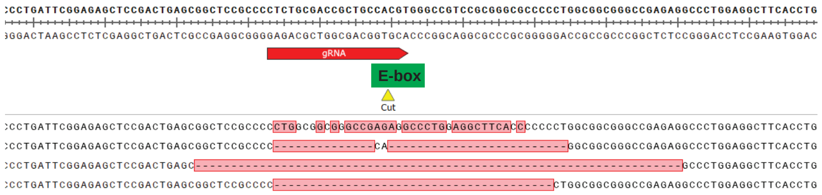


#### Tspo/Ttll12_TES_E-box

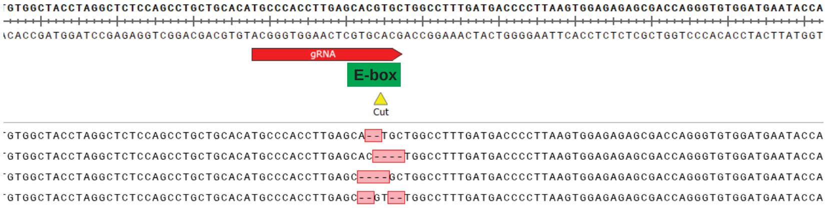


#### Hsp90aa1_TES_E-box

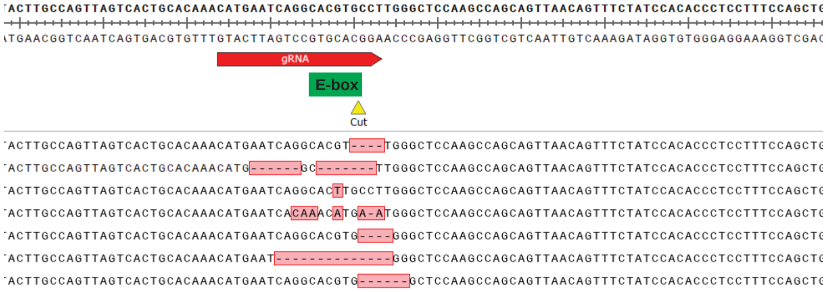


#### Dyrk3_TES_E-box

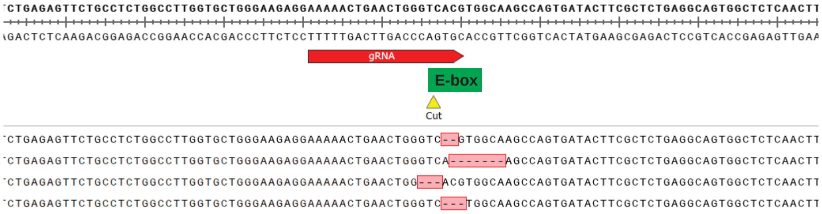


#### Rps19_TES_E-box

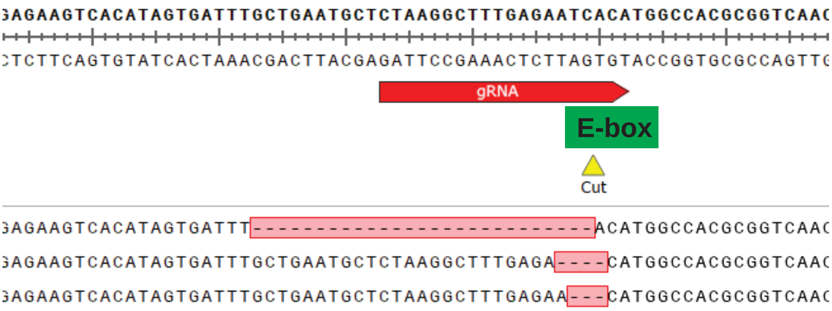

**Figure S8. Mutagenesis of TES-associated E boxes identified as sites of MYC+MAX binding in immortalized murine fibroblasts.**

The top portion of each panel displays the WT gene sequence surrounding the TES-associated E box, and the gRNA used for Crispr-mediated editing. Beneath this are the mutant sequences identified subsequent to Crispr-Cas9 targeting of the sites. In keeping with the fact that NIH3T3 cells are hyperdiploid,[71] all clones contained between 3 and 6 different mutant alleles.

Figure S9.

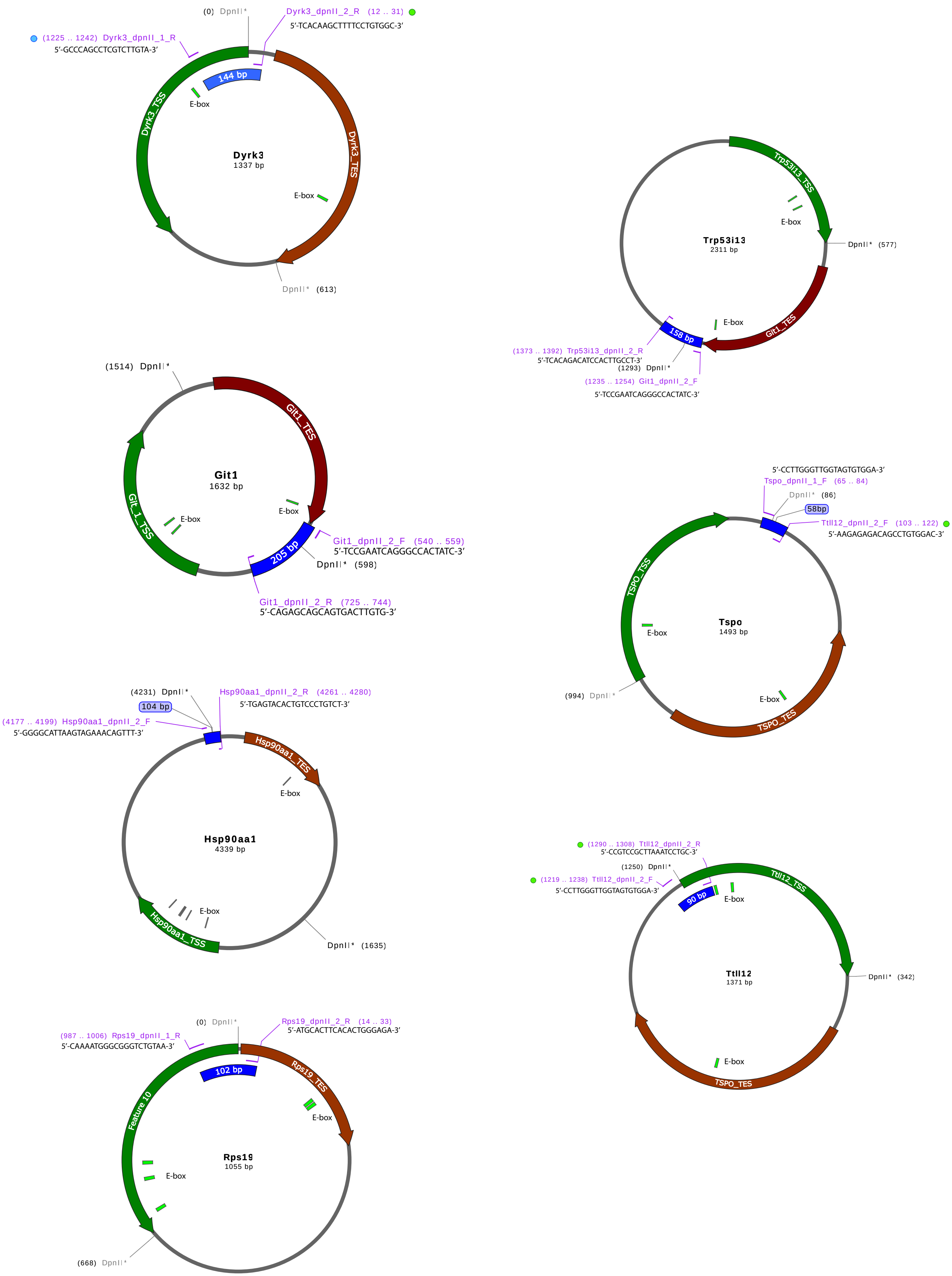

(0) DpnII*
Dyrk3_dpnII_2_R (12 .. 31)
5′-TCACAAGCTTTTCCTGTGGC-3′
(1225 .. 1242) Dyrk3_dpnII_1_R
5′-GCCCAGCCTCGTCTTGTA-3′
144 bp
E-box
Dyrk3_TSS
Dyrk3
1337 bp
Dyrk3_TES
E-box
DpnII* (613)
Trp53i13_TSS
E-box
DpnII* (577)
Trp53i13
2311 bp
E-box
Git1_TES
158 bp
(1373 .. 1392) Trp53i13_dpnII_2_R
5′-TCACAGACATCCACTTGCCT-3′
(1293) DpnII*
(1235 .. 1254) Git1_dpnII_2_F
5′-TCCGAATCAGGGCCACTATC-3′
(1514) DpnII*
Git1_TES
Git1
1632 bp
E-box
E-box
Git_1_TSS
205 bp
Git1_dpnII_2_F (540 .. 559)
5′-TCCGAATCAGGGCCACTATC-3′
DpnII* (598)
Git1_dpnII_2_R (725 .. 744)
5′-CAGAGCAGCAGTGACTTGTG-3′
5′-CCTTGGGTTGGTAGTGTGGA-3′
Tspo_dpnII_1_F (65 .. 84)
DpnII* (86)
58bp
Ttll12_dpnII_2_F (103 .. 122)
5′-AAGAGAGACAGCCTGTGGAC-3′
TSPO_TSS
E-box
Tspo
1493 bp
(994) DpnII*
E-box
TSPO_TES
(4231) DpnII*
Hsp90aa1_dpnII_2_R (4261 .. 4280)
5′-TGAGTACACTGTCCCTGTCT-3′
104 bp
(4177 .. 4199) Hsp90aa1_dpnII_2_F
5′-GGGGCATTAAGTAGAAACAGTTT-3′
Hsp90aa1_TES
E-box
Hsp90aa1
4339 bp
E-box
Hsp90aa1_TSS
DpnII* (1635)
(1290 .. 1308) Ttll12_dpnII_2_R
5′-CCGTCCGCTTAAATCCTGC-3′
(1250) DpnII*
(1219 .. 1238) Ttll12_dpnII_2_F
5′-CCTTGGGTTGGTAGTGTGGA-3′
90 bp
E-box
Ttll12_TSS
Ttll12
1371 bp
DpnII* (342)
E-box
TSPO_TES
(0) DpnII*
Rps19_dpnII_2_R (14 .. 33)
5′-ATGCACTTCACACTGGGAGA-3′
(987 .. 1006) Rps19_dpnII_1_R
5′-CAAAATGGGCGGGTCTGTAA-3′
102 bp
Rps19_TES
E-box
Feature 10
Rps19
1055 bp
E-box
(668) DpnII*

**Figure S9. Strategy for quantifying TSS-TES E box contacts in murine fibroblasts and list of qPCR primers used for this purpose.**

Each circle shows the expected structure and size (in bps) of loops formed between TSS- and TES-proximal regions (green and brown thick lines, respectively) following DpnII digestion of formaldehyde cross-linked genomic DNA and re-ligation. Locations of ligated DpnII sites and E boxes (or their previous locations) are indicated. Sequences of PCR primers used to amplify ligated regions flanking the sites of DpnII digestion/re-ligation are indicated as are the predicted sizes of the amplified DNA products, the primers are also listed in Table S5.

**File S1.** Rank of 1210 TFs and transcriptional co-factors and the frequency with which they bound to MYC-associated TSSs, TESs and intragenic and distal enhancer elements. The ReMap2022 database (https://remap2022.univ-amu.fr/) was used a source of binding profile.
